# Discovering Genetic Modulators of the Protein Homeostasis System through Multilevel Analysis

**DOI:** 10.1101/2024.02.26.582154

**Authors:** Vishal Sarsani, Berent Aldikacti, Tingting Zhao, Shai He, Peter Chien, Patrick Flaherty

## Abstract

Every protein progresses through a natural lifecycle from birth to maturation to death; this process is coordinated by the protein homeostasis system. Environmental or physiological conditions trigger pathways that maintain the homeostasis of the proteome. An open question is how these pathways are modulated to respond to the many stresses that an organism encounters during its lifetime. To address this question, we tested how the fitness landscape changes in response to environmental and genetic perturbations using directed and massively parallel transposon mutagenesis in *Caulobacter crescentus*. We developed a general computational pipeline for the analysis of gene-by-environment interactions in transposon mutagenesis experiments. This pipeline uses a combination of general linear models (GLMs), statistical knockoffs, and a nonparametric Bayesian statistical model to identify essential genetic network components that are shared across environmental perturbations. This analysis allows us to quantify the similarity of proteotoxic environmental perturbations from the perspective of the fitness landscape. We find that essential genes vary more by genetic background than by environmental conditions, with limited overlap among mutant strains targeting different facets of the protein homeostasis system. We also identified 146 unique fitness determinants across different strains, with 19 genes common to at least two strains, showing varying resilience to proteotoxic stresses. Experiments exposing cells to a combination of genetic perturbations and dual environmental stressors show that perturbations that are quantitatively dissimilar from the perspective of the fitness landscape are likely to have a synergistic effect on the growth defect.

**Significance Statement:** This study provides critical insights into how cells adapt to environmental and genetic challenges affecting protein homeostasis. Using multilevel statistical analysis and transposon mutagenesis, we find that a model organism, *Caulobacter crescentus*, lacks a universal redundancy mechanism for coping with stress, as evidenced by the limited overlap in essential genes across different environmental and genetic perturbations. Our methods also pinpoint key fitness determinants and enable the prediction of perturbation combinations that synergistically affect cell growth.

---

Protein homeostasis is the maintenance of the balance of protein synthesis, protein folding, trafficking, and degradation within a cell. The protein quality control system primarily contains a collection of chaperones and proteases that maintain the homeostatic balance of folding and degradation. Changes in environment, age, or stress can cause imbalances in the healthy proteome. Dysfunction in proteome homeostasis impacts the onset of various metabolic, oncological, cardiovascular, and neurodegenerative diseases (1–3). Understanding the components and pathways in dysregulated proteostasis is critical for developing novel drug development strategies. The proteomes in bacteria are much smaller and less complex than those of humans. Still, most proteostasis network components, like chaperones and proteases, are conserved during billion years of evolution (4). Notably, research on Caulobacter crescentus underscores the dynamic roles of these networks in regulating both the cell cycle and stress responses (5).

Large-scale genome-wide screening can link genes to phenotypes on a comprehensive level. The recent decade has seen the advent of several high-throughput technologies for gene disruption and interaction discovery in microorganisms, enabling the functional annotation of microbial genomes and discovering intricate biological pathways. These approaches include CRISPR-based methods for gene knockdowns (6) and transposon-insertion sequencing (TIS), which was initially proposed as a highly reliable and sensitive technique for detecting modifications in mutant fitness with adequate density across all regions in a genome (7). Random barcode transposon-site sequencing (RB-Tn-Seq) overcomes the cost and scale of the multistep library preparations in the traditional TIS experiments (8) by faster screening via one-step PCR barcode amplification and tracking of mutant frequencies. Despite the advances, identifying essential genes using TIS is still challenging due to variations in experimental parameters such as the transposon used, experimental conditions, and library complexity (9, 10). Studying shared patterns of essentiality across environments or understanding the conserved patterns of essential genes across multiple conditions is critical for understanding complex systems like protein homeostasis.

In this work, we propose a systematic multilevel analysis approach to dissect the genetic modulators of protein homeostasis in *Caulobacter crescentus*. Our primary objective is to investigate how the fitness landscape changes in response to environmental and genetic perturbations by combining proteotoxic stresses and functional inactivation of protein homeostasis genes using massively parallel transposon mutagenesis in *Caulobacter crescentus*. Sequencing is utilized to quantify the frequency of transposon-induced mutations and identify a set of conditionally essential, beneficial, or detrimental genes for each environment by applying a regularized negative binomial regression combined with local False Discovery Rate (FDR) testing within a general linear model (GLM) framework. While determining the overall fitness contribution under selective depletion or stress can be achieved through the number of conditionally essential, beneficial, or detrimental genes, assessing the marginal contribution of a specific gene to overall fitness remains challenging. To address this challenge, we employ the statistical knockoffs methodology (11, 12) to identify important fitness determinants while controlling for the overall false discovery rate. Finally, we apply a nonparametric Bayesian model (13) to understand the associations among a strain’s most predictive fitness determinants. The utility of our analysis is highlighted by experiments that reveal strain-specific interactions between proteotoxic stresses, using growth curves to probe the adaptability of the protein homeostasis network.

## Results

### Genome-wide analysis of conditional essentiality

We focused on proteotoxic stresses and those genes responsible for maintaining protein homeostasis as major players in this stress response are well characterized. Heat stress causes general protein misfolding and thermal denaturation (14), hydrogen peroxide induced oxidative stress modifies ligands and proteins to induce protein misfolding (15), and as an uncharged analog of arginine, canavanine causes protein misfolding upon incorporation into translated polypeptides (16). Proteases responsible for degradation of misfolded proteins (17, 18) and unfoldases that rescue aggregated proteins (ClpB (19) and ClpA) were targeted for deletion in this current study. Chaperones play a crucial role in folding proteins en route to the native state and are upregulated upon proteotoxic stress. Because the Hsp70 chaperone DnaK is essential in Caulobacter (20), we took advantage of a non-stress inducible (dnaK-NI) variant to generate sufficient DnaK protein for viability, but this construct is incapable of normal stress induced upregulation.

Our genome-wide profiling reveals higher median unique insertion counts across all genes in wild-type and Δ*lon* strains compared to Δ*clpA*, Δ*clpB*, and dnaKJ-NI (SI Appendix, Fig. S2-S3, Table. S2). To analyze gene dependency in the system, we assess the proportion of essential genes under varying stress conditions within different strains. Figure 2A shows a tabulation of the counts of genes that are conditionally essential, beneficial, or detrimental for each gene-by-environment condition. These counts, adjusted relative to each strain’s genetic background, isolate the effects of environmental perturbations and align with the generalized linear model structure employed in our analysis. The dnaKJ-NI strain exhibits a higher average number of conditionally essential genes across all environmental perturbations compared to all other strains. In contrast, the wild-type strain shows the lowest average number of such genes. This suggests that the dnaKJ-NI strain may be more sensitive to environmental changes, requiring a greater number of essential genes for survival, while the wild-type strain appears to be more robust, relying on fewer essential genes. The combination of Δlon and high oxidative stress led to the most significant changes in the count of conditionally essential genes, highlighting the heightened sensitivity of the protein homeostasis system in the Δlon background to oxidative stress. We also observed that a gene may be conditionally beneficial under a particular condition but may change its essentiality under a different proteotoxic stress or stress level (SI Appendix, Fig. S4-S10). In Figure 2B, we assess the degree of overlap in essential genes between various gene-by-environment conditions. Interestingly, within each strain background, environmental perturbations show a high degree of overlap (SI Appendix, Fig. S10-S12). This suggests that genetic background has a stronger influence on the essential gene profile than the environmental conditions themselves. Notably, the highest degree of overlap was observed between the Δ*clpA* and wild-type strains, while the other strains exhibited minimal overlap. Recall that our genetic perturbations were designed to target different facets of the protein homeostasis system (see Figure 1). Therefore, these results suggest the involvement of a unique set of proteins specific to different aspects of the protein homeostasis system.

**Fig. 1.**
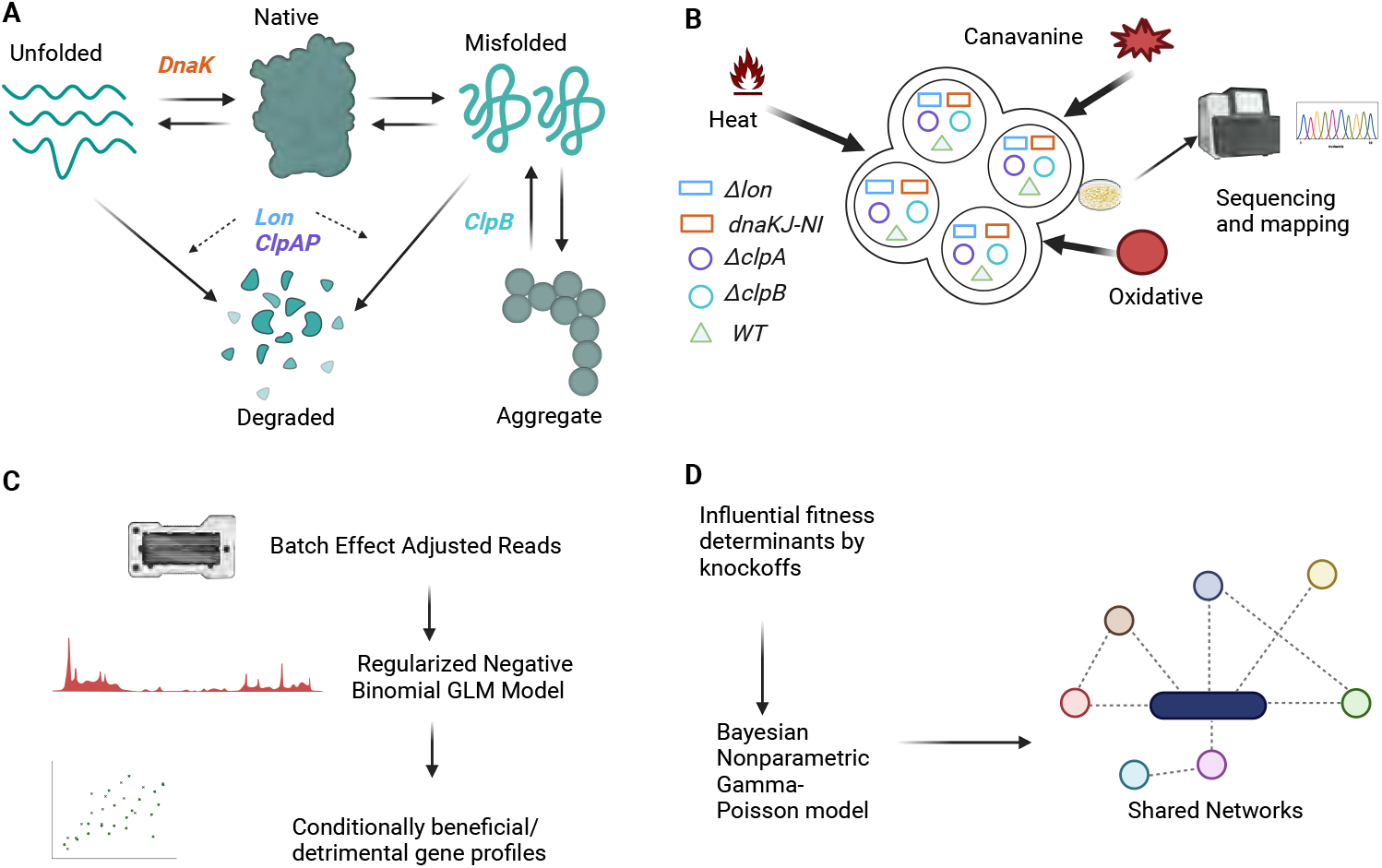
A schematic pipeline for identifying genetic modulators of protein homeostasis system in *Caulobacter crescentus*. **A**. A schematic representation of the *Caulobacter crescentus* proteostasis network and some key regulators. DnaK assists proteins in folding into their functional native state. Lon and ClpAP degrade and eliminate the unfolded and misfolded proteins. ClpB mediates the disaggregation of misfolded and aggregated proteins. **B**. Transposon insertion sequencing is used to investigate the gene fitness landscape changes in response to proteotoxic stresses in the context of disruptions of protein homeostasis system components. Transposon libraries are constructed in wild-type *Caulobacter crescentus* and strains deficient in specific chaperone or protease genes responsible for protein homeostasis. These libraries were subjected to three different proteotoxic stresses (Canavanine, Heat, and Oxidative) at three different levels. **C**. The transposon insertion count data is corrected for batch effects, and a regularized negative binomial GLM model is fit. Significant changes in insertion counts due to changes in stress conditions are identified with local false discovery rate control to identify conditionally beneficial and detrimental genes. **D**. Genes that are important for discriminating between proteotoxic stresses in each background strain are identified by a model-Y knockoffs procedure (12). A Bayesian nonparametric Gamma-Poisson model is used to identify commonalities and differences in the network of genes that are important across stresses.

**Fig. 2.**
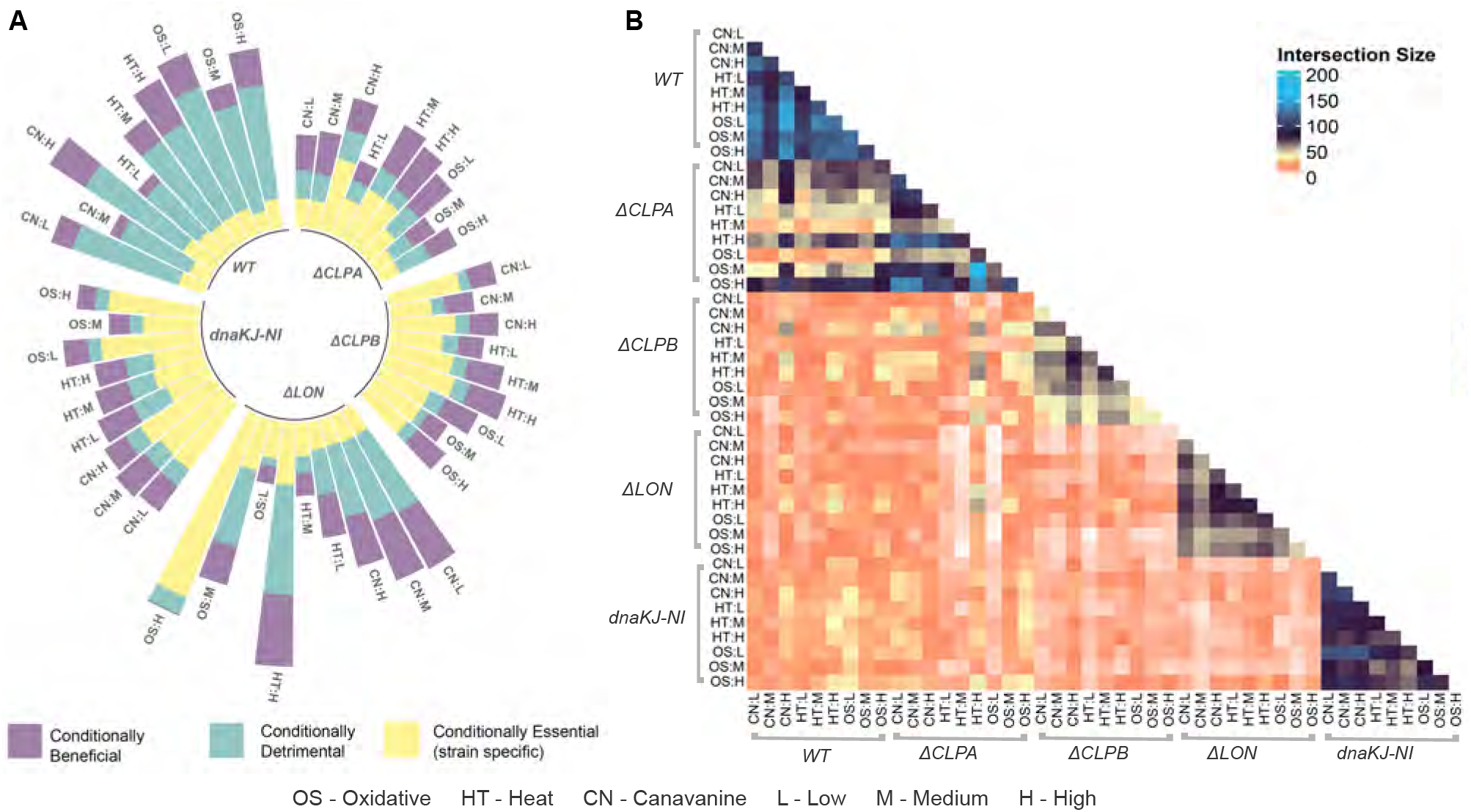
Genome-wide essentiality profiling in *Caulobacter crescentus*. **A**. Each portion inside a single bar represents the number of conditionally beneficial, detrimental, and essential components across various protein homeostasis components subjected to proteotoxic stresses of different levels. The Δ*lon* strain has a large number of conditionally essential genes in high oxidative stress compared to high canavanine, indicating that the homeostasis system is significantly sensitized to that proteotoxic stress. **B**. The pair-wise overlap of essentiality profiles between stress conditions. A larger overlap of essentiality profiles is seen in wild type and dnaKJ-NI compared to strains deficient in ClpA, Lon, or ClpB.

### Identification of perturbation predictors in the protein homeostasis system

Using the Model-Y knockoff framework (12), we identified sets of perturbation predictors for each strain: 20 genes in wild-type, 33 in Δ*lon*, 39 in Δ*clpA*, 44 in Δ*clpB*, and 38 in dnaKJ-NI, totaling 146 unique genes (Figure 3, SI Appendix, Fig. S13-S17, Table S3-4). Of these, 19 genes (excluding CCNA_00375) were common across at least two strains. Among the genes without prior functional characterization, the predicted acetyltransferase CCNA_02154 was found to be a consistent predictor across all strains. Its specific sensitivity to canavanine stress, without substantial impact on heat or oxidative stresses, suggests a role for this enzyme in specifically blocking the toxic effect of canavanine, likely by modifying this unnatural amino acid (SI Appendix, Fig. S18). Similarly, CCNA_03861 was identified as a significant gene in Δ*clpA*,

**Fig. 3.**
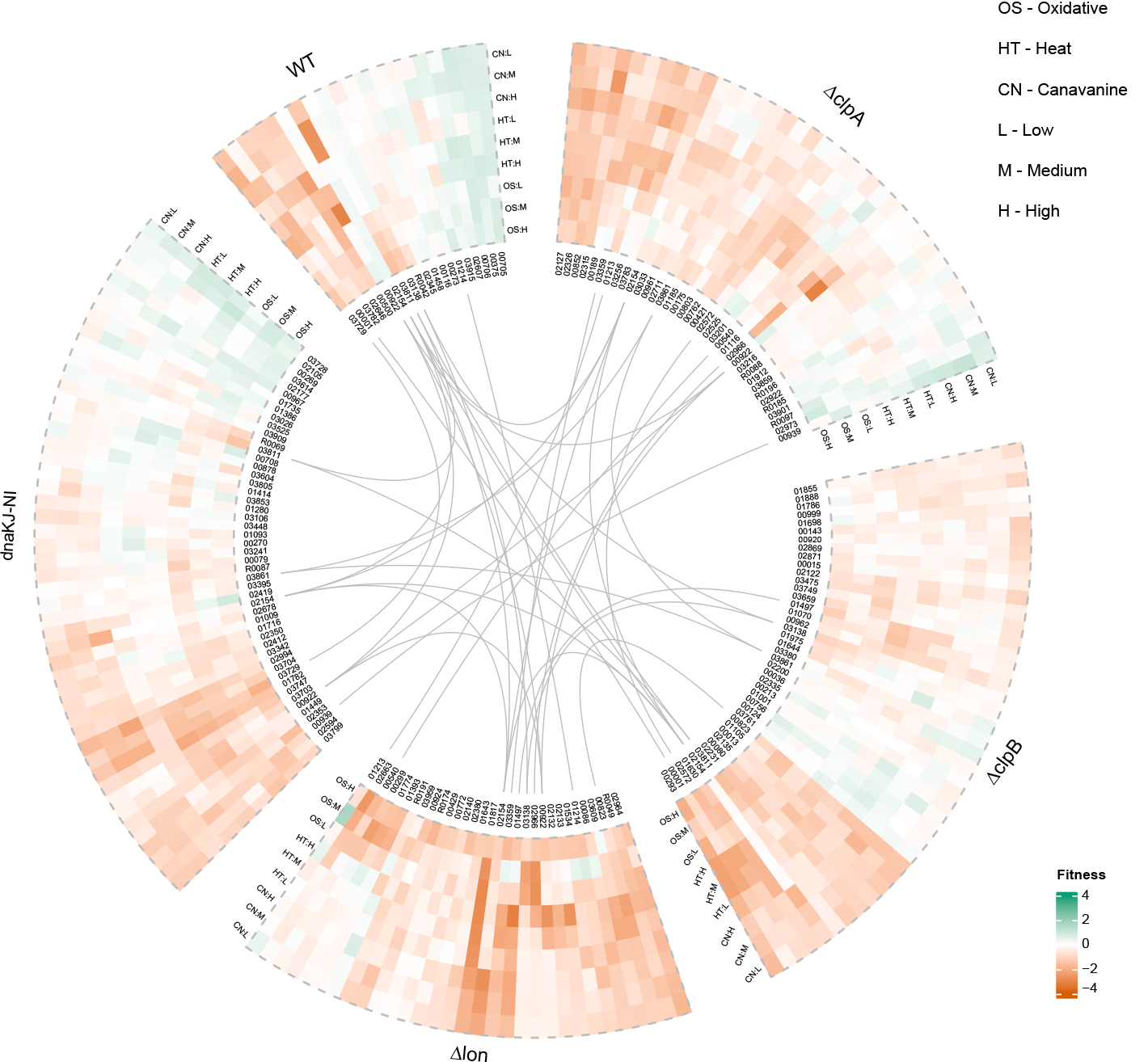
Proteotoxic stress predictors. The model-Y knockoff framework is used to identify predictors of proteotoxic stress within each genetic background. The plot shows genes that appear as predictors of proteotoxic stress in more than one genetic background and predictors that are unique to each genetic background.

Δ*clpB*, and dnaKJ-NI, potentially involved in pyridoxal phosphate homeostasis. Considering genes with known functions, as expected, ClpB (CCNA_00922) was identified as a perturbation predictor in multiple strains. Additionally, the catalase KatG (CCNA_03138), which plays a role in the hydrogen peroxide detoxification process, and OxyR (CCNA_03811), a transcription factor known to be important for the oxidative stress response, were also identified as key perturbation predictor in several strains. Across strains, the fitness values of these predictors remained consistent under proteotoxic stresses, including Heat, Oxidative, and Canavanine. However, differences were observed based on the stress severity. When homeostasis components were deleted, these predictors demonstrated resilience to stress conditions, with some still showing sensitivity to stress levels.

### Predicting *in vivo* growth in combinatorial perturbations from single perturbation transposon insertion sequencing data

Given transposon insertion sequencing data from a single environmental perturbation, we asked whether it is possible to predict the effect of multiple perturbations on growth rate. Double perturbations have the potential to overwhelm the compensatory mechanisms in the protein homeostasis system. Our hypothesis was that perturbations that have highly differentiated conditionally essential profiles will yield a synergistic effect on the inhibition of growth.

To quantify the distance between two conditionally essential profiles, we used Earth Mover’s Distance (EMD) between both total and unique insertion count distributions of genes identified by the GLM framework. In the Δ*clpB* strain, the EMD between heat and oxidative stress levels for both total and unique counts are notably consistent and low (Figures 4(A-B), SI Appendix, Fig. S19, Table. S5). This suggests that a combination of heat and oxidative stress will have a limited or perhaps additive effect on growth in cells lacking the ClpB disaggregase. In contrast, the EMD between heat and high oxidative stress in the Δ*clpA* is high in both total and unique count data, suggesting that the effect of the combination of the stresses on growth is synergistic if the hypothesis is true. Likewise, the EMD between heat and high oxidative stress in the wild-type background is high in unique count data but moderate in total count data. We sought to validate these predictions using individual growth curve measurements in gene-by-environment perturbations with double environmental stress conditions.

**Fig. 4.**
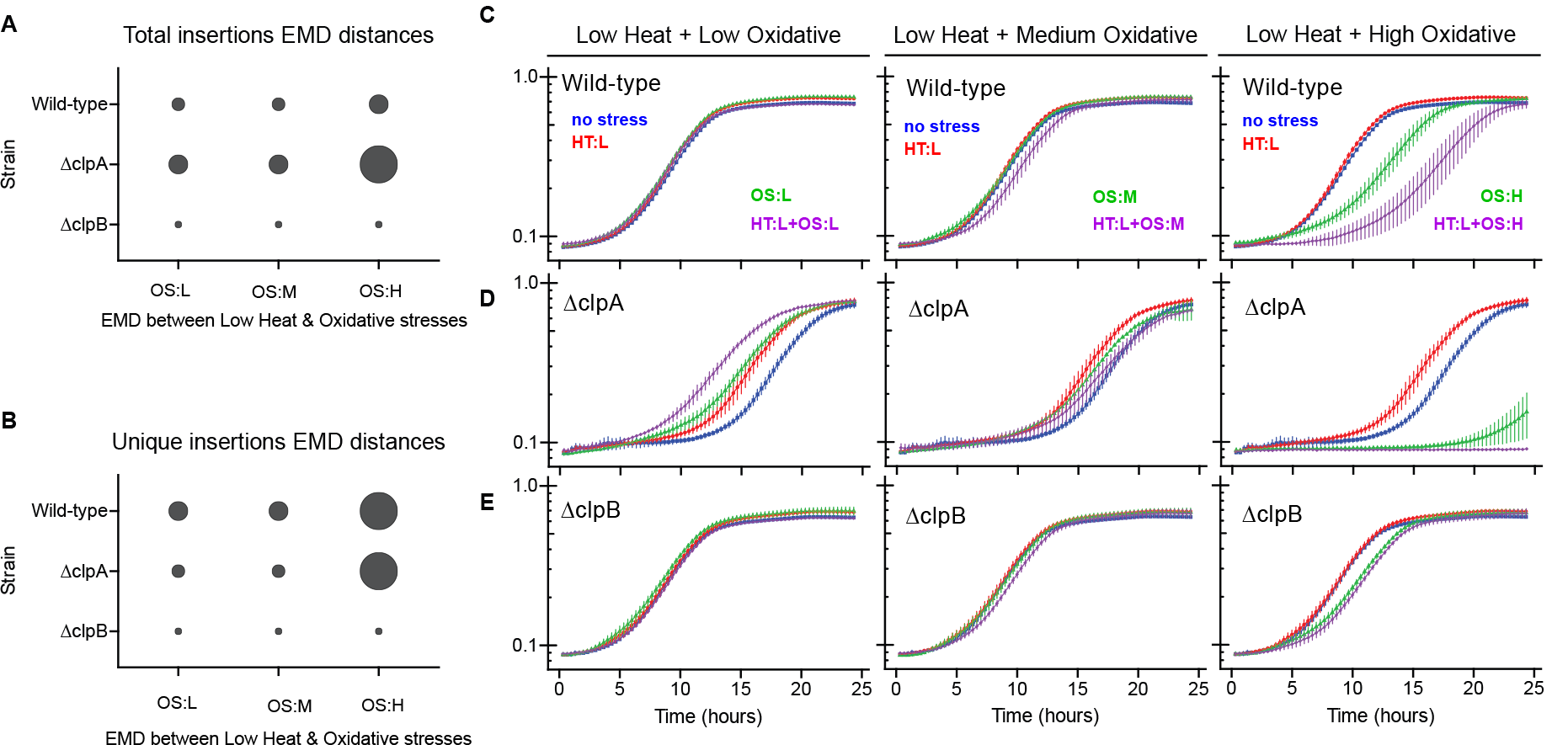
In-vivo Validation of Stress-Induced Fitness Effects. Using tnseq data, we assess the fitness variations under low heat and oxidative peroxide stress in WT, Δ*clpA*, and Δ*clpB* strains. Differences are quantified using EMD (Earth Mover’s Distance) based on both total (**A**) and unique(**B**) insertion counts. Bubble size corresponds to the magnitude of the EMD distances. The OD600 growth curves were generated to approximate the cell count of WT (**C**), Δ*clpA* (**D**), and Δ*clpB* (**E**) subjected to dilutions of low, medium, or high concentrations of hydrogen peroxide (0.025mM, 0.05mM, 0.1mM) after the heat shock treatment.

We subjected wild-type, Δ*clpA*, and Δ*clpB* strains to heat shock and subsequent exposures to varying hydrogen peroxide concentrations using optical density (OD 600) to measure cell density during a 24-hour growth period. As illustrated in Figures 4(C-E), the wild-type strain tolerates low heat and moderate oxidative stress well on its own, but combining low heat with high oxidative stress results in a substantial fitness defect compared to these stresses in isolation. Similarly, the Δ*clpA* strain shows synergistic declines in growth with low heat and high oxidative stress, although, for this strain, it is even more striking as individual low heat stress treatment improves growth, as we reported previously (21). By contrast, the Δ*clpB* strain consistently demonstrates a lack of synergistic growth defects when combined with heat and oxidative stress, supporting the predictions drawn from the analysis of the single-perturbation TIS data.

### Conditionally essential components shared among the proteotoxic stresses

We hypothesized that some clusters of conditionally essential genes could be shared across proteotoxic stress conditions within each genetic background. These co-essential clusters may lead to insights into the underlying structure of the protein homeostasis system. To identify these clusters, we fit a nonparametric Bayesian model based on a Gamma-Poisson model analogous to topic models like Latent Dirichlet Allocation (22, 23). The posterior distribution of the latent variables *H* and *θ*_*ijk*_ captures the clusters of essential genes and their relevance in each stress condition, respectively. In the wild-type (WT) strain, our findings show that the heat stress perturbations are characterized by the essentiality of CCNA_00922 and CCNA_00001, and the oxidative stress perturbations are characterized by the essentiality of CCNA_03811, CCNA_03138, CCNA_02646, and CCNA_00375 (SI Appendix, Fig. S20). On the other hand, CCNA_00708 is essential to all three stress conditions for the *dnaKJ − NI* strain, but CCNA_03811, CCNA_00293, and CCNA_00292 are essential only in oxidative stress conditions (SI Appendix, Fig. S21). Results for the remaining genetic perturbations are shown in SI Appendix, Fig. S22-24. This model-based representation of the TIS data enables a more thorough investigation of the overall changes in the pattern of essential genes induced by different stress conditions.

## Discussion

Understanding how bacteria handle stress is critical for developing novel antibacterial therapeutics and for understanding the fundamental mechanisms of robust and evolutionarily conserved systems. Our study examines the determinants of growth under combinations of genetic and environmental perturbations to the protein homeostasis system to better understand synergistic interactions in the system. A genome-wide analysis of perturbation growth data revealed a low amount of overlap among sets of essential genes across mutant strains with functional deletions targeting diverse aspects of the protein homeostasis system. In contrast, there is a high amount of overlap among sets of essential genes across environmental perturbations within each genetic background. A statistical knockoff strategy revealed important fitness determinants within each deletion strain. The earth-mover distance between sets of conditionally essential genes for single environmental perturbations was predictive of growth defect under combinations of environmental perturbations. Finally, a nonparametric hierarchical Bayesian model enabled the representation of a large amount of TIS data into clusters, or networks, of conditionally essential genes and the attribution of each stress response to a combination of those networks.

### Supporting Information Appendix (SI)

The appendix is available online.

## Materials and Methods

Figure 1 offers an overview of both the experimental and computational approaches employed to investigate the protein homeostasis system in *Caulobacter crescentus*.

### Experimental methods

A schematic representation of experimental data is shown in the SI Appendix, Fig. S1. Transposon mutagenesis libraries used in this study were generated as previously described (24). Briefly, *E. coli* cells containing randomly barcoded Tn5 plasmids (APA766, gift from Deutschbauer lab) are conjugated with wild-type (wt), Δ*lon*, Δ*clpA*, Δ*lon*, Δ*clpB*, and dnaKJ-NI (a non heat-inducible allele of dnaKJ) *Caulobacter crescentus* cells separately. E.*coli* donors are kanamycin-resistant and diaminopimelate (DAP) auxotrophs, requiring it to grow in the media. For conjugation, *E. coli* donor cells and *Caulobacter* strains were mixed at a 1:10 ratio overnight on a PYE agar plate supplemented with DAP (300 *μ*M). The next day, the conjugate was scraped, resuspended, and spread over 14 large (150 x 15 mm) PYE agar plates supplemented with kanamycin (25*μ*g/ml) without DAP per strain. In this culture, the donor cells will not survive due to no DAP, and acceptor *Caulobacter* cells will be selected for the Tn5 plasmid due to kanamycin selection. After 5 days of growth, the colonies were scraped, pooled, and frozen in PYE + 10% glycerol in 1 ml aliquots. For stress condition experiments, 1 aliquot per replicate per strain was thawed in 3.5 ml of PYE or PYE+0.2% xylose and recovered overnight in a 30°C shaker. For all dnaKJ-NI experiments, cells were recovered at saturating xylose concentrations (PYE+ 0.2% xylose), and the stress experiments were done at minimal xylose concentrations. (PYE+0.002%) All conditions were performed in quadruplicates, and optical density (OD) measurements were taken at 600nm. Experiments were done in multiple batches.

### Control environment

Libraries were back diluted to OD 0.008 into 7 ml of PYE or PYE+0.002% xylose and grown overnight until they reached saturation at OD *∼*1.6.

### Heat stress

Libraries diluted to OD of 1 and heat-stressed at low, medium, or high (37, 42, 43.8°C, respectively) for 45 minutes in a Biorad Thermocycler. After 45 minutes, cells diluted back to a final OD of 0.008 in 7 ml media for 24-hour growth.

### Oxidative stress

Libraries were directly diluted back to OD of 0.008 in 7 ml media that contains low, medium, or high (0.025mM, 0.05mM, 0.1mM) level hydrogen peroxide. Cells were grown for 24 hours in these chronic stress conditions.

### Canavanine stress

Libraries were directly diluted back to OD of 0.008 in 7 ml media that contains low, medium, or high (25ug/ml, 50ug/ml, 100ug/ml) levels of L-canavanine. Cells were grown for 24 hours in these chronic stress conditions.

### Library preparation

Following overnight growth, 1 ml of saturated culture from each Tn library was pelleted at 8000xg for 2 minutes. Genomic DNA was extracted by Monarch Genomics DNA Preparation Kit (NEB) according to the manufacturer’s protocol. Sequencing libraries were prepared for Next-generation sequencing via a custom three-step PCR protocol. Indexed libraries were pooled and sequenced on a NextSeq 500 device (Illumina) in the University of Massachusetts Amherst Genomics Core Facility.

### Computational methods

For more detailed descriptions of the computational methods, please refer to the SI Appendix, Supporting Text 1.1-1.7.

### Read mapping and preprocessing

Mapping and preprocessing of the TIS raw data was done as described previously with some modifications (25). Samples were de-multiplexed, and unique molecular identifiers (UMIs) were added during PCR steps removed using Je (26). The clipped reads were mapped to the *Caulobacter crescentus* NA1000 genome (NCBI Reference Sequence: NC011916.1) using bwa and sorted with samtools (27, 28). Duplicate transposon reads removed by Je and indexed with samtools. Genome positions are assigned to the 5*′* position of transposon insertions using bedtools genomecov (29). Subsequently, the bedtools map is used to count either the total number of transposon insertions per gene using the bedtools map -o sum argument or the unique number of insertions using the bedtools map -o count argument.

### Batch correction

We apply ComBat-seq (30) to estimate batch effects and perform library size correction. The unique insertion count data from the transposon insertion sequencing data is used as a response, and the adjusted data, which is integer-valued, is obtained by mapping the quantiles of the empirical distributions of data to the batch-free distributions.

### Classification of fitness effects

Based on the unique insertion counts, the genes are classified as essential, conditionally essential, conditionally beneficial, conditionally detrimental, or conditionally neutral as described previously (21) except median counts were used to increase robustness to outlying values.

### Generalized linear model with local false discovery control

We fit a regularized negative binomial regression model to unique counts to estimate the environmental and genetic fitness effects as done previously (21). We define a regression model for each gene or locus tag in the *Caulobacter crescentus* NA1000 genome. Let the batch-effect adjusted unique insertion count value for gene locus *l*, condition *i*, and replicate *j* be denoted *y*_*ijl*_. We assume that *y*_*ijl*_ follows a negative binomial distribution *NB*(*μ*_*il*_, *ϕ*_*il*_) independently for each *l*. The condition indexed by *i* is equivalent to the combination of the genetic background, *g ∈ 𝒢*; the proteotoxic stress, *e ∈ ℰ*; and stress level, *s ∈ 𝒮*. The model for transposon insertion counts of gene *l* across experiments is:

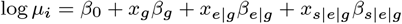

where *β*_0_ is the logarithm of expected counts for control samples. The vector *xg* is an indicator vector that selects the genetic background associated with condition *i*, and the parameter *βg* is the average effect of genetic background *g* on the log transposon counts for gene *l*. The vectors *x*_*e*|*g*_ and *x*_*s*|*e*|*g*_, and the parameters *β*_*e*|*g*_ and *β* _*s* |*e*|*g*_ have a similar interpretation for the stress type and stress level. The parameters for the regularized regression model are estimated by the coordinate descent algorithm as implemented in the `glmnet` package (31). Then, we used the local false discovery rate to control the proportion of false positives in the set of called beneficial/detrimental genes under the assumption that a majority of the genes are non-essential (32).

### EMD distance

To assess the fitness differences between the two stress conditions in a given strain, we utilize the Earth Mover’s Distance (EMD) to compare the median counts (both total and insertion counts) of genes selected through the GLM framework. EMD, also known as the Wasserstein metric, is a measure that quantifies the amount of work required to transform one distribution into another, taking into account both the weight of the distribution that needs to be moved and the distance it has to travel (33).

### Fitness defect

Batch-adjusted unique insertion counts were used to calculate the fitness values for subsequent model-Y knockoff analysis. The fitness values for each strain are

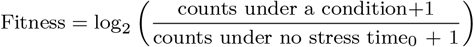

The normalized fitness values allow comparing the changes in the relative abundance of each gene between different samples. We perform a log transformation to transform count data to a Gaussian distribution and add 1 to counts for all the genes before the log transformation to eliminate the negative values or zero denominators in the log function.

### Data subsetting

For subsequent analysis, only conditionally essential, conditionally beneficial, and conditionally detrimental genes derived from the GLM framework were retained.

### Statistical knockoffs

Let *X*_*i*_ encode the *i*-th condition(proteotoxic stress/stress level) and let *Y*_*i*_ encode the fitness value measurement vector in response to the *i*-th condition. For example, for three stress levels (heat, canavanine, oxidative), *X*_*i*_ is an indicator vector for the proteotoxic stress over different stress levels, and *Y*_*i*_ is the *r*-dimensional fitness profile. The roles of *X* and *Y* can be swapped while fitting a model to perform response selection, making the original response variables *Y* the features in the swapped model. The detailed procedure and the key steps are described elsewhere (12).

### Hierarchical Gamma-Poisson model

We analyze the data with a nonparametric Bayesian model based on a Gamma-Poisson hierarchy to identify shared essentiality patterns across conditions within each genetic perturbation strain. Let *y*_*ijl*_ be the count of unique transposon inserts in condition *i*, replicate *j*, and gene locus *l*. The model learns *k ∈ {*1, …, *K}* clusters or networks of genes. The hierarchical Gamma-Poisson model is illustrated as the following:

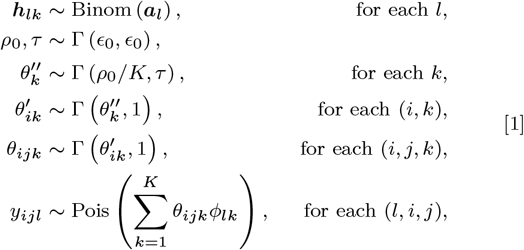

The rate parameter in the Poisson model is the sum of *K* products, denoted as *θ*_*ijk*_ and *ϕ*_*lk*_, where *θ*_*ijk*_ is the propensity of component *K* for sample *j* in condition *i*, and *ϕ*_*lk*_ = ***T***_*l*_***h***_*lk*_ represents the expected number of insertions for gene *l* in component *k*. The model employs an *L ×* 2 matrix, ***T***, to allow for a gene-specific threshold for calling a gene as essential or non-essential.

The term “essential” here indicates a relative reduction in mean insertion counts, signifying a positive fitness contribution. The hyperparameter *a*_*l*_ indicates the prior probability that locus *l* is essential. The common set of essential gene components or networks is represented by *H ∈ {*0, 1*}*^*K×L*^, and the prior for *H* is *h*_*lk*_ *∼* Binom (1, *a*_*l*_).

### Estimation of *T*

The insertion count threshold for calling a gene “essential” can vary from gene to gene. A Gaussian mixture model with two components is fit to each gene to determine the values for each row of the ***T*** matrix, which encodes the information about expected reads for essential/non-essential genes. The batch-adjusted unique insertion counts for all predictive genes for each strain are passed as input to the GaussianMixture function in the sklearn package in Python to estimate the parameters of the model. We restrict the upper bound of the estimated mean of the essential threshold to 10.

### Model inference

The augment-and-marginalize method is used to construct a full analytical steps Gibbs sampler (34). Details can be found at (13, 35) and SI Appendix, Supporting Text 1.7.

## ACKNOWLEDGMENTS

This works was supported by NIH 5R01GM135931. The authors thank the University of Massachusetts Amherst Genomics Core Facility (RRID: SCR 017907) for providing sequencing services.

