## Supplementary Information for "Discovering Genetic Modulators of the Protein Homeostasis System through Multilevel Analysis"

#### This PDF file includes:

Supporting text
Figs. S1 to S42
Tables S1 to S5
SI References

### Supporting Information Text

### 1. Experimental Methods

**A. Tnseq PCR library preparation.** The sequencing libraries for Next-generation sequencing were prepared using a customized three-step PCR protocol (PCR1-2-3). Genomic DNA input was normalized to a concentration of 100 ng/ul. The initial transposon junction amplification (PCR1) was performed using an arbitrary PCR amplification method. This step utilized a forward primer designed (PCR1_F_RBTn5) to align with one end of the transposon and three reverse arbitrary primers (PCR1_R_arb1-2-3). The PCR1 process employed a 2-step cycling protocol with annealing temperatures set at 42°C and 58°C and the number of cycles at 6 and 15, respectively. Subsequently, the second PCR step (PCR2) involved the addition of 16S adapters for Illumina indexing, along with unique molecular identifiers (UMI), and amplification of the library for 36 cycles. Post-PCR2 cleanup was performed using Aline PCRClean DX magnetic beads. The final PCR step (PCR3) incorporated NexteraXT dual indexes in accordance with the manufacturer's protocol. Post-indexed library (PCR3) cleanup was performed using Aline PCRClean DX magnetic beads.

### 2. Computational Methods

**A. Batch correction.** We apply a modified version of ComBat-seq (1), a tool in RNA-seq studies that estimates batch effects using a negative binomial regression model. The unique insertion count data from the transposon insertion sequencing data is used as a response, and the parameter estimation is done by established methods (2–4). Due to hierarchical empirical Bayes modeling, information across genes is pooled to make robust adjustments in case of small sample sizes and/or outlying values. The adjusted data, which is integer-valued, is obtained by mapping the quantiles of the empirical distributions of data to the batch-free distributions (expected distributions if there were no batch effects in the data based on the model).

Here, we describe the ComBat-seq model and adjustment as described in (1): We define a regression model for each gene or locus tag in the *Caulobacter crescentus* NA1000 genome. Let the transposon unique insertion count value for gene i of sample j from batch k be denoted by $y_{ijk}$. We assume that $y_{ijk}$ follows a negative binomial distribution $NB(\mu_{ijk}, \phi_{ik})$,where $\mu_{ijk}$ and $\phi_{ik}$ are the mean and the dispersion parameters. The gene-wise model will be:

$$\begin{aligned} \log \mu_{ijk} &= \beta_0 + \mathbf{X}_j \beta_i + \gamma_{ik} + \log N_j \\ var(y_{ijk}) &= \mu_{ijk} + \phi_{ik} \mu_{ijk}^2 \end{aligned}$$

Where $\beta_0$ denotes the logarithm of expected counts for control samples (wild-type no stress in our case). $\mathbf{X}_j \beta_i$ reflects changes to the log of expected counts due to biological conditions like strain, proteotoxic stress, and stress level, which is preserved in the data after adjustment. In this term, $X_j$ may be an indicator of the strain/condition/stress level for sample j. $\beta_i$ denotes the corresponding regression coefficient. $N_j$ represents the library size, i.e., total unique counts across all genes in sample j. The mean and dispersion batch effect parameters are denoted by $\gamma_{ik}$ and $\phi_{ik}$, respectively, modeling the effect of batch k on the gene i and estimated by established methods like Fisher scoring iteration and Cox-Reid adjusted profile likelihood (APL) (2–4).

**Adjustment** After estimation of $\gamma_{ik}$, $\mu_{ijk}$ and $\phi_{ik}$, parameters for batch-free distributions are calculated from the 'batch-free' negative binomial distribution $NB(\mu^*_{ij}, \phi^*_i)$, where parameters are calculated as

$$\begin{aligned} \log \mu^*_{ij} &= \log \hat{\mu}_{ijk} - \hat{\gamma}_{ik} \\ \phi^*_i &= \frac{1}{N_{batch}} \sum_i \hat{\phi}_{ik} \end{aligned}$$

The adjusted values $y^*_{ij}$ is calculated by finding the closest quantile such that $F^*(y^*_{ij}) = P(y^* \leq y^*_{ij})$ is closest in absolute value to F($y_{ijk}$) = P($y \leq y_{ijk}$). The adjusted counts are then used as a response in the negative binomial regression model.

**B. Categorization of insertions.** The categorization of insertions is done as described previously (5) except the usage of median counts to make it robust in case of outlying values. Let the genes under investigation be $\mathcal{S}$. We partition $\mathcal{S}$ into a set of essential genes, $\mathcal{A}$, and the set of nonessential genes, $\overline{\mathcal{A}} = \mathcal{S} \setminus \mathcal{A}$. Essential genes show no growth when disrupted in a particular strain of interest in standard media. Then, we partition the set of nonessential genes into a set of conditionally essential genes, $\mathcal{B} \subseteq \overline{\mathcal{A}}$, and a set of conditionally nonessential genes, $\overline{\mathcal{B}} = \overline{\mathcal{A}} \setminus \mathcal{B}$. The conditionally essential set contains genes that, when disrupted, still yield growth in no-stress or control conditions but do not grow when the stress conditions are varied; this set depends on the stress condition. To identify genes that produce a quantitative, rather than qualitative, change in fitness as measured by the growth rate, the set of conditionally nonessential genes is further partitioned into the following mutually exclusive, collectively exhaustive sets: conditionally beneficial ($\mathcal{C}$), conditionally neutral ($\mathcal{D}$), and conditionally detrimental ($\mathcal{F}$). If the disruption of a gene decreases growth, it is called conditionally beneficial ($\mathcal{C}$). If the disruption of a gene does not affect growth, it is called conditionally neutral ($\mathcal{D}$). And, if the disruption of a gene improves growth, it is called conditionally detrimental ($\mathcal{F}$).

We denote the genetic background of the experiment $g \in \mathcal{G}$, environmental condition $e \in \mathcal{E}$, and the stress level. For a given combination $(g, e, s)$, the data set contains $R_{ges}$ replicate experiments; we index the replicate with $r$. In experiment $(g, e, s, r)$, there are $N_{gesr}$ observed transposon insertions that are mapped to genes (perhaps excluding some trimmed region around

| **Primers** | **Sequence** | |
|---|---|---|
| PCR1_F_RBTn5 | CGCCCTGCAGGGATGTCCACGAG | |
| PCR1_R_arb1 | GTCTCGTGGGCTCGGAGATGTGTATAAGAGACAGNNNNNNNNNNACGCC | |
| PCR1_R_arb2 | GTCTCGTGGGCTCGGAGATGTGTATAAGAGACAGNNNNNNNNNNCCTGG | |
| PCR1_R_arb3 | GTCTCGTGGGCTCGGAGATGTGTATAAGAGACAGNNNNNNNNNNCCTCG | |
| PCR2_F6_Rbjunc2 | TCGTCGGCAGCGTCAGATGTGTATAAGAGACAGNNNNNNNGCCGGCCGGTTGAGATGTGTA | |
| PCR2_R_universal | GTCTCGTGGGCTCGGAGATG | |
| **PCR1 Thermocycling conditions** | | |
| 95C | 3min | |
| 95C | 30s | 6 cycles |
| 42C | 30s | |
| 72C | 40s | |
| 95C | 30s | 15 cycles |
| 58C | 30s | |
| 72C | 40s | |
| 72C | 1min | |
| 4C | Forever | |
| **PCR2 Thermocycling conditions** | | |
| 95C | 1min | |
| 95C | 30s | 36 cycles |
| 60C | 20s | |
| 72C | 40s | |
| 72C | 1min | |
| 4C | Forever | |

**Table S1. Tnseq PCR library preparation primers and thermocycling conditions**

the start and stop codon of the gene). We primarily use the count of unique insertions. The median unique counts across $r$ replicates is $\bar{y}^{\text{uniq}}_{gesi}$. We define the control condition to be a wild-type genetic background when no stress is applied within standard PYE media, and we denote this condition $(g', e', s')$. The median control condition counts for gene $i$ are then $\bar{y}^{\text{uniq}}_{g'e's'i}$. We estimate the set of *conditionally essential genes* in condition $(g, e, s)$ as $\hat{\mathcal{B}}_{ges} = \left\{ i | \bar{y}^{\text{uniq}}_{gesi} \leq 1, \forall i \in \overline{\mathcal{A}} \right\}$

**C. Negative binomial regression model.** We define a regression model for each gene or locus tag in the *Caulobacter crescentus* NA1000 genome. Let the batch-effect adjusted unique insertion count value for gene i of sample j be denoted by $y_{ij}$. We assume that $y_{ij}$ follows a negative binomial distribution $NB(\mu_{ij}, \phi_i)$, where $\mu_{ij}$ and $\phi_i$ are the mean and the dispersion parameters. The gene-wise model will be:

$$\begin{aligned} \log \mu_{ij} &= \beta_0 + \mathbf{X}_g \beta_g + \mathbf{X}_{e|g} \beta_{e|g} + \mathbf{X}_{s|e|g} \beta_{s|e|g} \\ var(y_{ij}) &= \mu_{ij} + \phi_i \mu_{ij}^2 \end{aligned}$$

where $\beta_0$ denotes the logarithm of expected counts for control samples. $\mathbf{X}_g \beta_g$ reflects changes to the log of expected counts due to mutant background, which is preserved in the data after adjustment. $\mathbf{X}_g$ indicates the mutant background for sample j. $\beta_g$ denotes the corresponding regression coefficient. $\mathbf{X}_{e|g} \beta_{e|g}$ reflects changes to the log of expected counts due to proteotoxic stress nested within the mutant background. $\beta_{e|g}$ indicates the proteotoxic stress (condition) nested within the mutant background for sample j. $\beta_{e|g}$ denotes the corresponding regression coefficient. $\mathbf{X}_{s|e|g} \beta_{s|e|g}$ reflects changes to the log of expected counts due to stress level nested within proteotoxic stress and mutant background. $\beta_{s|e|g}$ indicates the stress level nested within proteotoxic stress and mutant background for sample j. $\beta_{s|e|g}$ denotes the corresponding regression coefficient.

**D. Regularization.** Inflated regression coefficients and high variance in the model can occur due to the low or sparse counts in response variables, resulting in poor model generalization. Filtration (6) or the adding pseudo-counts (7) have been proposed to address this issue. Regularization can be applied to generalized linear models to avoid generalization issues. A regularized regression model can also be viewed as a penalized likelihood function:

$$\ell(\beta; y) = \ell(\beta; y) - \lambda \sum_{j=1}^{p} \beta_j^{c2}$$

where $\ell(\beta, \theta; y)$ is the log-likelihood function of Negative binomial distribution and $\sum_{j=1}^{p} \beta_j^{c2}$ is the ridge penalty($l_2$) function with tuning parameter $\lambda$ which controls the amount of shrinkage and size of coefficients. The parameters for the penalized count regression are estimated by the coordinate descent algorithm as implemented in the `glmnet` package (8).

**E. Local false discovery rate.** Our primary goal is to identify the conditionally beneficial/detrimental effects among a large number of effects after fitting the regularized negative binomial generalized linear model to unique counts independently for each gene $i$. Having a larger number of effects( both beneficial/detrimental and non-essential) allows the possibility of local inference where an effect such as $z = -1.96$ is judged based on the empirical distribution and not with respect to the theoretical possibility of more extreme results. We used the local false discovery rate to control the proportion of false positives in the set of called beneficial/detrimental genes (9) under the assumption that a majority of the genes modeled are non-essential. Suppose that each of the N z-values falls into one of two classes, “non-essential” or ”beneficial/detrimental” corresponding to whether or not generated $z_i$ is equal to zero, with prior probabilities $p_0$ and $p_1 = 1 - p_0$. Assume that $z_i$ has a density of either $f_0(z)$ or $f_1(z)$ depending on its class. Then

$$\begin{aligned} p_0 &= Pr(\text{non-essential}) \\ p_1 &= Pr(\text{beneficial/detrimental}) \\ f_0(z) &= \text{density if gene is non-essential} \\ f_0(z) &= \text{density if gene is beneficial/detrimental.} \end{aligned}$$

Then the mixture densiy $f(z)$ will be $p_0 f_0(z) + p_1 f_1(z)$. The Bayes posterior probability that a case is non-essential given z, by definition, the local false discovery rate, is

$$fdr(z) \equiv Pr(\text{nonessential}|z) = p_0 f_0(z)/f(z)$$

Estimation of $f(z)$ and calculation of fdr values is done by `locfdr` package (9). The local nature of $fdr(z)$ helps us to interpret the results from the individual effects under each mutant background, nested proteotoxic stress, and stress levels.

**F. Knockoffs.**

**Formulation** The key ideas of applying $Y$ knockoffs are described in detail in (10). Briefly, Let $X_i$ encode the $i$-th condition(proteotoxic stress/stress level) and let $Y_i$ encode the fitness value measurement vector calculated as described before in response to the $i$-th condition. Assume we have $n$ i.i.d. random variables $(X_i, Y_i)$, where $X_i \in \mathbb{R}^p$, $Y_i \in \mathbb{R}^r$ assembled into two data matrices $\mathbf{X} \in \mathbb{R}^{n\times p}$ and $\mathbf{Y} \in \mathbb{R}^{n\times r}$ such that the $i$-th row of $\mathbf{X}$ is $X_i$ and the $i$-th row of $\mathbf{Y}$ is $Y_i$. For example, for three stress levels (heat, canavanine, oxidative), $X_i$ is an indicator vector for the proteotoxic stress over different stress levels, and $Y_i$ is the $r$-dimensional fitness profile. A response variable $Y_j$ for $j \in \{1, \dots, r\}$ is said to be unimportant if and only if $Y_j$ is independent of $X$ conditionally on the other responses $Y_{-j}$, where $Y_{-j} = \{Y_1, \dots, Y_r\} \backslash \{Y_j\}$. The set of unimportant variable indices is denoted by $\mathcal{H}_0$, and we call a variable $Y_j$ important if $j \notin \mathcal{H}_0$. The set of the important variable indices is denoted as $\mathcal{S}$. The unimportant responses are conditionally independent of the covariates given the important responses $\{Y_j\}_{j\in\mathcal{H}_0} \perp\!\!\!\perp X \mid \{Y_j\}_{j\in\mathcal{S}}$. The goal of $Y$ knockoffs is to identify the most influential fitness predictors while keeping the FDR under control, where $\mathrm{FDR} = \mathbb{E}\left[|\hat{\mathcal{S}} \cap \mathcal{H}_0|/(|\hat{\mathcal{S}}| \vee 1)\right]$ and $\hat{\mathcal{S}}$ denotes a selected subset of the important response variable indices.

The key ideas of Model-Y Knockoffs are described in detail in (10). In Model-X knockoffs, marginal distribution $F_X$ is known exactly, allowing the conditional distribution $F_{Y|X}$ to be unspecified. Using the fact that the joint distribution can be factorized as $F_{XY} = F_{X|Y}F_Y$ and abundant information availability about the marginal distribution $F_Y$, we can define model-Y knockoffs taking advantage of symmetry as below :

**Model-Y Knockoffs.** A random vector $\tilde{Y} = (\tilde{Y}_1, \dots, \tilde{Y}_r)$ is the model-Y knockoffs of a random vector $Y = (Y_1, \dots, Y_r)$ if it satisfies two properties: (1) Pairwise exchangeability: for any subset $\mathcal{D} \subset [r]$, $(Y, \tilde{Y})_{\mathrm{swap}(\mathcal{D})} \overset{\mathrm{d}}{=} (Y, \tilde{Y})$, i.e. swapping the entries $Y_j$ and $\tilde{Y}_j$ for each $j \in \mathcal{D}$ leaves the joint distribution invariant; and (2) Conditional independence: $\tilde{Y} \perp\!\!\!\perp X \mid Y$, i.e., $\tilde{Y}$ is independent of features $X$ given the response $Y$, which can be guaranteed by constructing $\tilde{Y}$ without looking at $X$.

The theorem to establish the finite sample FDR control for response variables as proposed in (10) is described below

**Model-Y Knockoffs:** Let $W_j = w_j([\mathbf{Y}, \widetilde{\mathbf{Y}}], \mathbf{X})$ be the knockoff statistic to measure the importance for each $Y_j$, $j \in [r]$. Here, functions $w_j, j \in [r]$ are required to satisfy the following flip-sign property:

$$w_j\left([\mathbf{Y}, \widetilde{\mathbf{Y}}]_{\mathrm{swap}(\mathcal{D})}, \mathbf{X}\right) = \begin{cases} w_j([\mathbf{Y}, \widetilde{\mathbf{Y}}], \mathbf{X}), & j \notin \mathcal{D}, \\ -w_j([\mathbf{Y}, \widetilde{\mathbf{Y}}], \mathbf{X}), & j \in \mathcal{D}. \end{cases} \qquad [1]$$

For any $q \in [0, 1]$, let

$$\tau = \min\left\{t > 0 : \frac{1 + |\{j : W_j \le -t\}|}{|\{j : W_j \ge t\}|} \le q\right\}. \qquad [2]$$

Then we can control the $\mathrm{FDR} = \mathbb{E}\left[|\hat{\mathcal{S}} \cap \mathcal{H}_0|/(|\hat{\mathcal{S}}| \vee 1)\right]$ at the target level of $q$ by selecting importance responses

$$\hat{\mathcal{S}} = \{j : W_j \ge \tau\}.$$

**Algorithm** The knockoffs can still be validly generated by swapping the roles of $X$ and $Y$ in a variable response selection as shown in (11). The roles of $X$ and $Y$ can be swapped while fitting a model to perform response selection, making the original response variables $Y$ the features in the swapped model. We summarize the key steps in the Algorithm as described in (11). The algorithm has three steps. The first step is to generate knockoffs $\tilde{Y}$ that satisfy the Model-Y Knockoffs Definition. The second step is to define the feature importance measure and the knockoff statistic for each $Y_j$, $j \in [r]$, where $[r] = \{1, 2, \dots, r\}$. The third step is to decide the filtering threshold to guarantee the controlled FDR level.

**Algorithm** Response Selection using Knockoffs

1: Generate the knockoffs of the original response variable such that

$$(\tilde{Y}, Y) \sim \mathcal{N}(\mathbf{0}, \mathbf{G}), \text{ s.t. } \mathbf{G} = \begin{pmatrix} \boldsymbol{\Sigma} & \boldsymbol{\Sigma} - \mathrm{diag}\{\mathbf{s}\} \\ \boldsymbol{\Sigma} - \mathrm{diag}\{\mathbf{s}\} & \boldsymbol{\Sigma} \end{pmatrix}.$$

2: Construct the response importance measure and knockoff statistics using Lasso with Model-Y Knockoffs (Lasso-MYK)

3: Filter the important response using the threshold $\tau$ in EquationFigure 2 with the target FDR level $q$.

**Generalized Linear Model-Y Knockoffs** The response selection generalized linear model setting is performed by fitting a regularized multinomial logistic regression model using Lasso (12). The knockoffs $\tilde{Y}$ are augmented to the original response $Y$ in the role of the covariate variables and the condition indicators (heat, canavanine, oxidative) $X$ in the role of the response variable. The optimal Lasso solution is

$$\hat{\beta}(\lambda) \in \underset{\beta\in\mathbb{R}^{2r\times p}}{\arg\min}\left\{\frac{1}{2}\|\mathbf{Y} - [\mathbf{X}, \widetilde{\mathbf{X}}]\beta\|_2^2 + \lambda\|\beta\|_1\right\}.$$

We define $Z_j = \sup\left\{\lambda : \hat{\boldsymbol{\beta}}_j(\lambda) \neq 0\right\}$ to be the point, $\lambda$, on the Lasso path at which the response $Y_j$ first enters the model. The standard knockoff statistics $W_j = Z_j - \tilde{Z}_j$ are computed, and a set of important responses, $\hat{\mathcal{S}}$, are selected with a controlled FDR of 0.1.

**G. Hierarchical Gamma-Poisson Model.** Our transposon sequencing experiment data has a nested experimental design as illustrated in Figure S1. Briefly, the genetic background is first generated by deleting possible homeostasis components (clpA, clpB, lon, DnaK) or wild-type; each genetic background is subjected to multiple stress conditions (canavanine, heat, oxidative). For each stress condition, strain is grown under multiple stress levels, and replicates are collected to reduce the effect of technical variation in the experimental measurements.

Let $y_{ijl}$ be the count of unique transposon inserts in a condition or proteotoxic stress $i$, stress level $j$, and locus (gene) $l \in \{1, \ldots, L\}$. The common set of essential gene components or networks are represented by a $K \times L$ matrix $H$; wherein an entry $h_{kl} \in \{0, 1\}$ indicates that gene $l$ in the network is essential. Here "essential" means the relative reduction of mean insertion counts indicating a positive fitness contribution. Let $a_l$ denote the prior probability that locus $l$ is essential, and independent prior for $H$ can be denoted by $h_{lk} \sim \text{Multi}\,(1, a_l)$. The $a_l$ denotes the prior probability that locus $l$ is essential, and independent prior for $H$ is denoted by $h_{lk}$. We encode the information on how many insertions should be essential or non-essential in transition matrix $T$, which is a $L \cdot 2$ matrix. The rate parameter in Poisson is the summation of K products, where $\theta_{ijk}$ is the propensity of component K for sample j in stress or condition i(drawn by the gamma distribution) and $\phi_{lk} = \boldsymbol{T}_l \cdot \boldsymbol{h}_{lk}$ is the expected number of insertions for the gene at l in component k.

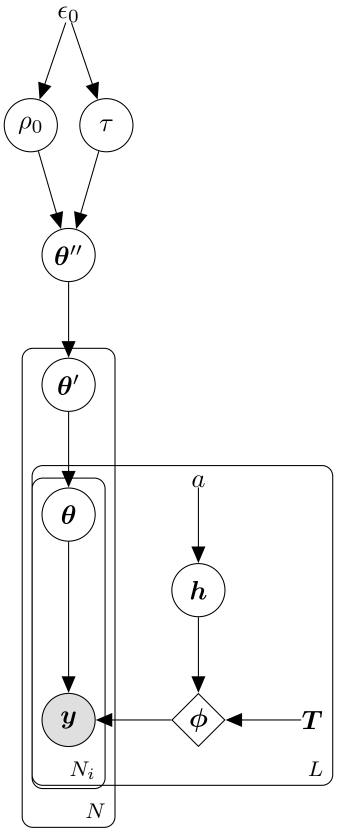


**Graphical Model of hGP** This graph shows the generative process of hGP. Adopted from (13, 14).

The details of the sampling algorithm are summarized as follows:

**Algorithm 1 Sampling Algorithm** Auxiliary variable Gibbs sampler for Gamma-Poisson Model. Adopted from (13, 14)

**for each** (i,j,l) **do** Sample $\left(y_{ijl}^{(k)}\right)_{k=1}^{K} | y_{ijl}, \theta_{ijk}, \phi_{lk} \sim \text{Multi}\left(y_{ijl}, \left(\frac{\theta_{ijk}\phi_{lk}}{\sum_{k'=1}^{K}\theta_{ijk'}\phi_{lk'}}\right)_{k=1}^{K}\right)$

**for each** (i,j,k) **do** Sample $w_{ijk} | y_{ij}^{(k)}, \theta'_{ik} \sim \text{CRT}\left(y_{ij}^{(k)}, \theta'_{ik}\right)$

**for each** (i,k) **do** Sample $w'_{ik} | w_{ik}, \theta''_{k} \sim \text{CRT}\left(w_{ik}, \theta''_{k}\right)$

**for each** (k) **do** Sample $w''_{k} | w'_{k}, \rho_0 \sim \text{CRT}\left(w'_{k}, \rho_0/K\right)$

Sample $\rho_0 | w''_1, \ldots, w''_K \sim \Gamma\left(\epsilon_0 + \sum_{k=1}^{K} w''_k, \epsilon_0 + \log(1 + N\log(1 + N_i\log(1+\phi_{lk})))/\tau\right)$

**for each** (k) **do** Sample $\theta''_{k} | w'_{ik}, \rho_0, \tau \sim \Gamma\left(\rho_0/K + w'_k, 1 + N\log(1 + N_i\log(1+\phi_{lk}))\right)$

**for each** (i,k) **do** Sample $\theta'_{ik} | w_{ik}, \theta''_{k} \sim \Gamma\left(\theta''_k + w_{ik}, 1 + N_i\log(1+\phi_{lk})\right)$

**for each** (i,j,k) **do** Sample $\theta_{ijk} | y_{ijk}, \theta'_{ik} \sim \Gamma\left(\theta'_{ik} + y_{ijk}, 1 + \phi_{lk}\right)$

**for each** (l,k) **do** Sample $h_{lk} \sim \text{Multi}\left(h_{lk} | y_{ijl}^{(k)}, a_l, \theta_{ijk}\right)$

### 3. Results

**A. Genome-wide analysis of conditional essentiality.** The experimental setup is illustrated in Figure S1. Sequencing is used to quantify the frequency of transposon-induced mutations, and a set of conditionally essential, beneficial, or detrimental genes for each target environment (and stress) is identified by applying a regularized negative binomial regression combined with local FDR testing in a general linear model (GLM) framework described in the methods section.

As described in (5), two types of count measures can be studied in transposon sequencing: Total and unique. For our modeling purposes, we primarily use unique counts since they nicely capture site-specific variations and satisfy the assumptions of many models. The count measures are sometimes prone to library and batch size variations. The library size is studied by summing up the counts across all the insertion sites in a gene. Our data is also generated for multiple replicates in five batches. For unique insertion counts, we observe an average library size of 89,739 insertions across all batches. A more detailed breakdown of library size by strain, batch, and condition for unique insertions is reported in Table S2 and Figure S2. Overall, wild-type and $\Delta lon$ strains show similarly enriched insertion counts, a pattern reflected even after the library size correction.

In some cases, there might be an inconsequential insertion hotspot near the tail ends due to an amplification error or a genome-specific bias, causing some unique insertions to have larger than expected enrichment. The majority of data is left-skewed, as expected, with most of the count in the biological experiments. As described in methods, we apply a modified version of ComBat-seq (1) to correct for both batch and library size effects. After the correction, we primarily observe higher median unique insertion counts across all genes in wild-type and $\Delta lon$ strains compared to $\Delta clpA$, $\Delta clpB$, and $\Delta lon$. We observe that overall fitness is much more sensitive to a homeostatic component deletion than stress, as illustrated in Figure S3.

For this study, we define "Essential genes" as genes that show no growth when disrupted in a no-stress or control condition in standard media of a strain. The *Caulobacter crescentus* has a total of 4084 genes, including many RNA genes. The wild-type has 580 essential genes, $\Delta lon$ has 621, whereas $\Delta clpA$, $\Delta clpB$, and dnaKJ-NI have 964, 1183, and 1103 essential genes, respectively. The number of conditionally beneficial genes are much more sensitive to proteotoxic stress and stress levels than a typical gene deletion, suggesting an absence of compensatory mechanisms and the cell's inability to cope with severe environmental stresses, reflecting their importance in protein homeostasis. For example, under oxidative stress, under $\Delta lon$, the system is much more sensitive to the stress level (medium vs. high). Many conditionally beneficial genes in oxidative stress, like KatG, are degraded or inactivated when subject to higher stress reflecting their vital role in stress response and fitness adaptation Figure S4. A certain gene may be conditionally beneficial under a particular condition but may change its essentiality under a different proteotoxic stress or stress level Figure S9. A specific gene may also be beneficial or detrimental under several conditions but have various effects on fitness (Figure S5, Figure S7, Figure S8, Figure S6). When we look into the overlap size of the conditionally essential genes as illustrated in Figure S10, a higher overlap is seen within stresses of dnaKJ-NI and wild-type, $\Delta lon$ to a certain extent. The conditionally beneficial genes show lower overlap as illustrated in Figure S11. The wild-type stresses have a higher degree of overlap in conditionally detrimental genes compared to the deficient strains, as illustrated in Figure S12.

**B. Identification of most predictive fitness determinants in a protein homeostasis system.** Using the model-Y knockoff framework, we selected 20 genes in wild-type, 33 genes in $\Delta lon$, 39 genes in $\Delta clpA$, 44 genes in $\Delta clpB$, and 38 genes in dnaKJ-NI as fitness predictors. We selected a total of 146 genes combined across all strains as fitness predictors. Among 146 genes, 19 genes (excluding CCNA_0375) are shared among at least two strains, as shown in Table S3. The shared fitness predictors are primarily involved in ATP/ADP, DNA-binding, and cytoplasmic enzyme activity.

Among the novel genes (functionally uncharacterized), CCNA_02154 is identified as a universal fitness determinant across all strains (wild-type, $\Delta lon$, $\Delta clpA$, $\Delta clpB$, dnaKJ-NI). The CCNA_02154 is also observed to be more sensitive to canavanine stress than heat or oxidative and is predicted to be involved in acyltransferase activity.

The CCNA_03861 is a universal fitness determinant in $\Delta clpA$, $\Delta clpB$, and dnaKJ-NI.The fitness values for the fitness predictors for each strain are illustrated in Figure S9, S13, S14, S15, S16, S17. Among the well-studied genes, the CCNA_00922 was identified as a universal fitness determinant in wild type, $\Delta lon$, $\Delta clpA$, and dnaKJ-NI. The CCNA_00922 is a well-known chaperone protein involved in protein refolding. The catalase-peroxidase *KatG* (CCNA_03138) is a fitness determinant in wild-type, $\Delta lon$, $\Delta clpB$, and dnaKJ-NI. It is involved in the hydrogen peroxide catabolic process. The *LysR*-family transcriptional regulator (CCNA_03811) is a fitness determinant in wild-type, $\Delta clpB$ and dnaKJ-NI. CCNA_03861 is a universal fitness determinant in $\Delta clpA$, $\Delta clpB$, dnaKJ-NI and is predicted to be a pyridoxal phosphate homeostasis protein. Within each strain, most of the fitness predictors show conserved fitness values across proteotoxic stresses ( Heat, Oxidative, and Canavanine) but are much more sensitive to the stress level ( low, medium, high). After deletion of the homeostasis component, fitness predictors are much more robust to a stress condition, with some predictors sensitive to the level of stress. A more detailed view is presented in S13, S14, S15, S16, S17.

**C. Studying shared essentiality among the proteotoxic stresses.** In the wild-type, the posterior mass of heat stress is highly concentrated on component-1 compared to other components, besides showing some mass in components-2, 4, 8 as shown in Figure S20. In component-1, CCNA_00922 and CCNA_00001 are two genes that are classified as "essential" (insertion counts decrease relatively, indicating a positive fitness contribution). The CCNA_00922 is a ClpXP protease complex that plays a key role in protein refolding and has previously been reported to show a response to heat stress. The sensitivity of CCNA_00922 to heat stress compared to other stresses can be clearly seen in S13 and S29. We see the posterior mass of heat stress is highly concentrated on component-5 compared to other components. In component-5, CCNA_03811, CCNA_03138,

CCNA_02646, and CCNA_030375 are the genes that are classified as "essential". Our data shows that the catalase-peroxidase *KatG* (CCNA_03138) is sensitive to oxidative stress compared to other stresses S13, S38. Similarly, *LysR*-family transcriptional regulator (CCNA_03811) is much more sensitive to oxidative stress as shown in S13, S40.

In the dnaKJ-NI, component-1 has somewhat similar posterior mass concentrations among the canavanine, none, and oxidative stresses, indicating a potential shared network among the stresses related to component-1 (Figure S21). The CCNA_00708 is the only gene that is classified as "essential" in component-1, indicating the importance of its fitness contribution and robustness to stress under dnaKJ-NI. Figure S25 and S28 show the strain-specific fitness conservation of CCNA_00708 in dnaKJ-NI compared to the wild-type.

In the dnaKJ-NI, we see the posterior mass of heat stress is concentrated on component 2 compared to other components. In component-2, besides CCNA_00922, CCNA_01103 is shown in our data to be sensitive to heat stress (S17, S31). The oxidative stress has a relatively higher mass in component-5 in which besides CCNA_03811 and CCNA_00708, CCNA_00293 has shown oxidative specific stress response in our data (S27). The posterior distribution plots for other strains are presented in S22, S23, and S24.

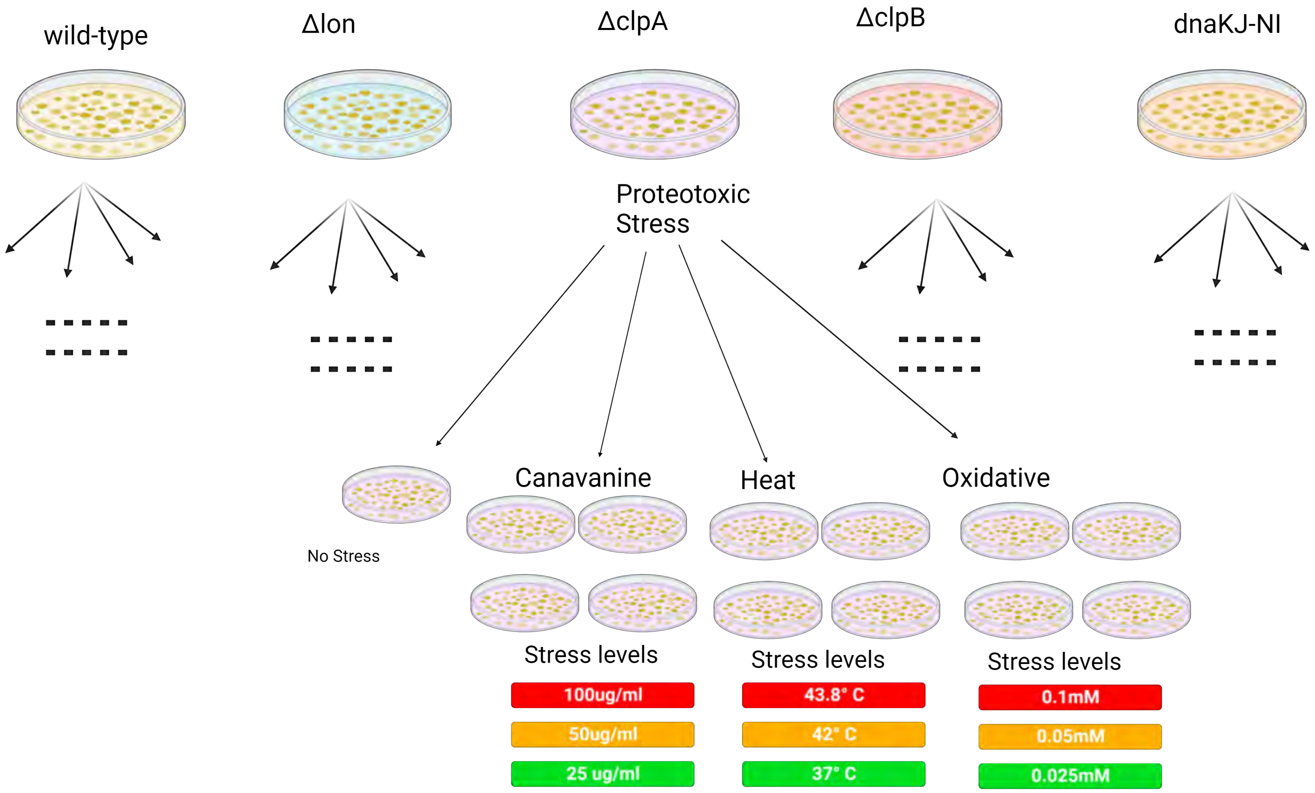


**Fig. S1.** **A schematic representation of the experimental data generated to study proteostasis network** We first constructed transposon libraries in wild-type *Caulobacter crescentus* and strains deficient in specific chaperone or protease genes: *clpA*, *clpB*, *lon*, *dnaK* responsible for protein homeostasis. These libraries were subjected to three different proteotoxic stresses (canavanine, heat, and oxidative) at three levels (low, medium, and high).

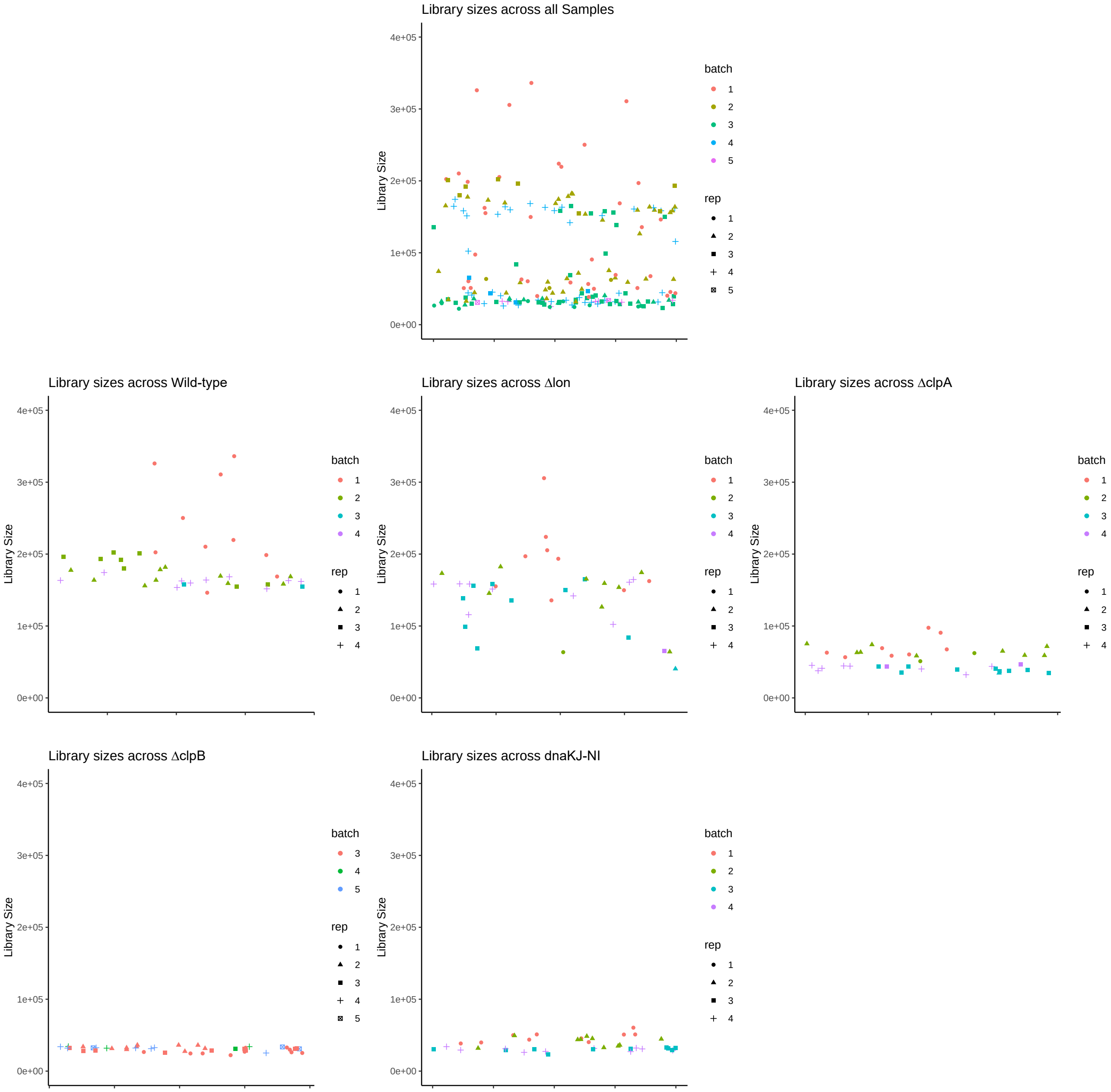


**Fig. S2. Library sizes for unique insertion counts across batches, replicates, and strains** The Y-axis represents the library size obtained by summing up the counts across all the insertion sites in a gene.

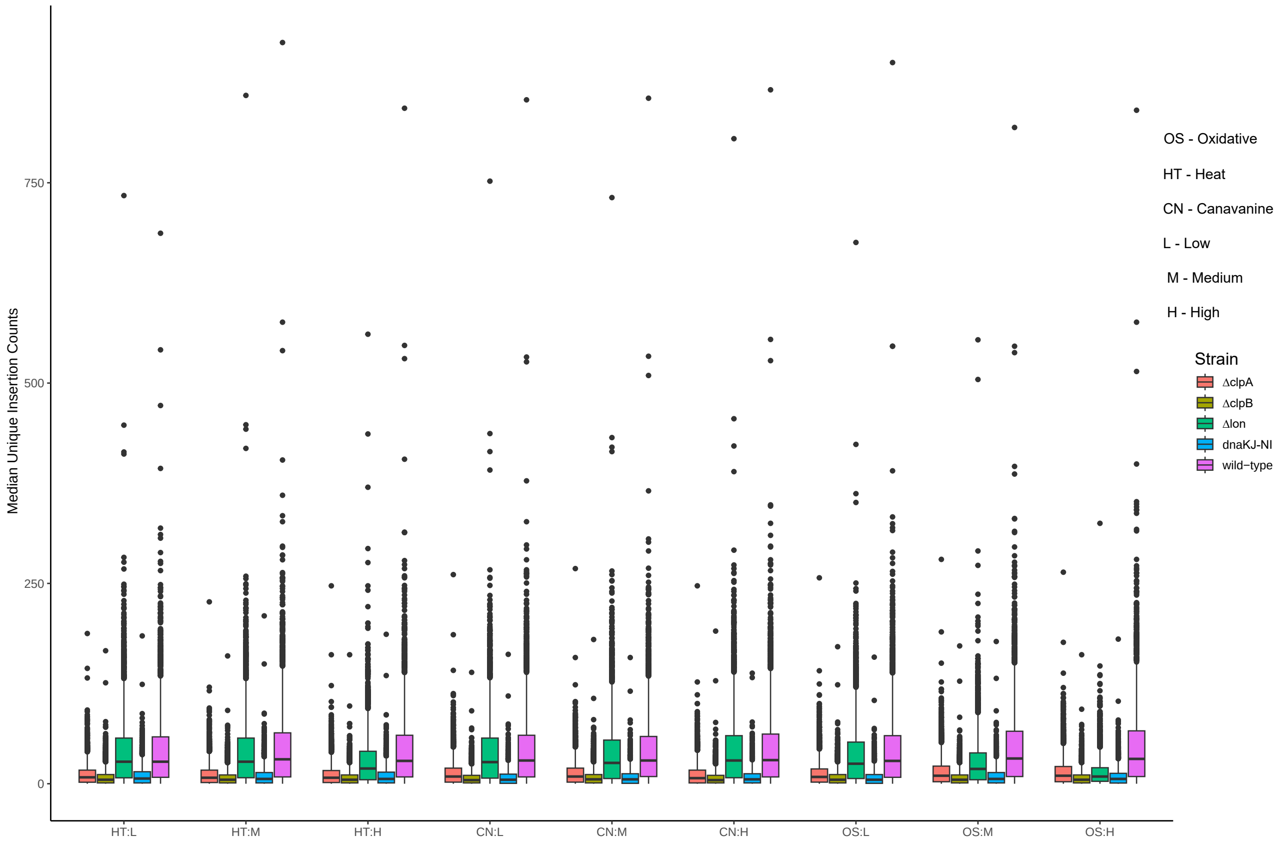


**Fig. S3. Median Unique insertion counts after the adjustment** The median of the batch and library size adjusted unique insertion counts are calculated across stress levels and replicates.

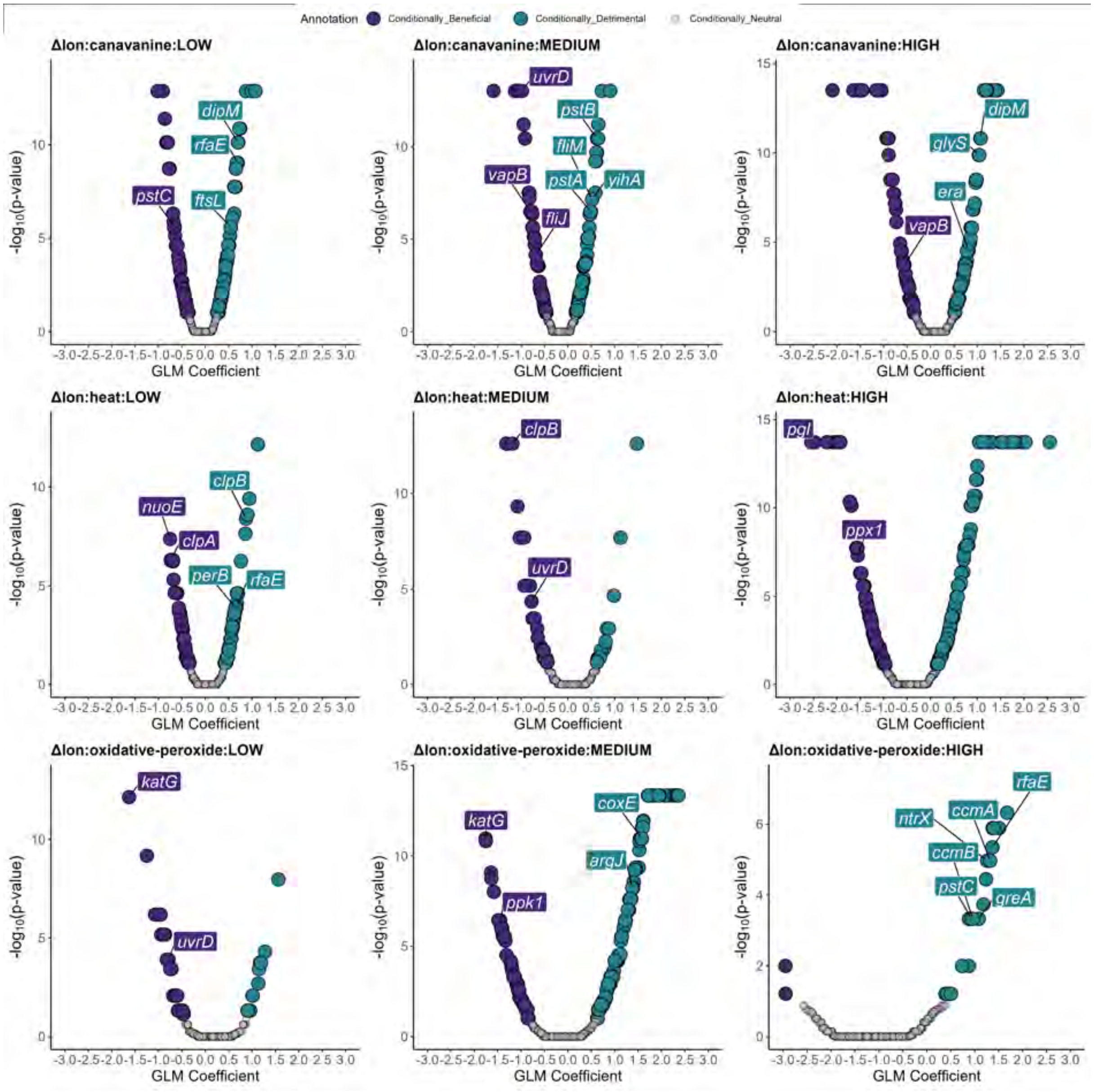


**Fig. S4. Conditionally beneficial and detrimental genes under different proteotoxic stresses for the $\Delta lon$.** The Y-axis represents local FDR $-log_{10}$(P) value, and the x-axis represents coefficients from the regularized negative binomial GLM model. Points are colored based on whether they are conditionally beneficial, neutral, or detrimental. Each proteotoxic stress (canavanine, heat, oxidative) is subjected to three stress levels. (low, medium, high).

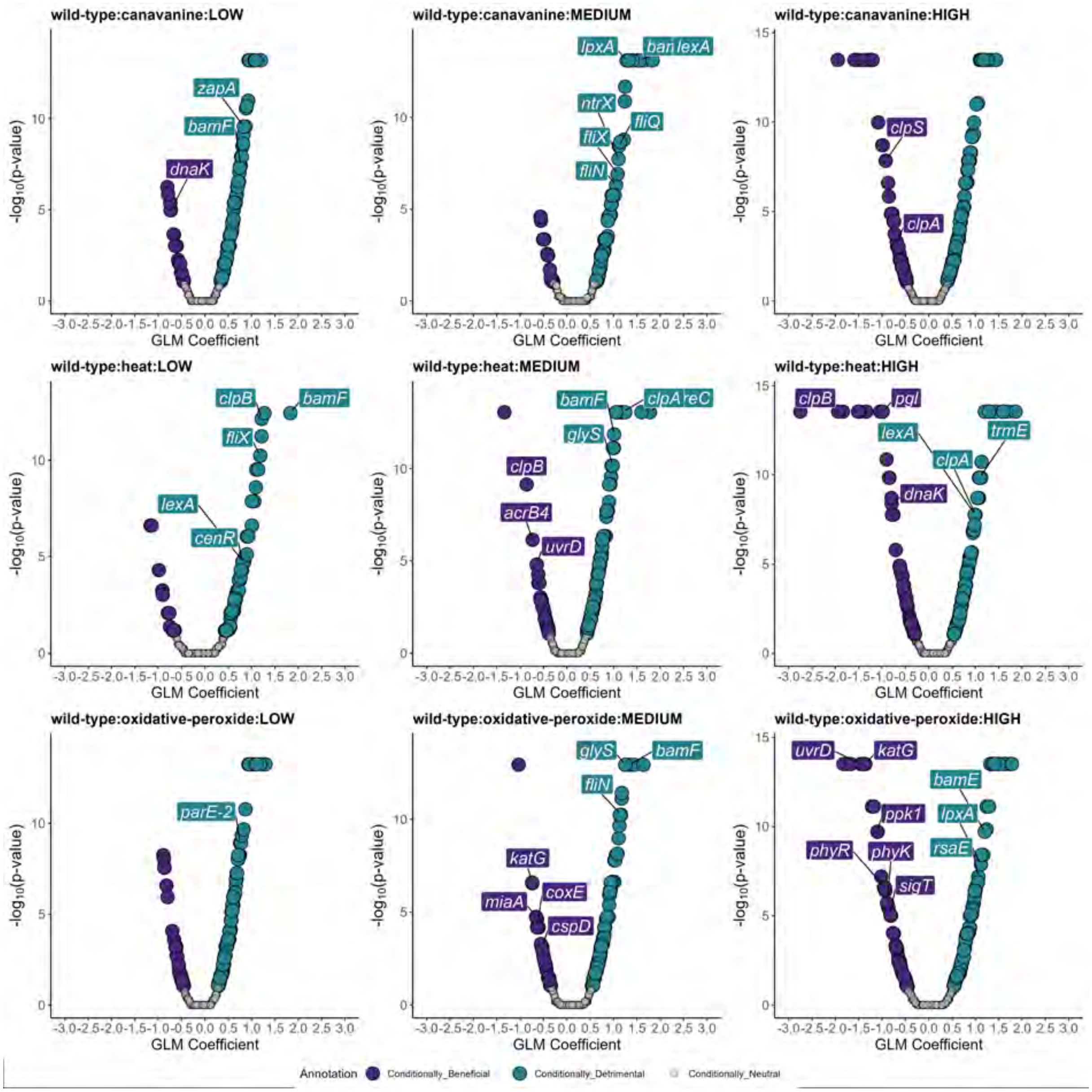


**Fig. S5. Conditionally beneficial and detrimental genes under different proteotoxic stresses for the wild-type.** The y-axis represents local FDR $-log_{10}$(P) value, and the x-axis represents coefficients from the regularized negative binomial GLM model. Points are colored based on whether they are conditionally beneficial, neutral, or detrimental. Each proteotoxic stress (canavanine, heat, oxidative) is subjected to three stress levels (low, medium, high).

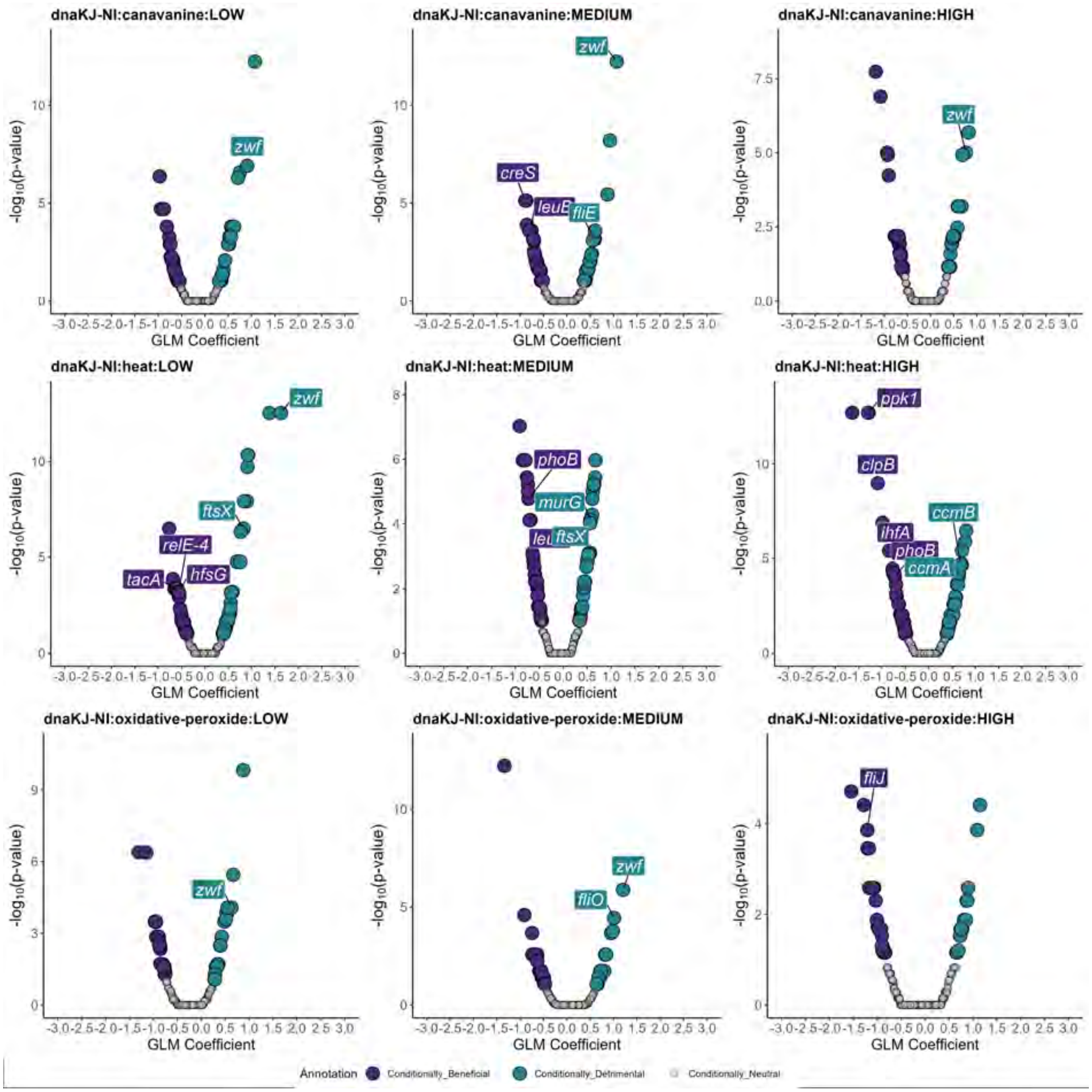


**Fig. S6. Conditionally beneficial and detrimental genes under different proteotoxic stresses for the dnakJ-NI.** The Y-axis represents local FDR $-log_{10}$(P) value, and the x-axis represents coefficients from the regularized negative binomial GLM model. Points are colored based on whether they are conditionally beneficial, neutral, or detrimental. Each proteotoxic stress (canavanine, heat, oxidative) is subjected to three stress levels (low, medium, high).

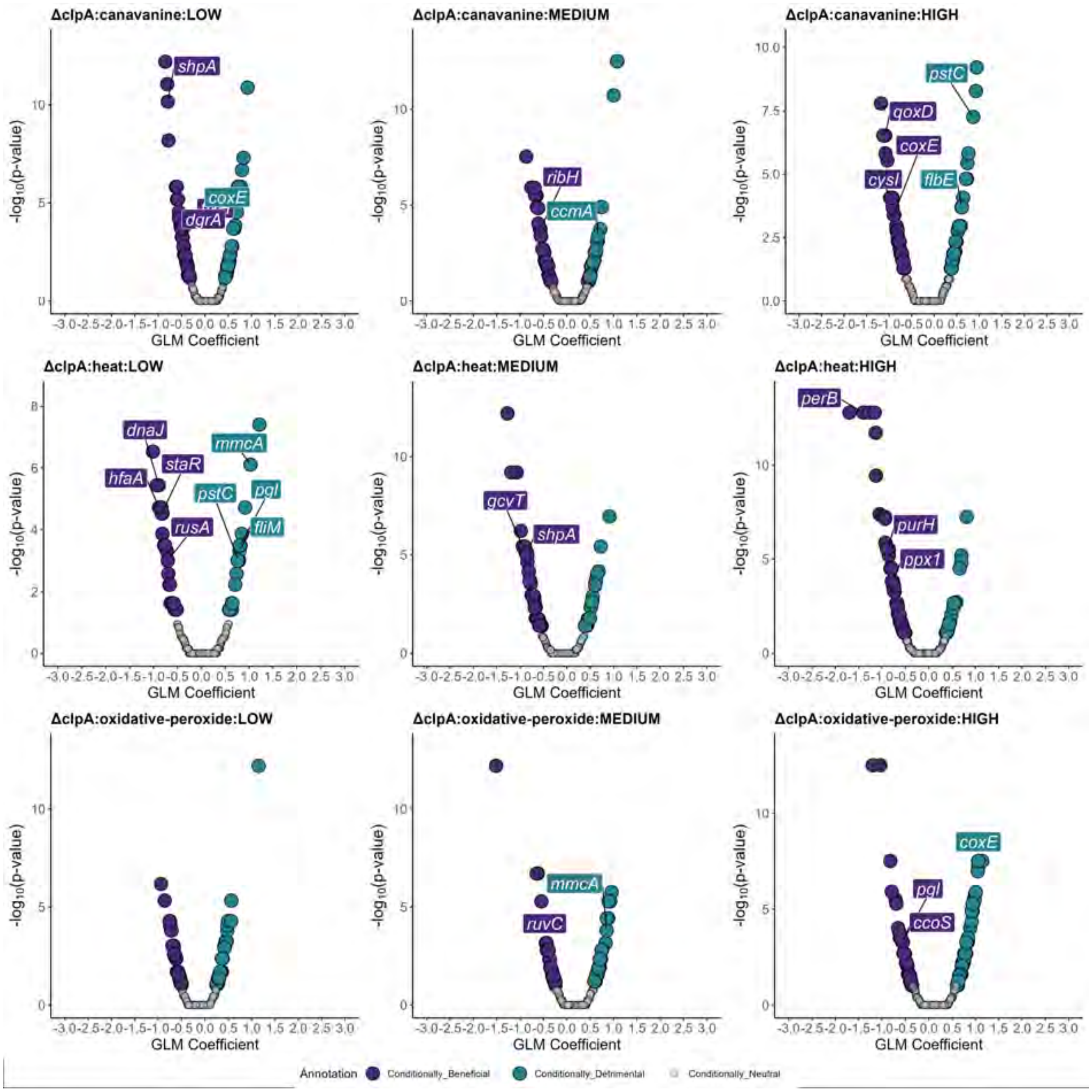


**Fig. S7. Conditionally beneficial and detrimental genes under different proteotoxic stresses for the $\Delta clpA$.** The Y-axis represents local FDR $-log_{10}$(P) value, and the x-axis represents coefficients from the regularized negative binomial GLM model. Points are colored based on whether they are conditionally beneficial, neutral, or detrimental. Each proteotoxic stress (canavanine, heat, oxidative) is subjected to three stress levels. (low, medium, high)

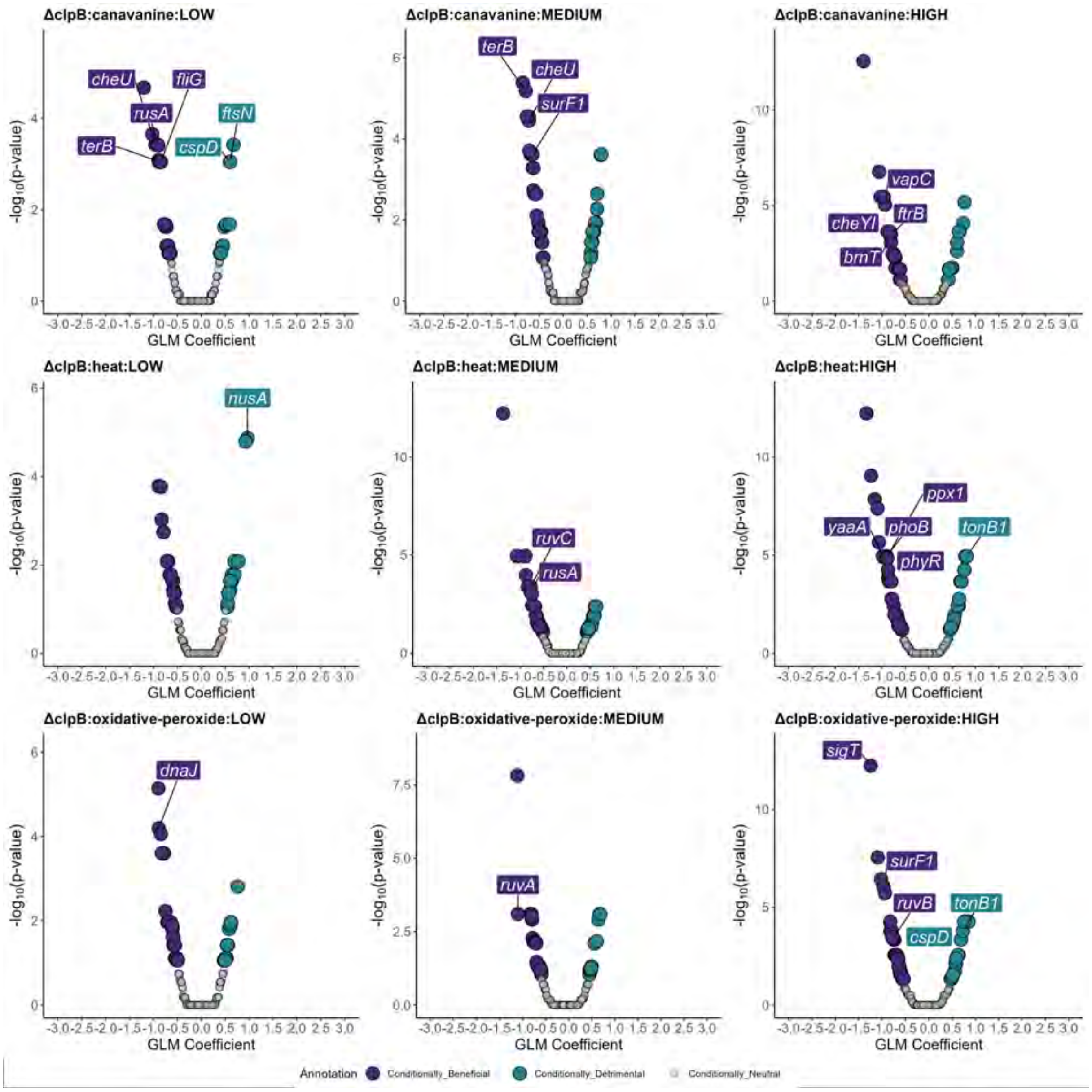


**Fig. S8. Conditionally beneficial and detrimental genes under different proteotoxic stresses for the $\Delta clpB$.** The Y-axis represents local FDR $-log_{10}$(P) value, and the x-axis represents coefficients from the regularized negative binomial GLM model. Points are colored based on whether they are conditionally beneficial, neutral, or detrimental. Each proteotoxic stress (canavanine, heat, oxidative) is subjected to three stress levels. (low, medium, high)

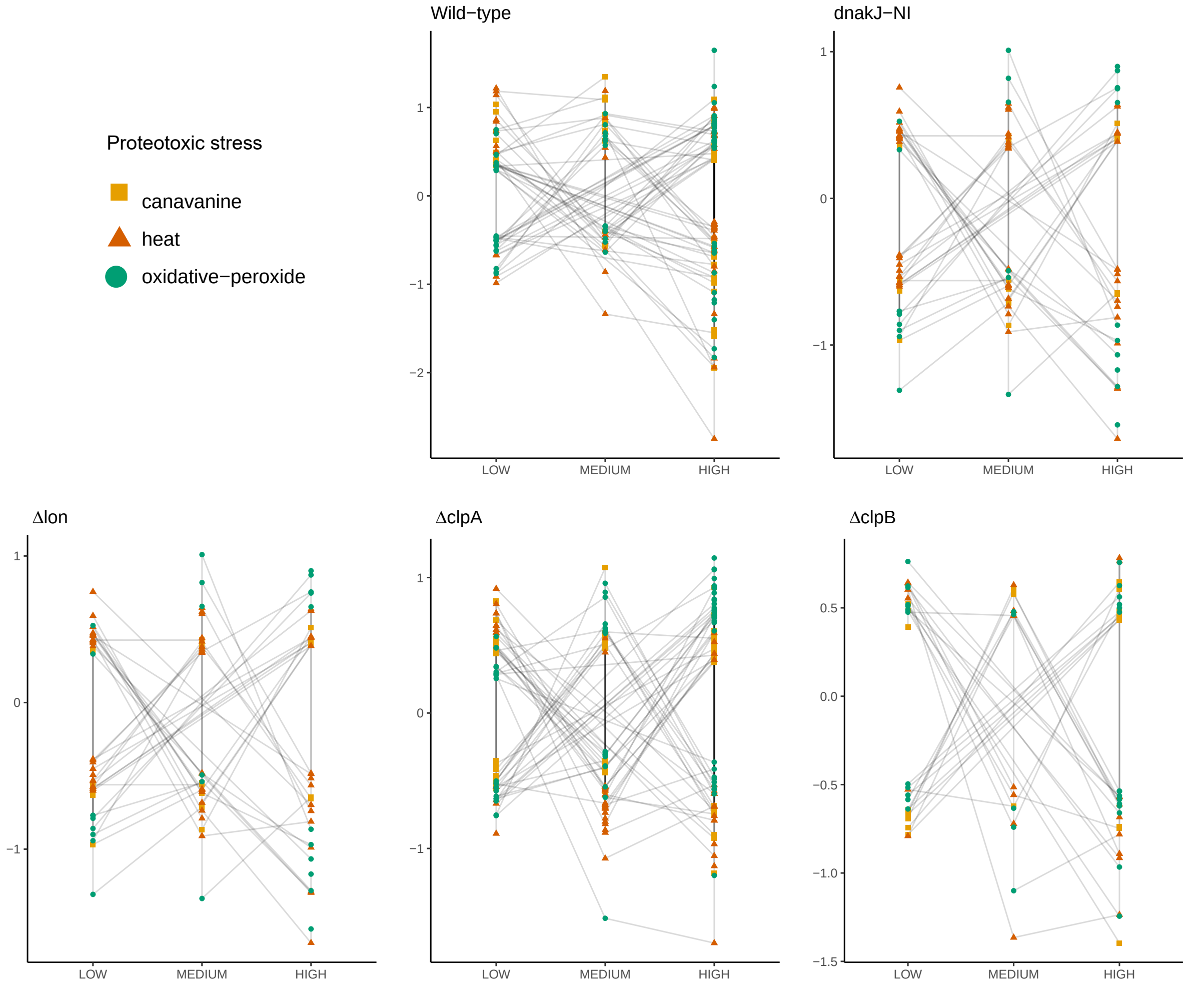


**Fig. S9. Change of fitness across different proteotoxic stress and stress levels.** The Y-axis represents coefficients from the regularized negative binomial GLM model. Points are colored based on stress. Multiple genes change their fitness between stress conditions and stress levels.

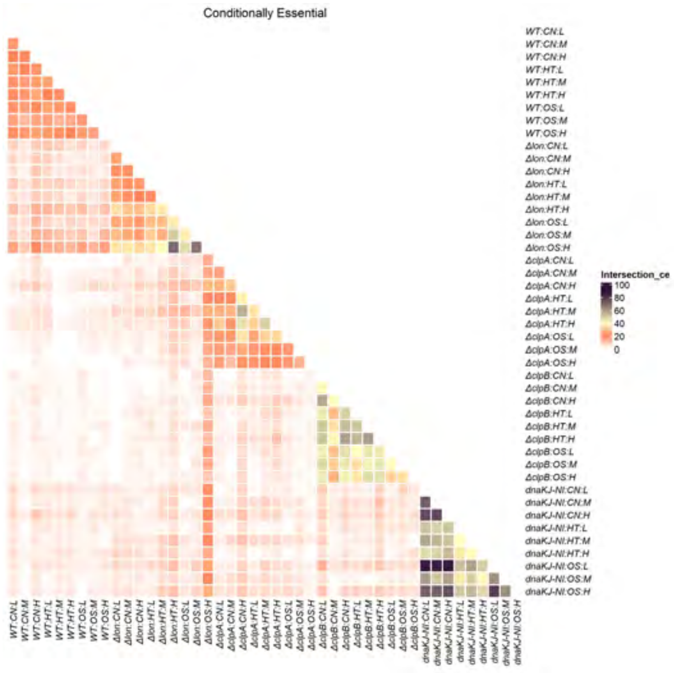


**Fig. S10. The pair-wise overlap of conditionally essential genes.** The pair-wise overlap of conditionally essential genes profiles between stress conditions. A larger overlap of essentiality profiles is seen in wild type and dnaKJ-NI compared to strains deficient in *clpA*, *lon*, or *clpB*.

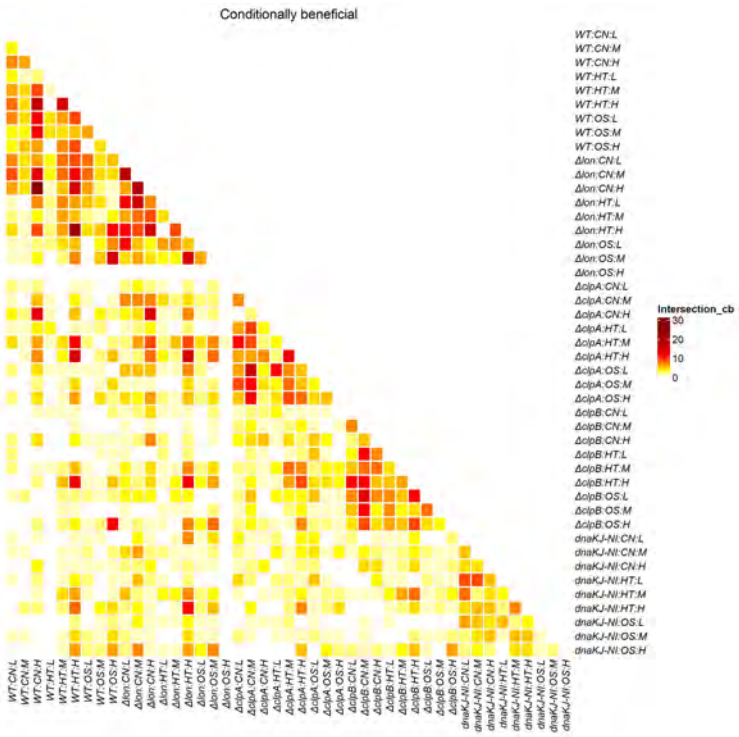


**Fig. S11. The pair-wise overlap of conditionally beneficial genes.** The pair-wise overlap of conditionally beneficial genes profiles between stress conditions. A larger overlap of essentiality profiles is seen in wild type and dnaKJ-NI compared to strains deficient in *clpA*, *lon*, or *clpB*.

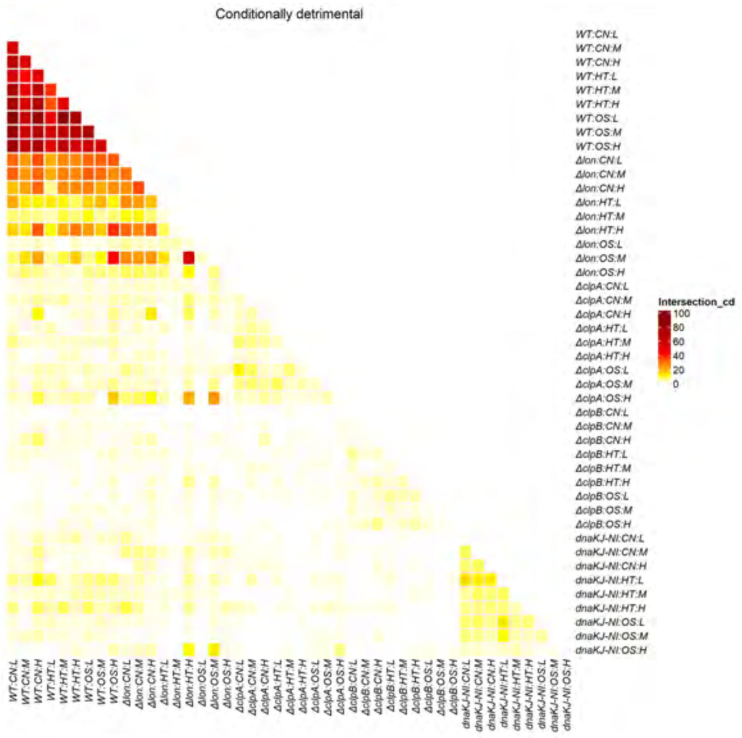


**Fig. S12. The pair-wise overlap of conditionally detrimental genes.** The pair-wise overlap of conditionally detrimental gene profiles between stress conditions. A larger overlap of essentiality profiles is seen in wild type and dnaKJ-NI compared to strains deficient in *clpA*, *lon*, or *clpB*.

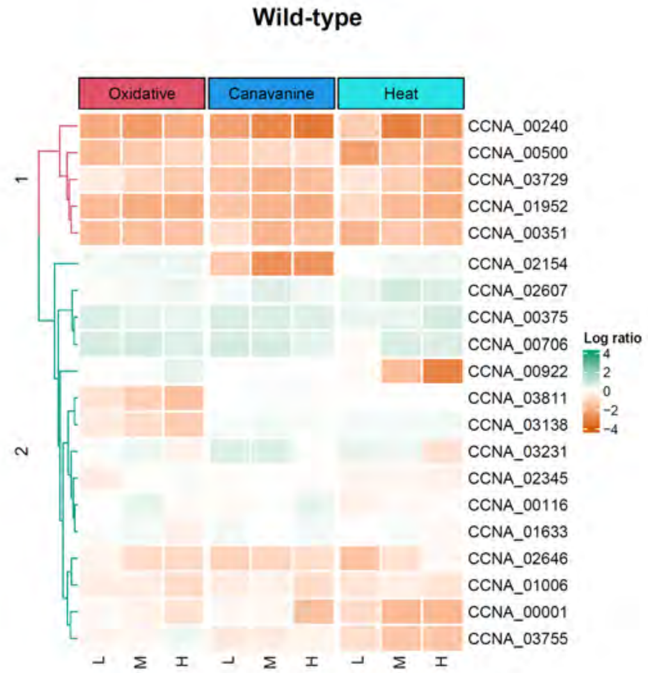


**Fig. S13. Conservation of fitness predictors in wild-type.** The model-Y knockoff framework is used to identify the most influential fitness determinants, and the fitness values of the genes across stresses are clustered in wild-type.

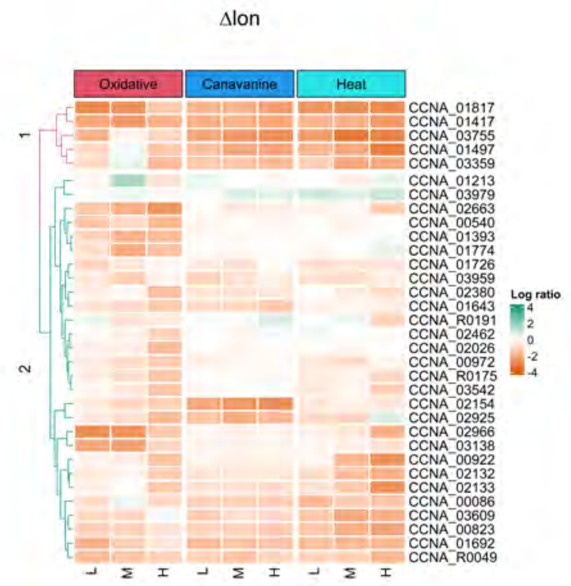


**Fig. S14. Conservation of fitness predictors in $\Delta lon$.** The model-Y knockoff framework is used to identify the most influential fitness determinants, and the fitness values of the genes across stresses are clustered in $\Delta lon$.

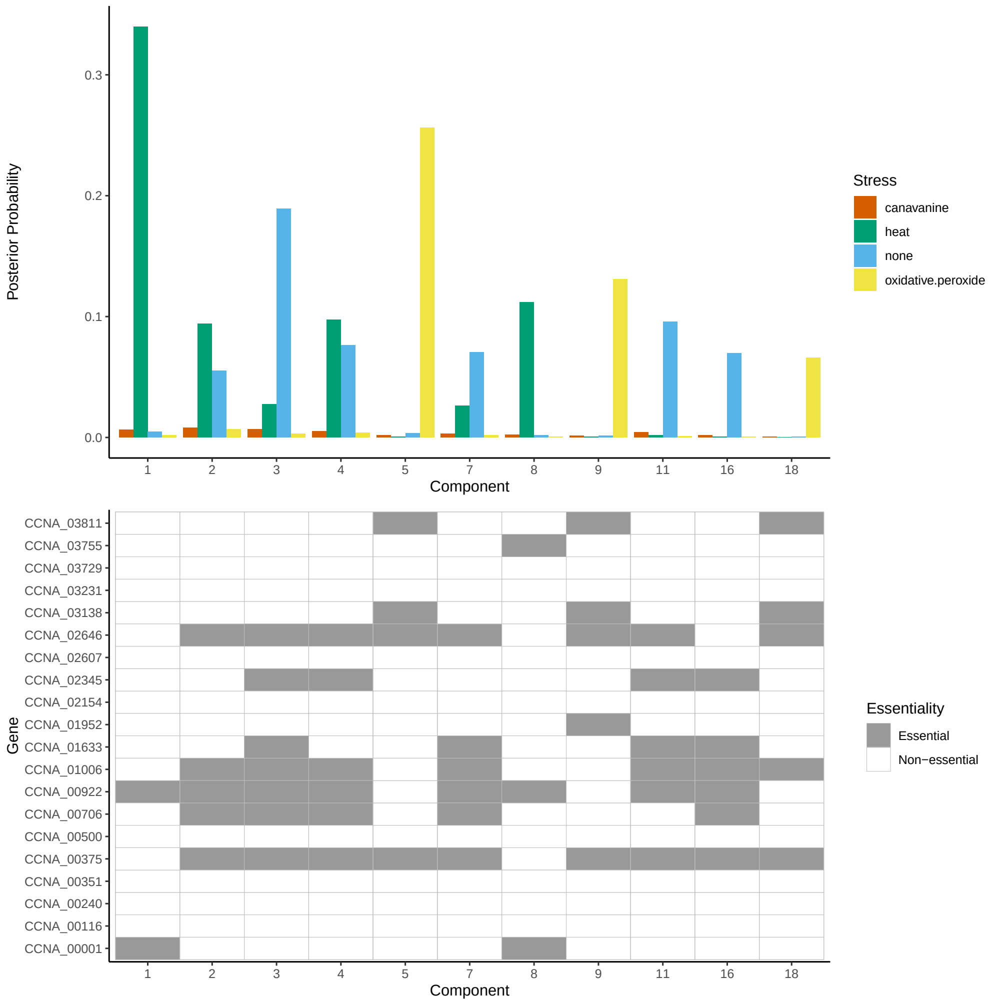


**Fig. S20. Posterior distribution plots for stresses under wild-type**. **Top** The barplot shows the distribution of stresses (none, heat, oxidative, canavanine) for each component. The majority of the posterior mass of heat stress is concentrated on component-1, whereas oxidative stress has a more posterior mass on component-5 than any other component. **Bottom** The heatmap shows the composition of each component where something is declared essential when the insertion counts decrease relatively, indicating a positive fitness contribution

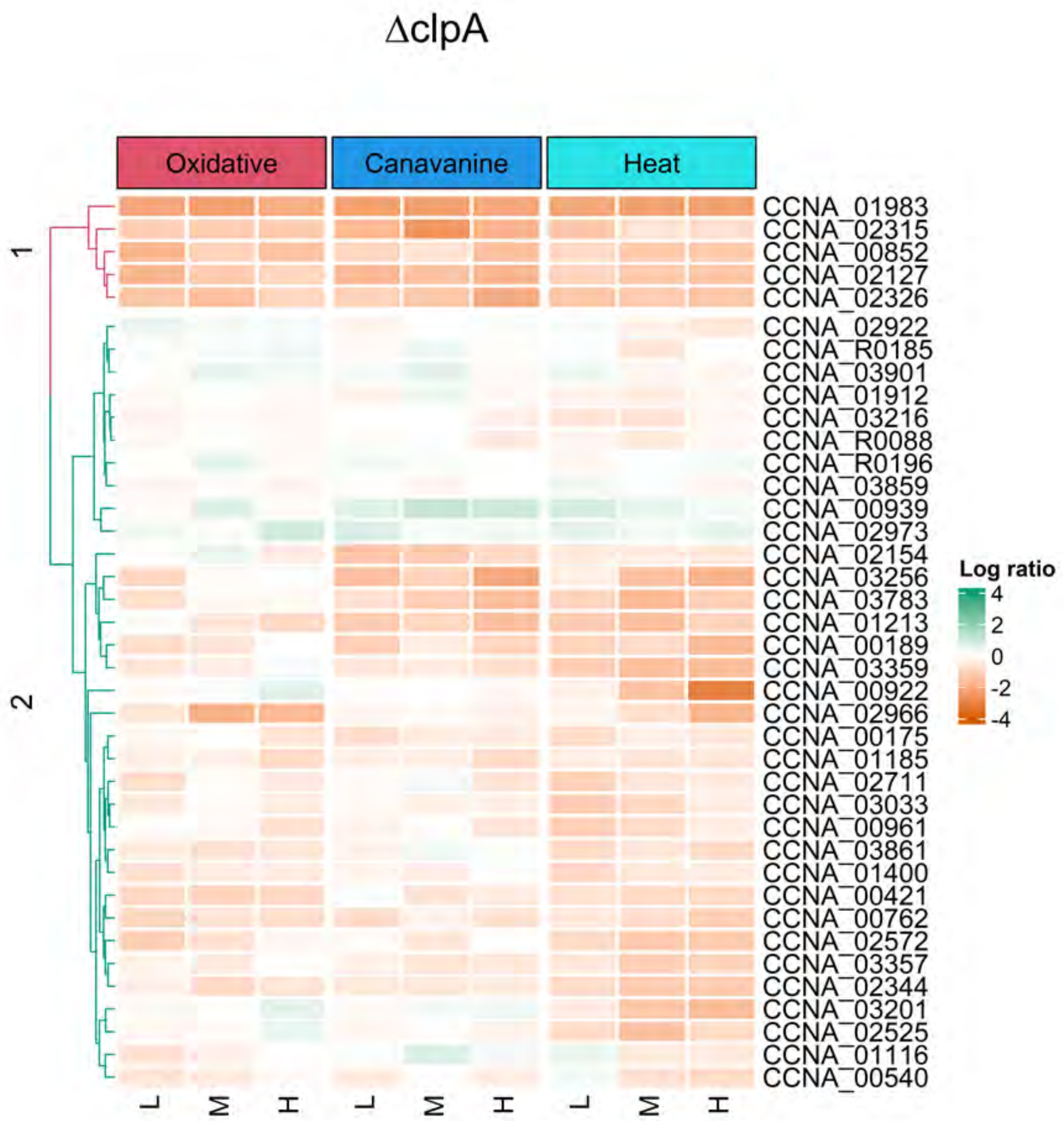


**Fig. S15. Conservation of fitness predictors in $\Delta clpA$.** The model-Y knockoff framework is used to identify the most influential fitness determinants, and the fitness values of the genes across stresses are clustered in $\Delta clpA$.

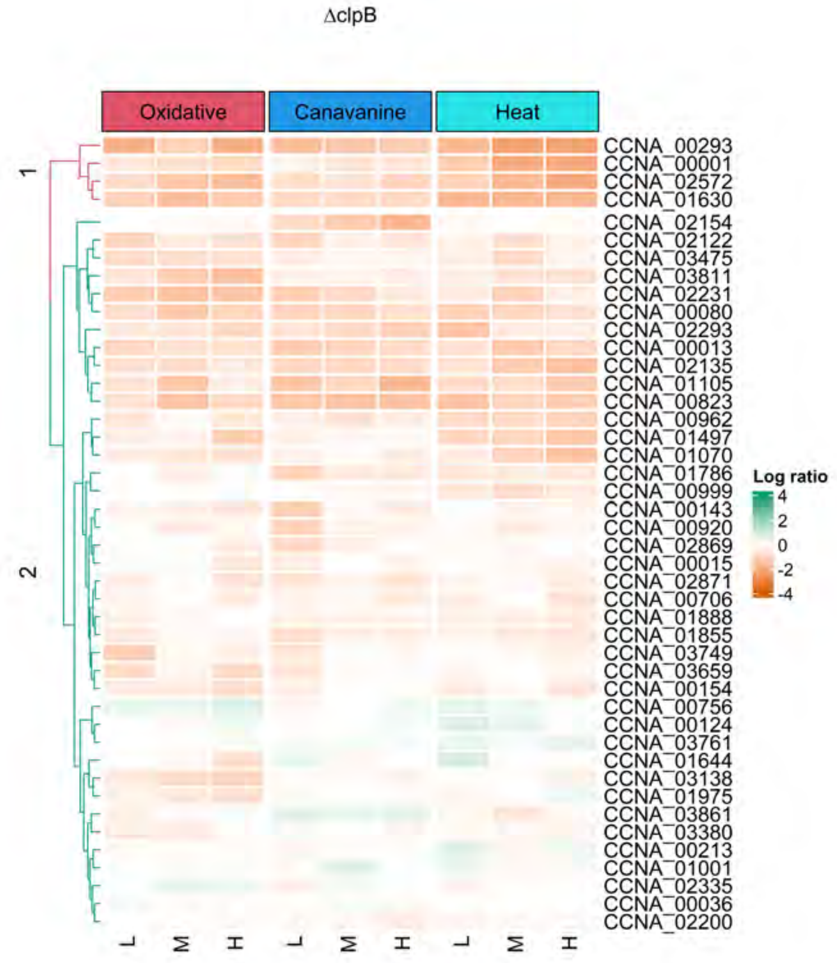


**Fig. S16. Conservation of fitness predictors in $\Delta clpB$.** The model-Y knockoff framework is used to identify the most influential fitness determinants, and the fitness values of the genes across stresses are clustered in $\Delta clpB$.

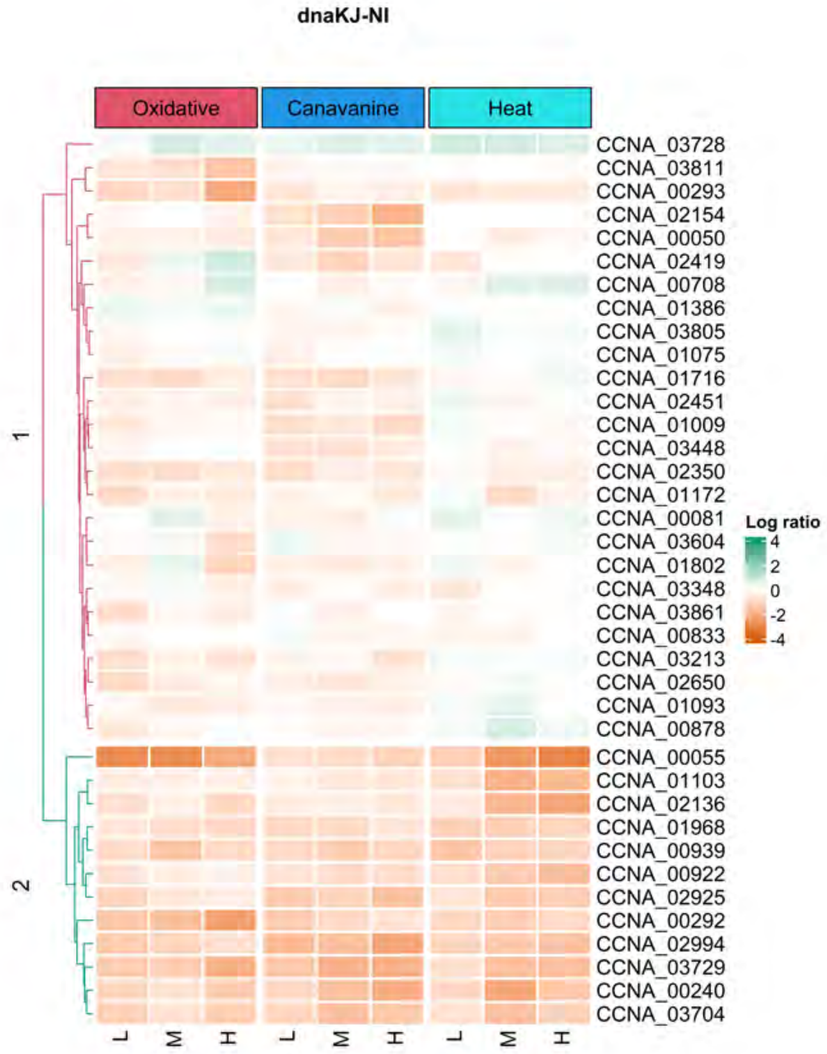


**Fig. S17. Conservation of fitness predictors in dnaKJ-NI.** The model-Y knockoff framework is used to identify the most influential fitness determinants, and the fitness values of the genes across stresses are clustered in dnaKJ-NI.

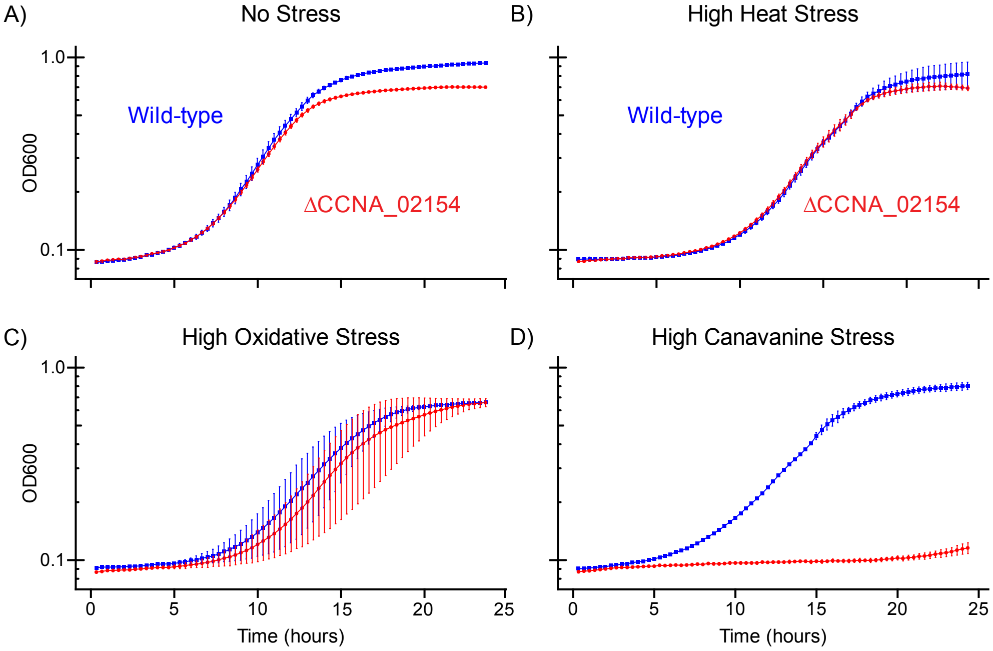


**Fig. S18.** ***In vivo*** **validation of △CCNA_02154 phenotype.** OD600 growth curves generated from wild-type and △CCNA_02154 strains subjected to heat, oxidative, and canavanine stresses. Growth curves showing **A)** strains grown in no stress condition, **B)** strains heat-shocked at 43.8C for 45 minutes and grown in 30C for 24 hours, **C)** strains grown in 0.1mM hydrogen peroxide, and **D)** strains grown in 100ug/ml l-canavanine.

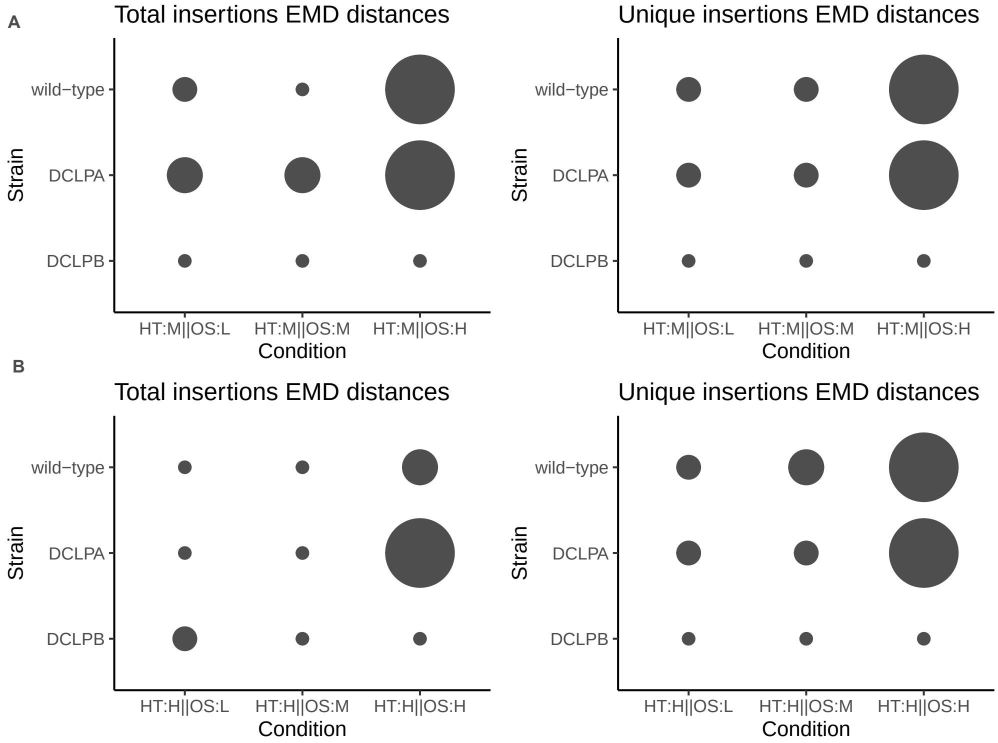


**Fig. S19. Stress-Induced Fitness Effects for medium and high heat.**. Using tnseq data, we assess the fitness variations under medium (A), high (B) heat, and oxidative peroxide stress in WT, $\Delta clpA$, and $\Delta clpB$ strains. Differences are quantified using EMD (Earth Mover's Distance) based on both total and unique insertion counts. Bubble size corresponds to the magnitude of the EMD distances

.

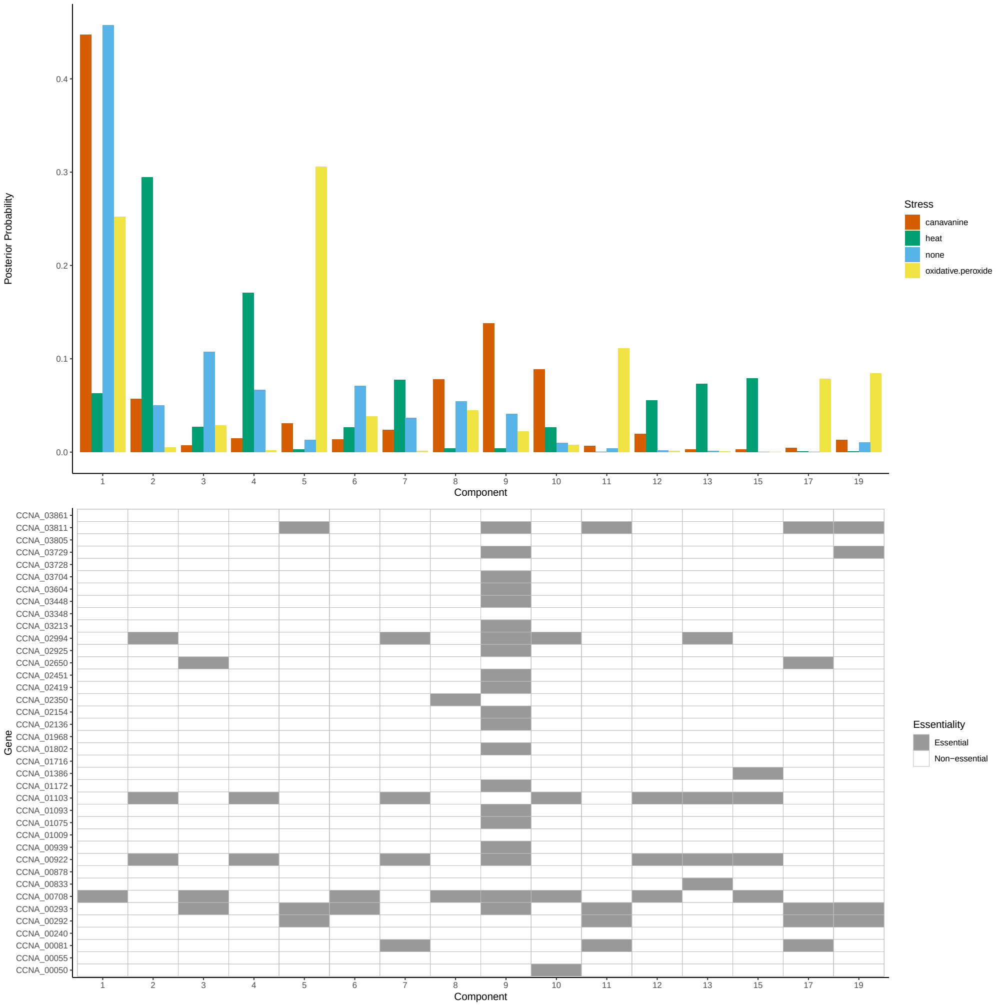


**Fig. S21. Posterior distribution plots for stresses under dnaKJ-NI**. **Top** The barplot shows the distribution of stresses (none, heat, oxidative, canavanine) for each component. Component-1 has similar mass concentrations among the different stress types, whereas heat shows concentration in component-2 and oxidative shows concentration in component-5. **Bottom** The heatmap shows the composition of each component where something is declared essential when the insertion counts decrease relatively, indicating a positive fitness contribution.

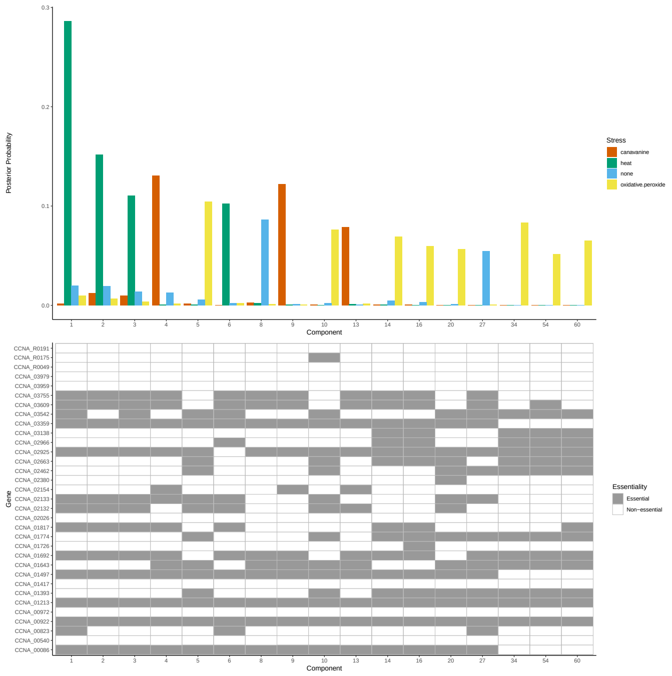


**Fig. S22. Posterior distribution plots for stresses under $\Delta lon$. Top** The barplot shows the distribution of stresses (none, heat, oxidative, canavanine) for each component. **Bottom** The heatmap shows the composition of each component where something is declared essential when the insertion counts decrease relatively, indicating a positive fitness contribution.

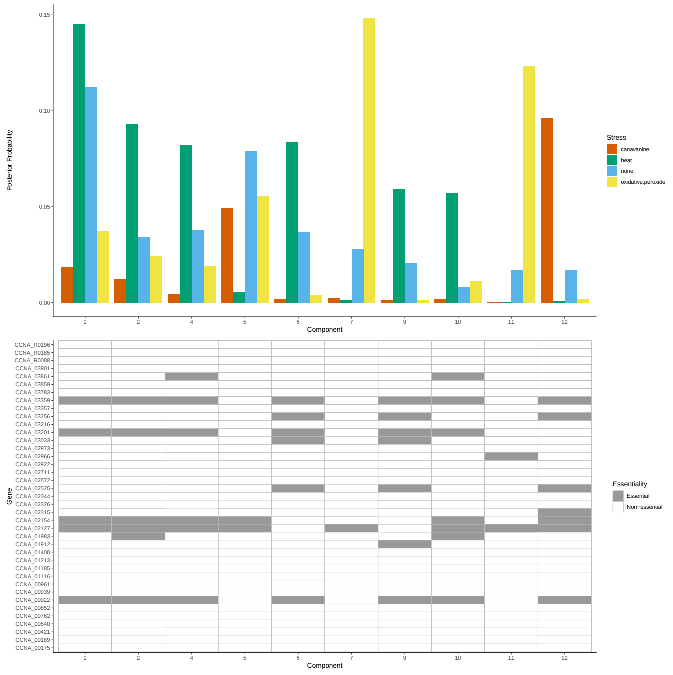


**Fig. S23. Posterior distribution plots for stresses under $\Delta clpA$. Top** The barplot shows the distribution of stresses (none, heat, oxidative, canavanine) for each component. **Bottom** The heatmap shows the composition of each component where something is declared essential when the insertion counts decrease relatively, indicating a positive fitness contribution.

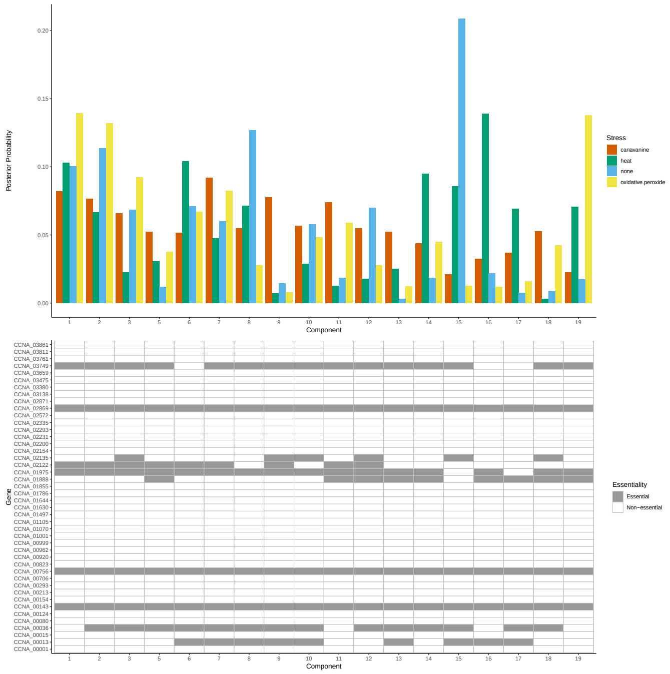


**Fig. S24. Posterior distribution plots for stresses under $\Delta clpB$. Top** The barplot shows the distribution of stresses (none, heat, oxidative, canavanine) for each component. **Bottom** The heatmap shows the composition of each component where something is declared essential when the insertion counts decrease relatively, indicating a positive fitness contribution.

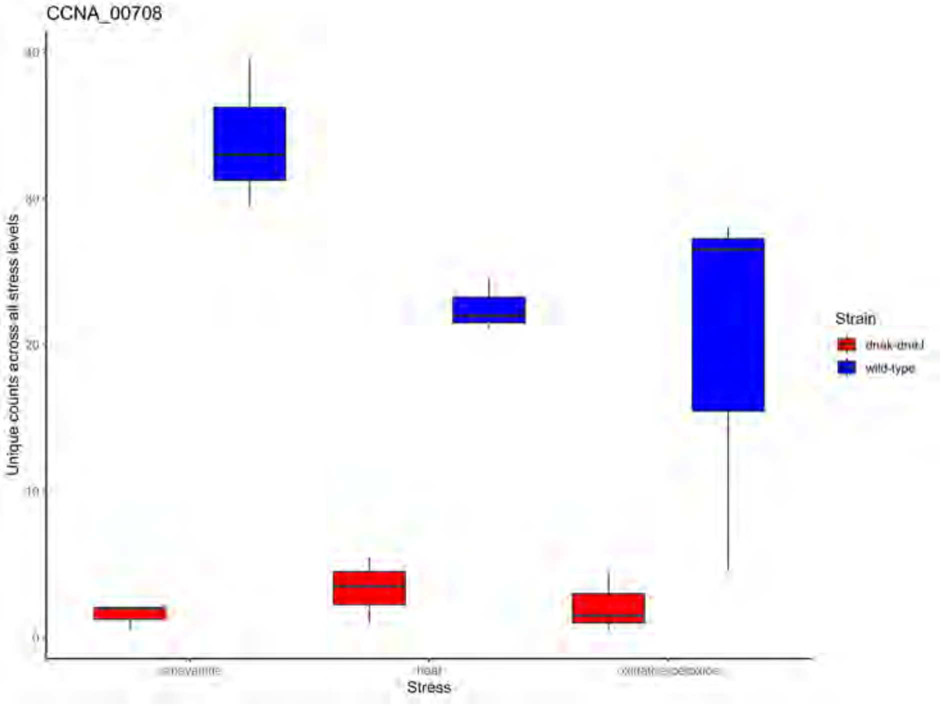


**Fig. S25. Unique count distribution of CCNA_00708 for different stresses under wild-type and dnaKJ-NI**. The y-axis denotes the median unique insertion counts of canavanine heat and oxidative stresses across all stress levels for the wild-type and dnaKJ-NI.

| Strain | Avg library size |
|---|---|
| $\Delta clpA$ | 53129.54 |
| $\Delta clpB$ | 30556 |
| dnaKJ-NI | 37059.25 |
| $\Delta lon$ | 146410.36 |
| Wild-type | 186481.55 |

| Condition | Avg library size |
|---|---|
| Canavanine | 91008.35 |
| No stress | 100604.5 |
| Heat | 90436.6 |
| Oxidative | 84013.42 |

| Batch | Avg library size |
|---|---|
| 1 | 138705.3 |
| 2 | 115869.9 |
| 3 | 50190.66 |
| 4 | 87492.5 |
| 5 | 31657.8 |

**Table S2. The average library size of unique insertions across different strains, conditions, and batches.**

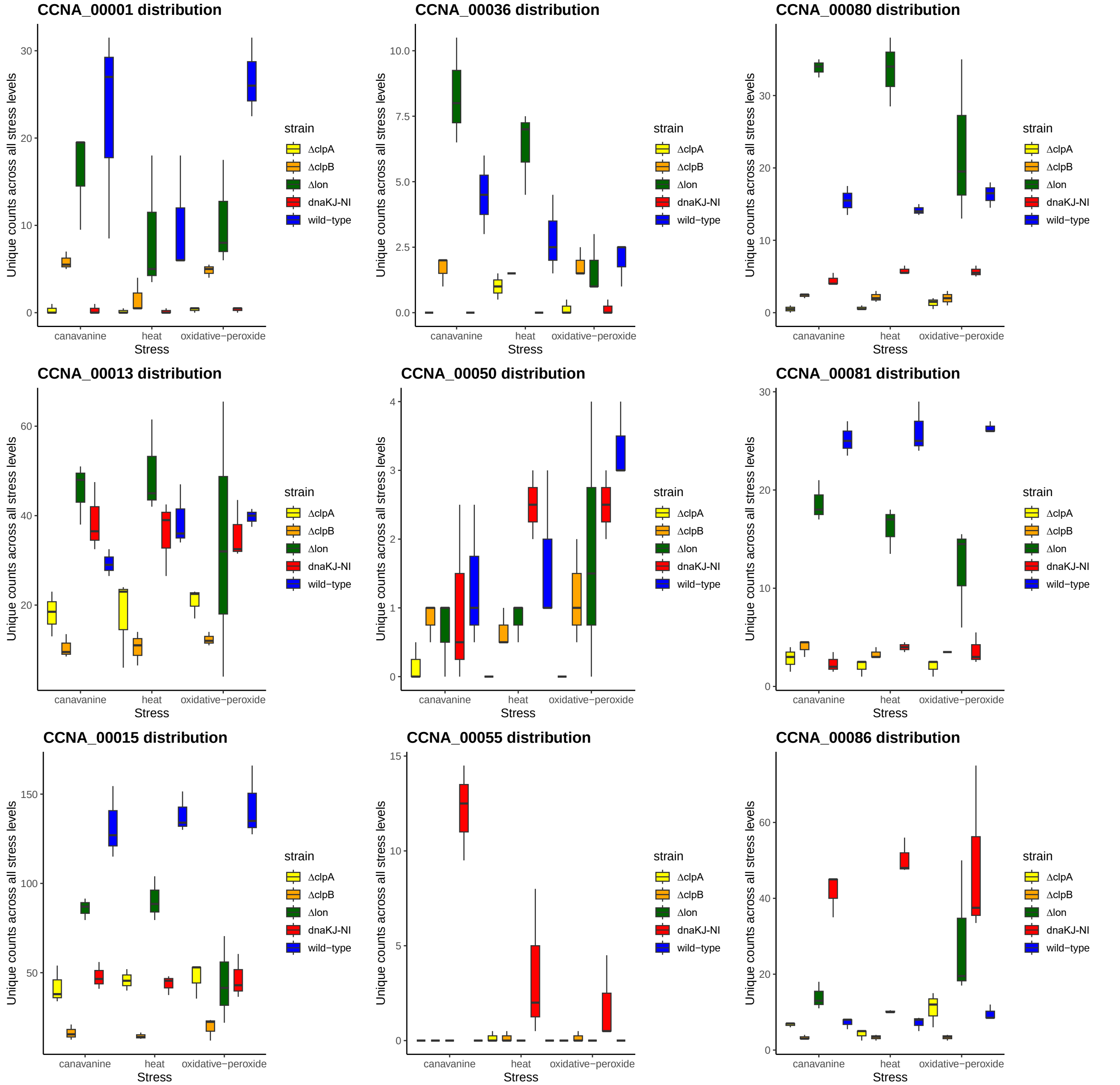


**Fig. S26. The unique count insertion distributions of fitness determinants** The Y-axis shows the unique count insertion distributions of genes that are predicted as fitness determinants by the model-Y knockoffs framework. This plot shows the distribution of CCNA_00001, CCNA_00036, CCNA_00080, CCNA_00013, CCNA_00050, CCNA_00081, CCNA_00015, CCNA_00055, CCNA_00086.

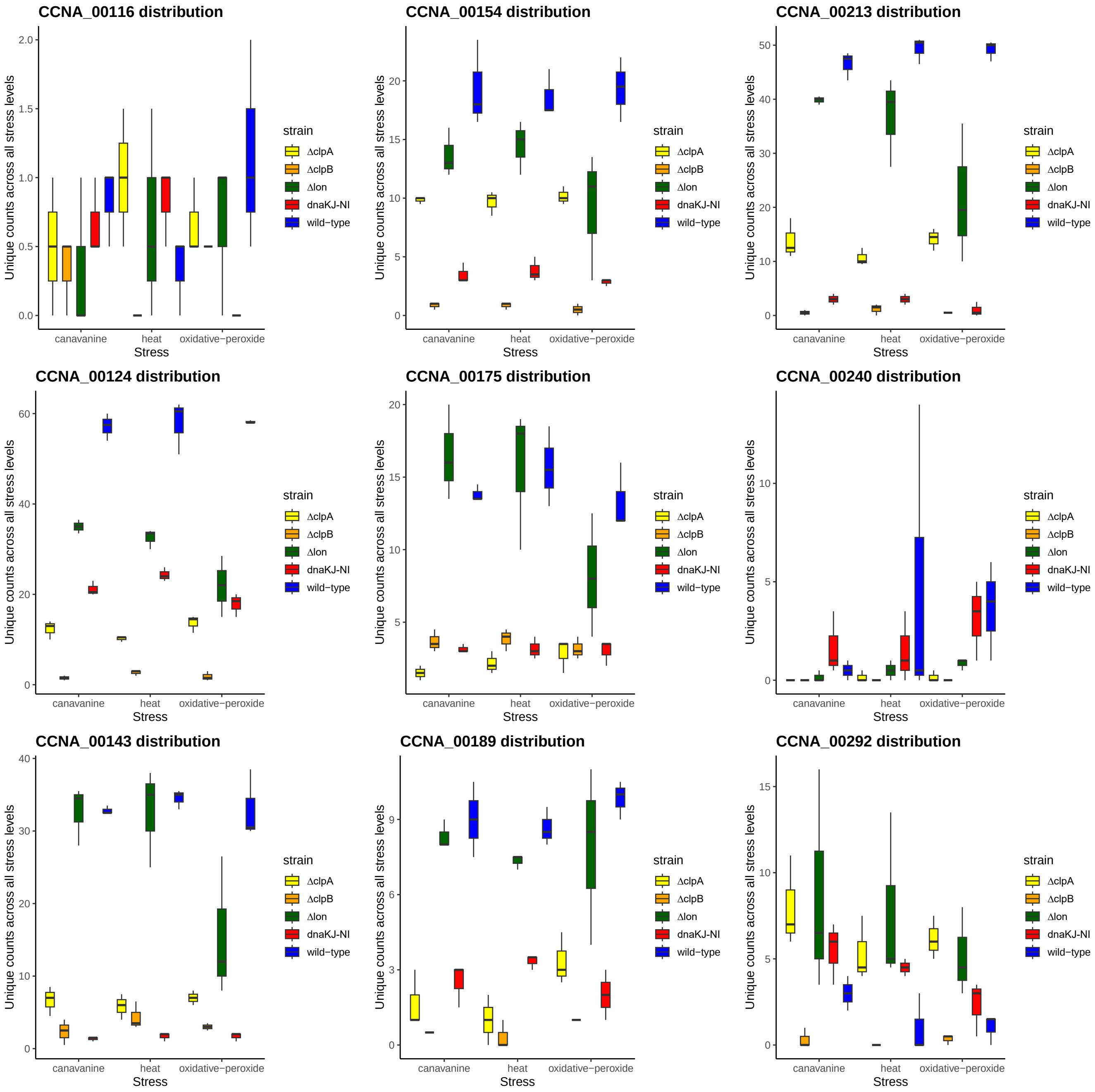


**Fig. S27. The unique count insertion distributions of fitness determinants** The Y-axis shows the unique count insertion distributions of genes that are predicted as fitness determinants by the model-Y knockoffs framework. This plot shows the distribution of CCNA_00116, CCNA_00154, CCNA_00213, CCNA_00124, CCNA_00175, CCNA_00240, CCNA_00143, CCNA_00189, CCNA_00292.

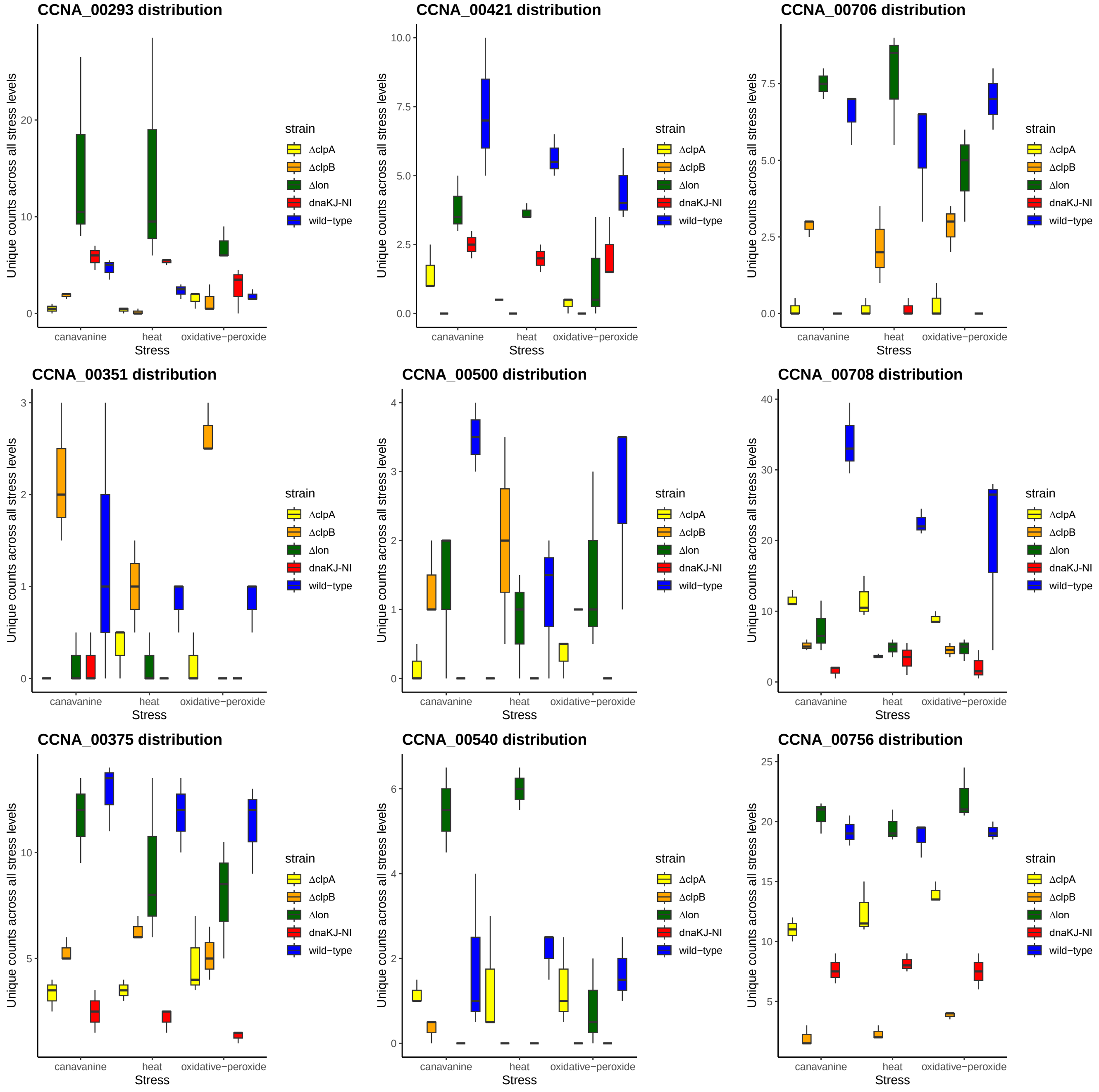


**Fig. S28. The unique count insertion distributions of fitness determinants** The Y-axis shows the unique count insertion distributions of genes that are predicted as fitness determinants by the model-Y knockoffs framework. This plot shows the distribution of CCNA_00293, CCNA_00421, CCNA_00706, CCNA_00351, CCNA_00500, CCNA_00708, CCNA_00375, CCNA_00540, CCNA_00756.

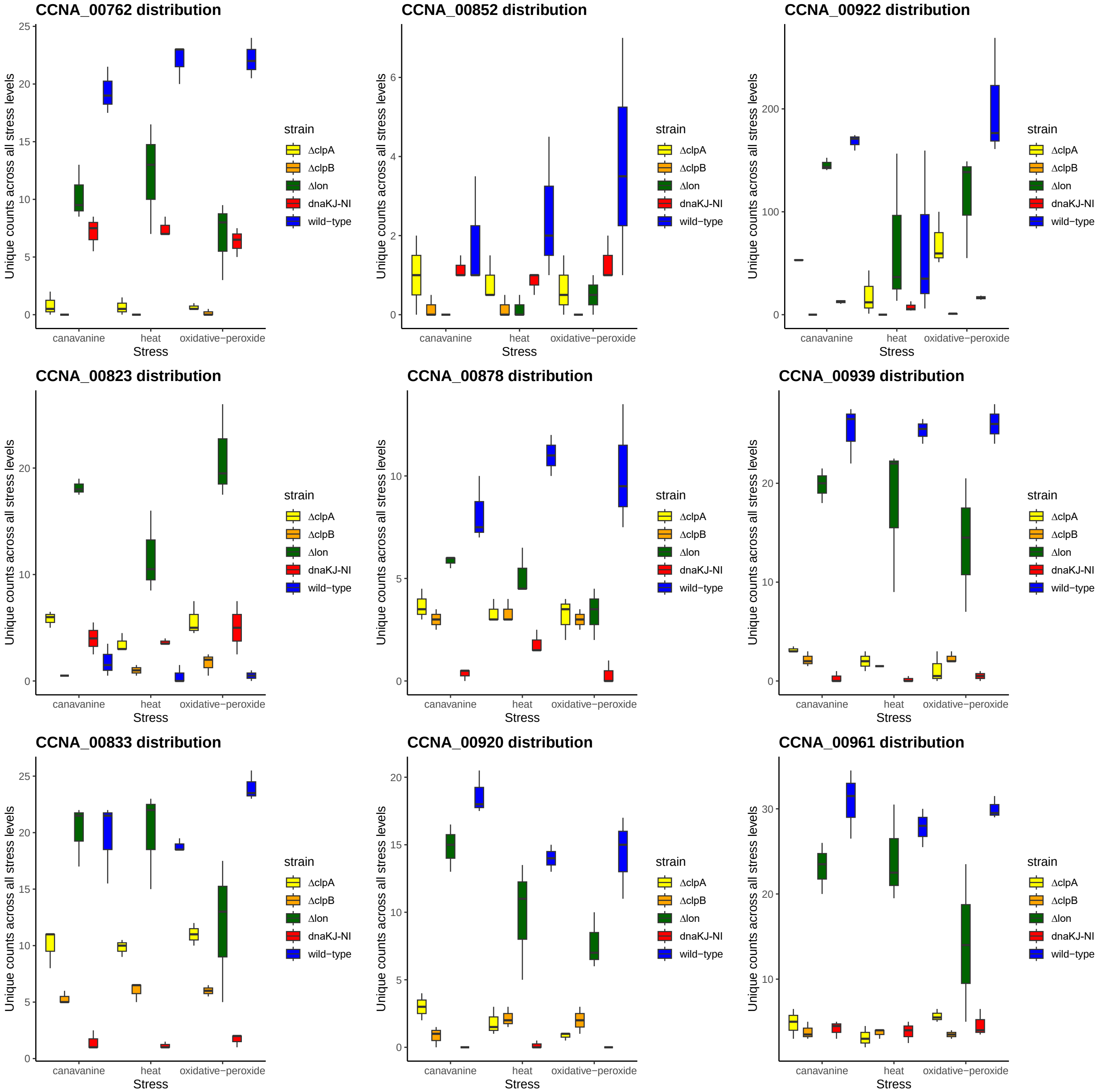


**Fig. S29. The unique count insertion distributions of fitness determinants** The Y-axis shows the unique count insertion distributions of genes that are predicted as fitness determinants by the model-Y knockoffs framework. This plot shows the distribution of CCNA_00762, CCNA_00852, CCNA_00922, CCNA_00823, CCNA_00878, CCNA_00939, CCNA_00833, CCNA_00920, CCNA_00961.

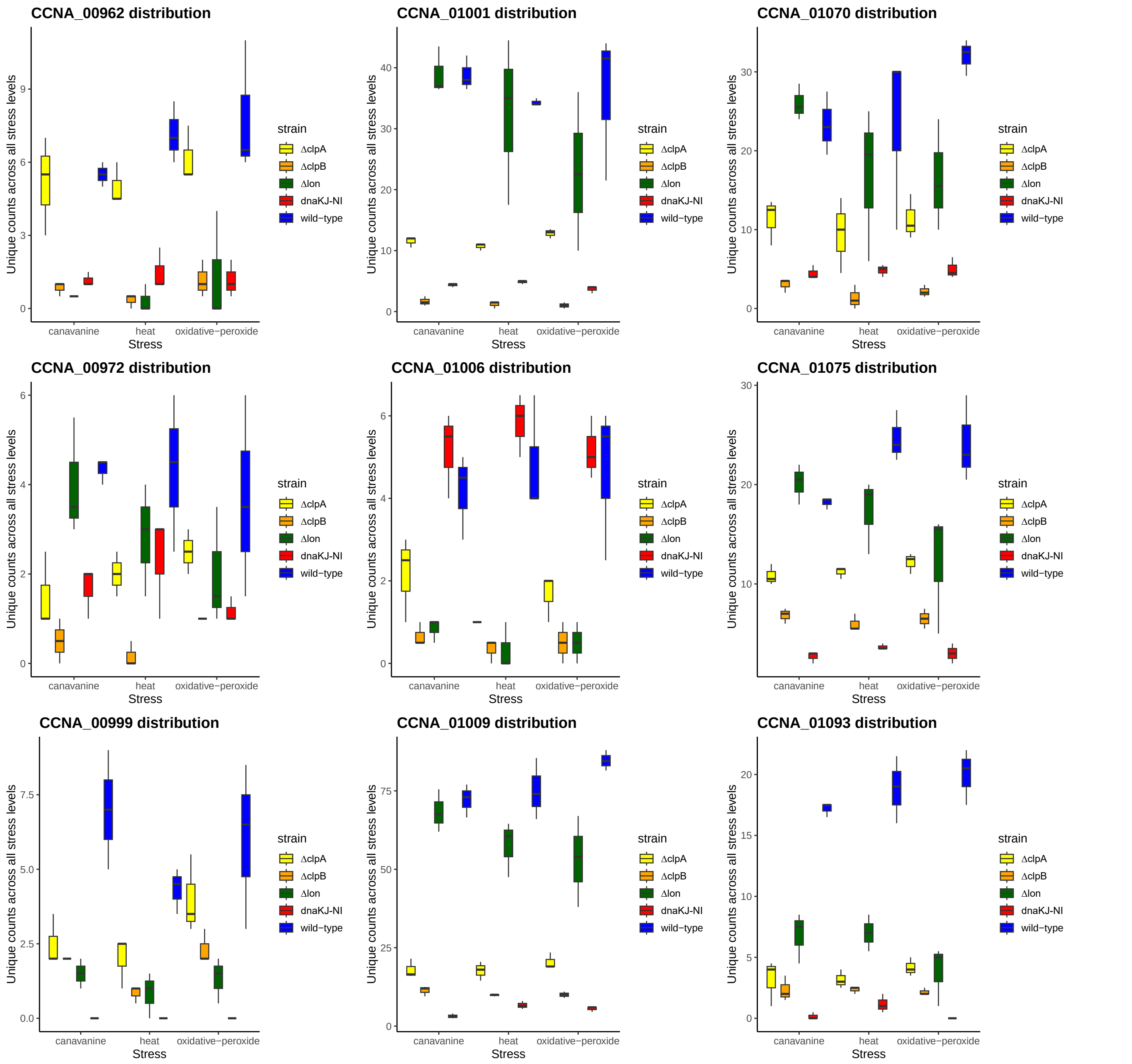


**Fig. S30. The unique count insertion distributions of fitness determinants** The Y-axis shows the unique count insertion distributions of genes that are predicted as fitness determinants by the model-Y knockoffs framework. This plot shows the distribution of CCNA_00962, CCNA_01001, CCNA_01070, CCNA_00972, CCNA_01006, CCNA_01075, CCNA_00999, CCNA_01009, CCNA_01093.

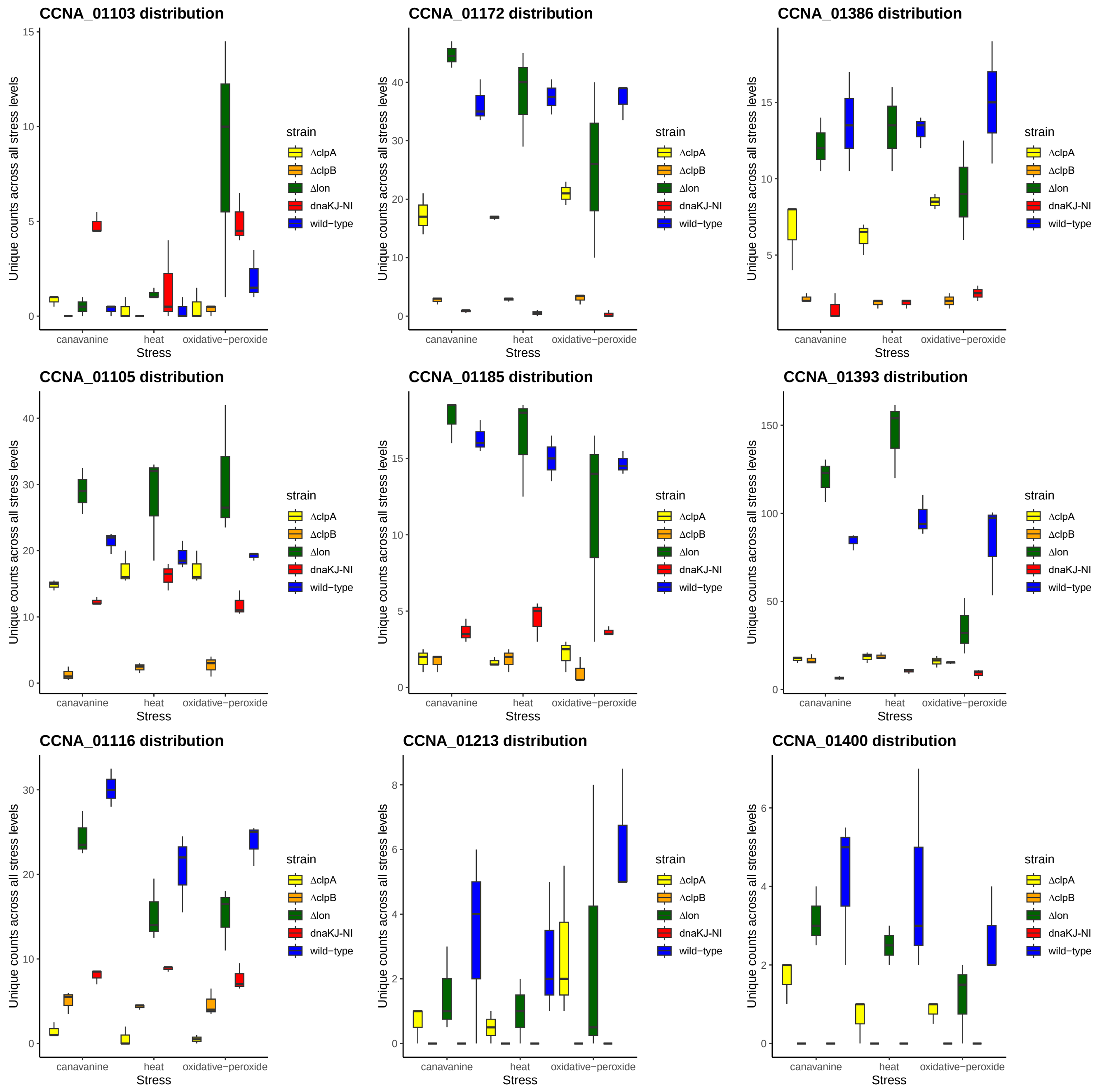


**Fig. S31. The unique count insertion distributions of fitness determinants** The Y-axis shows the unique count insertion distributions of genes that are predicted as fitness determinants by the model-Y knockoffs framework. This plot shows the distribution of CCNA_01103, CCNA_01172, CCNA_01386, CCNA_01105, CCNA_01185, CCNA_01393, CCNA_01116, CCNA_01213, CCNA_01400.

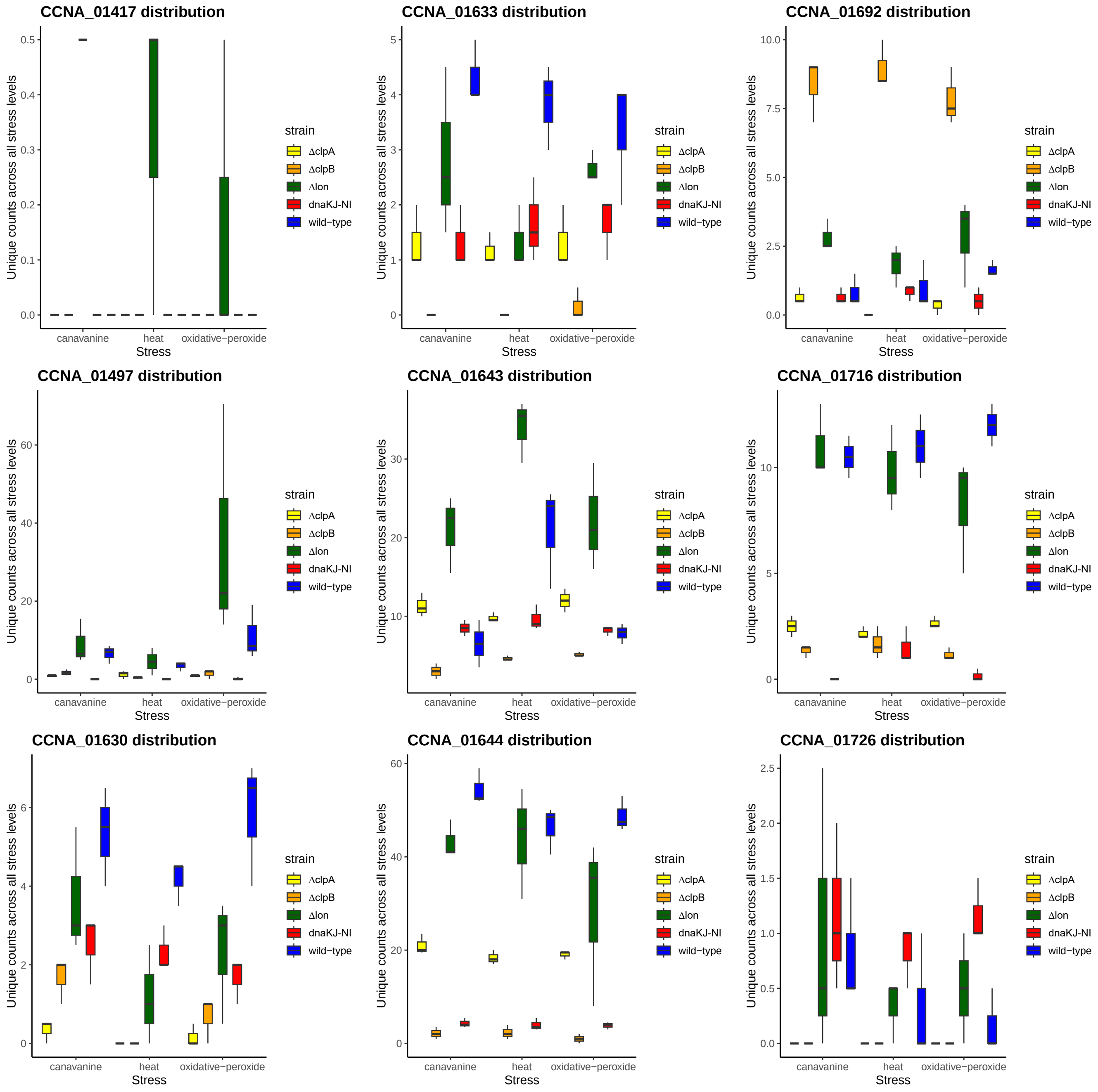


**Fig. S32. The unique count insertion distributions of fitness determinants** The Y-axis shows the unique count insertion distributions of genes that are predicted as fitness determinants by the model-Y knockoffs framework. This plot shows the distribution of CCNA_01417, CCNA_01633, CCNA_01692, CCNA_01497, CCNA_01643, CCNA_01716, CCNA_01630, CCNA_01644, CCNA_01726.

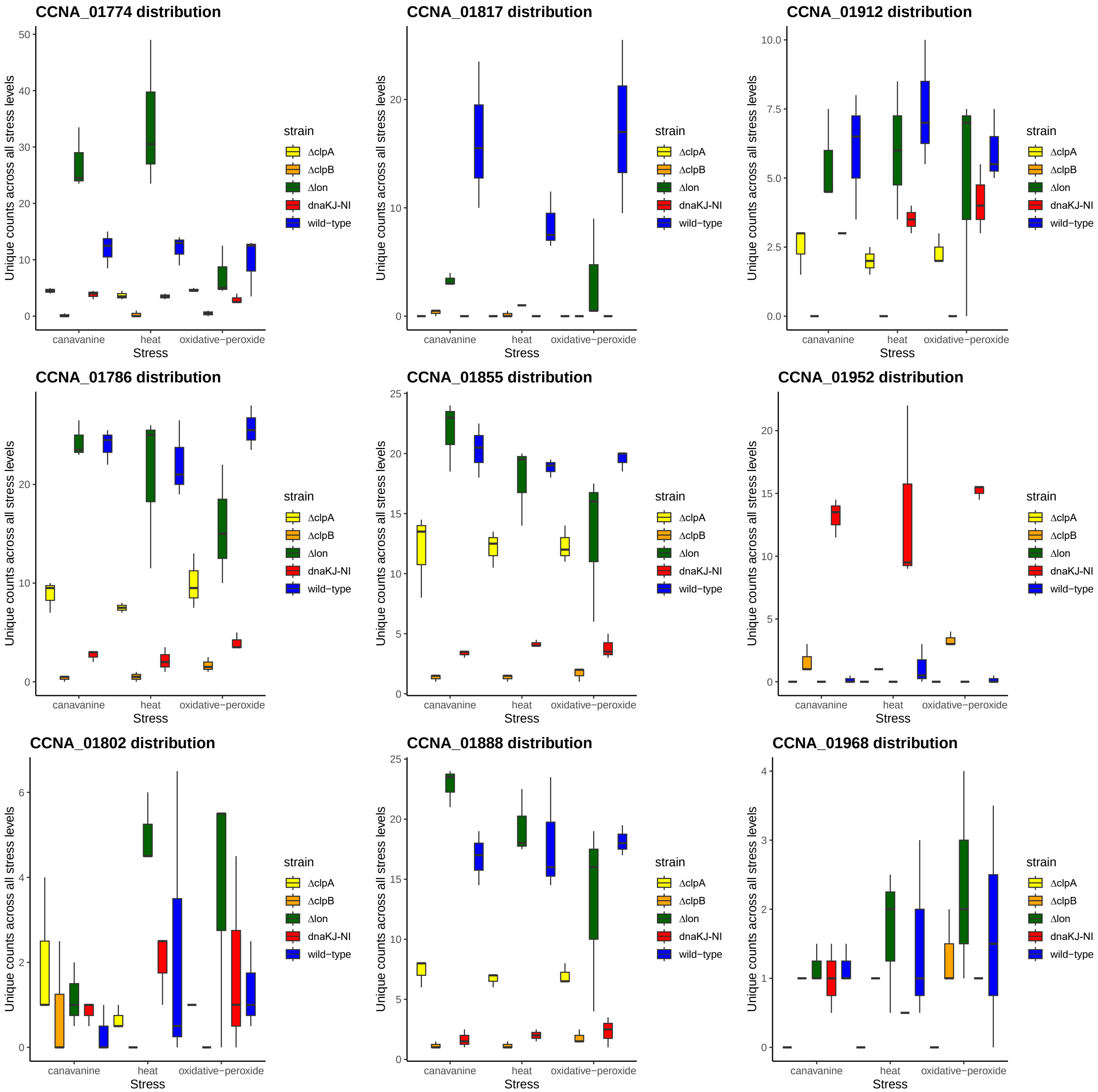


**Fig. S33. The unique count insertion distributions of fitness determinants** The Y-axis shows the unique count insertion distributions of genes that are predicted as fitness determinants by the model-Y knockoffs framework. This plot shows the distribution of CCNA_01774, CCNA_01817, CCNA_01912, CCNA_01786, CCNA_01855, CCNA_01952, CCNA_01802, CCNA_01888, CCNA_01968.

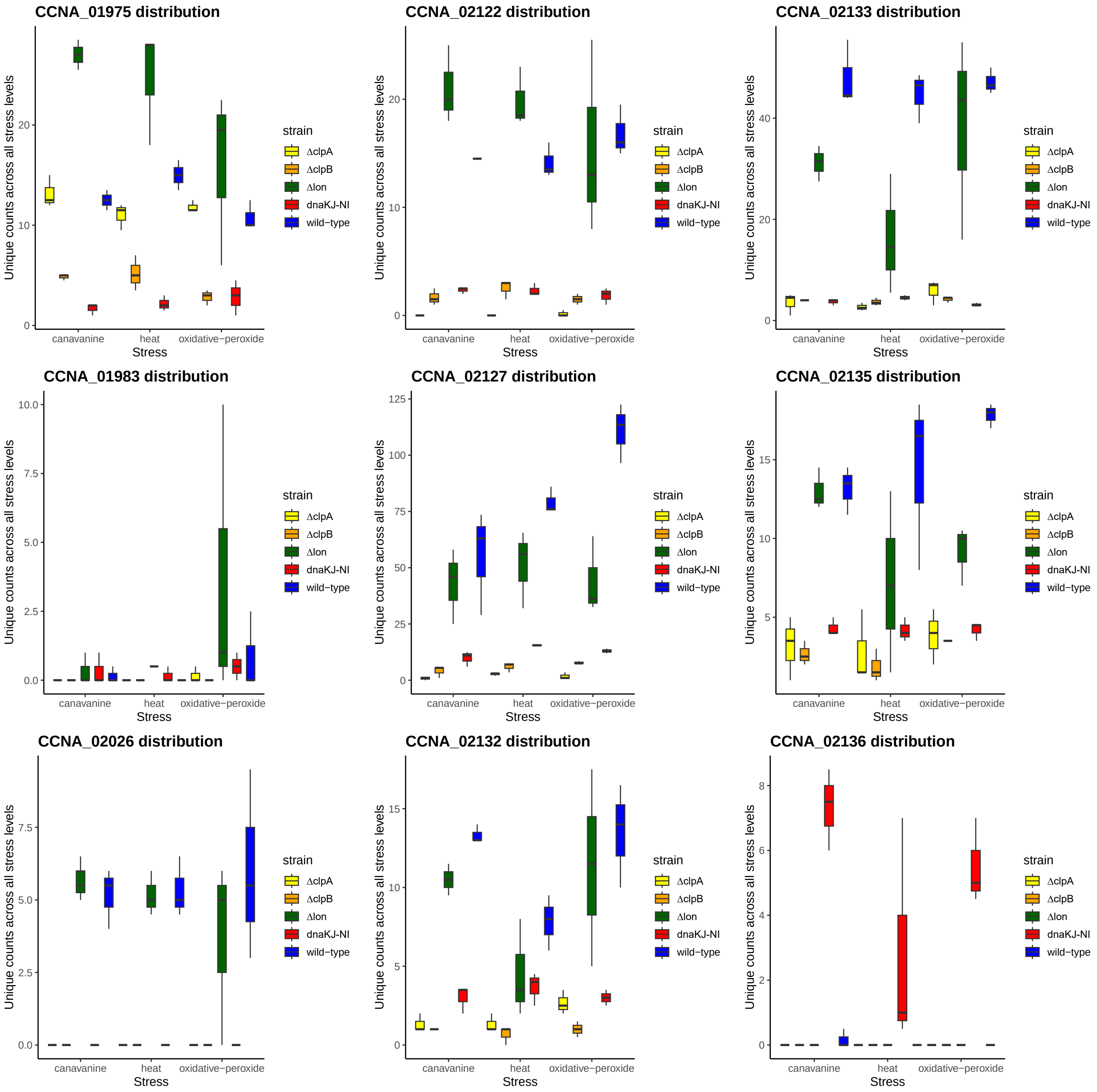


**Fig. S34. The unique count insertion distributions of fitness determinants** The Y-axis shows the unique count insertion distributions of genes that are predicted as fitness determinants by the model-Y knockoffs framework. This plot shows the distribution of CCNA_01975, CCNA_02122, CCNA_02133, CCNA_01983, CCNA_02127, CCNA_02135, CCNA_02026, CCNA_02132, CCNA_02136.

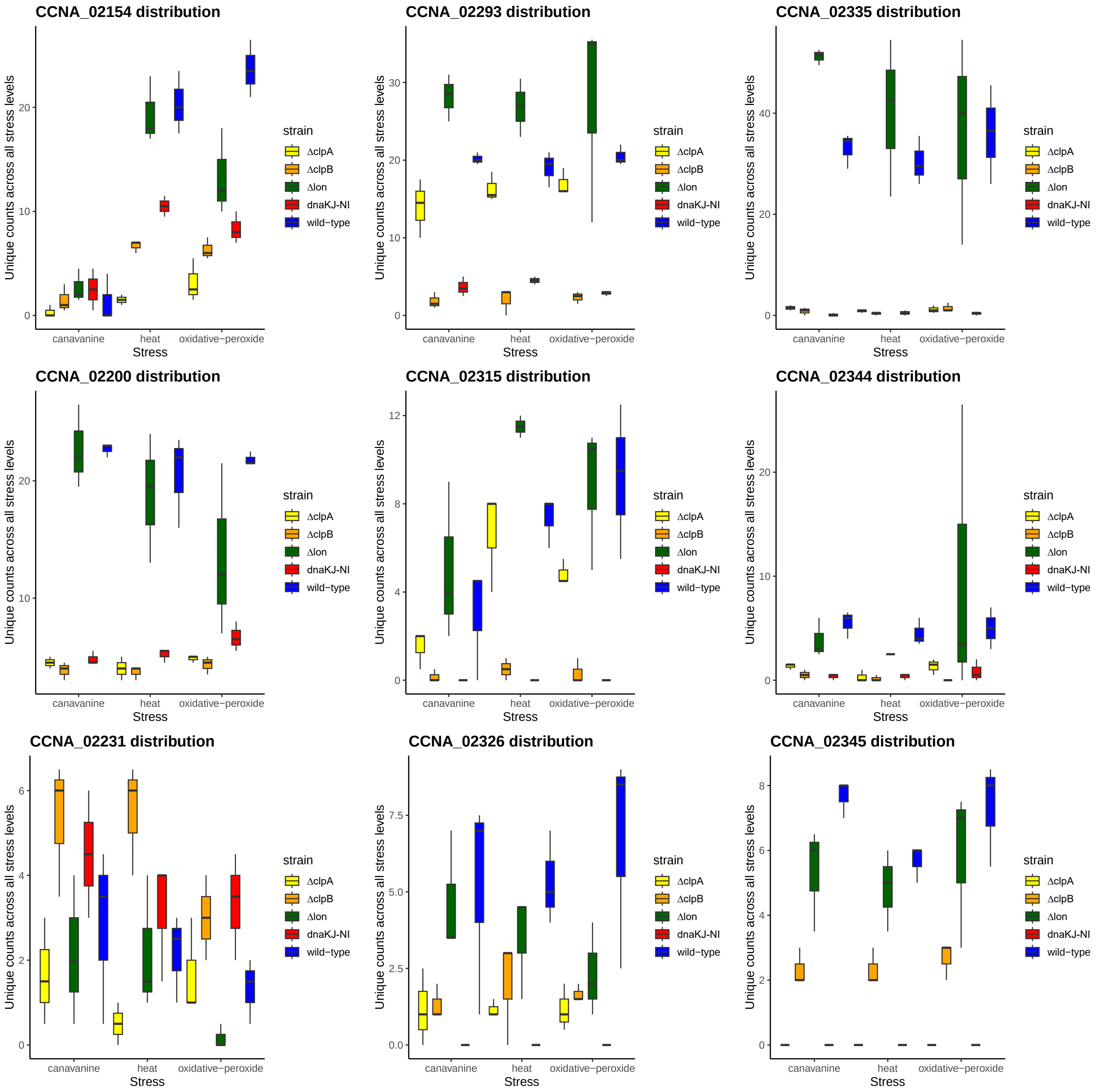


**Fig. S35. The unique count insertion distributions of fitness determinants** The Y-axis shows the unique count insertion distributions of genes that are predicted as fitness determinants by the model-Y knockoffs framework. This plot shows the distribution of CCNA_02154, CCNA_02293, CCNA_02335, CCNA_02200, CCNA_02315, CCNA_02344, CCNA_02231, CCNA_02326, CCNA_02345.

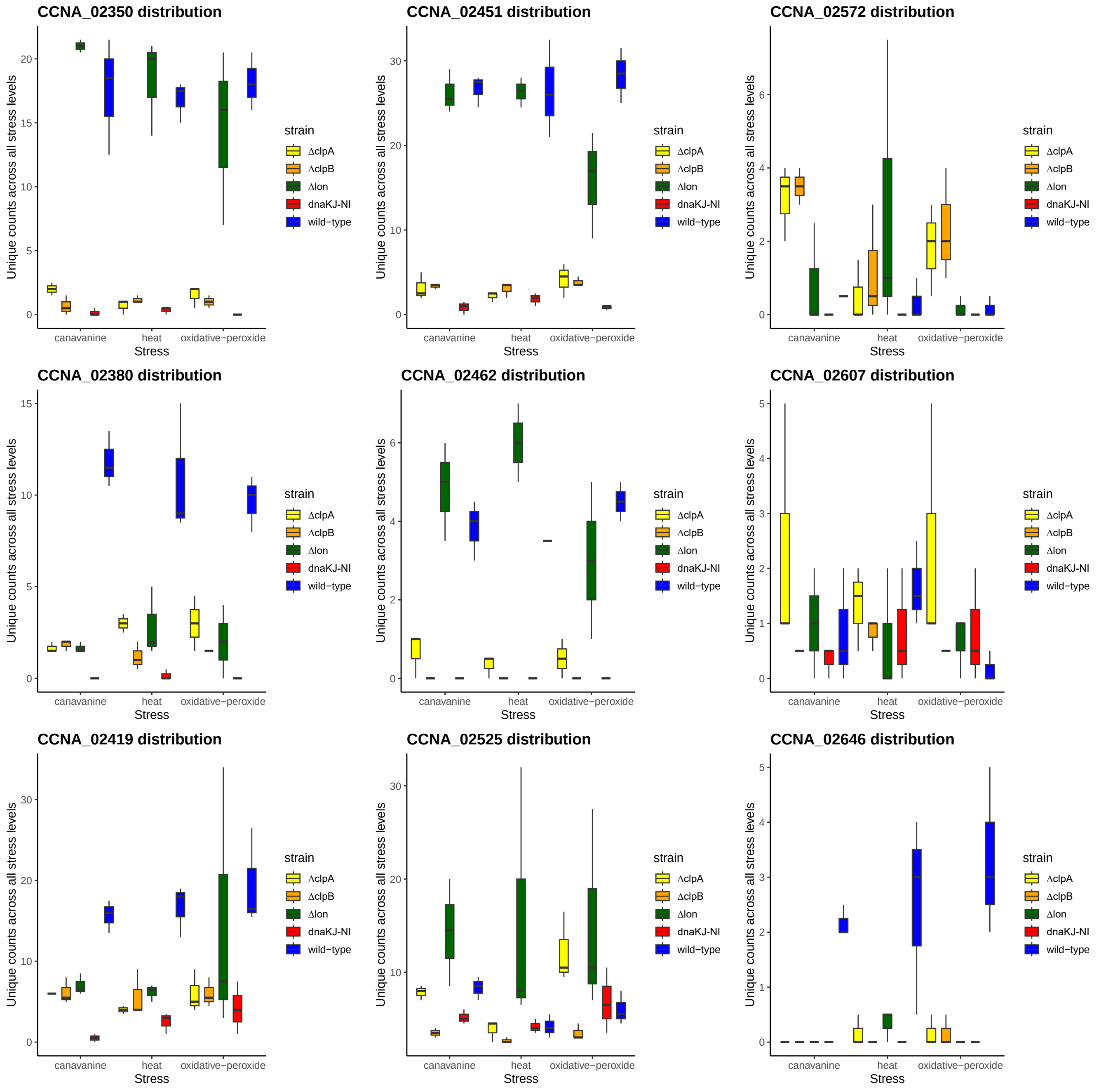


**Fig. S36. The unique count insertion distributions of fitness determinants** The Y-axis shows the unique count insertion distributions of genes that are predicted as fitness determinants by the model-Y knockoffs framework. This plot shows the distribution of CCNA_02350, CCNA_02451, CCNA_02572, CCNA_02380, CCNA_02462, CCNA_02607, CCNA_02419, CCNA_02525, CCNA_02646.

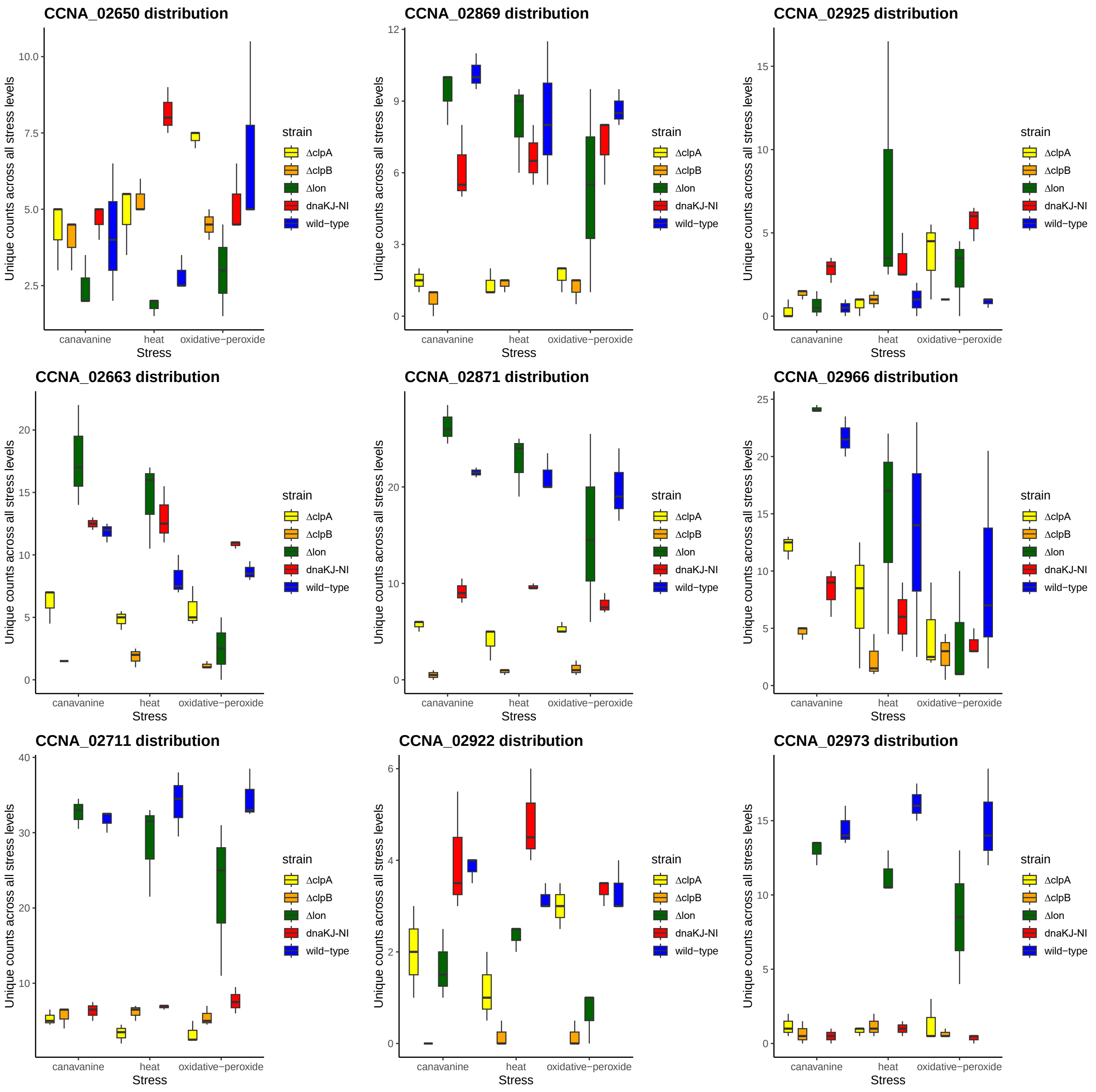


**Fig. S37. The unique count insertion distributions of fitness determinants** The Y-axis shows the unique count insertion distributions of genes that are predicted as fitness determinants by the model-Y knockoffs framework. This plot shows the distribution of CCNA_02650, CCNA_02869, CCNA_02925, CCNA_02663, CCNA_02871, CCNA_02966, CCNA_02711, CCNA_02922, CCNA_02973.

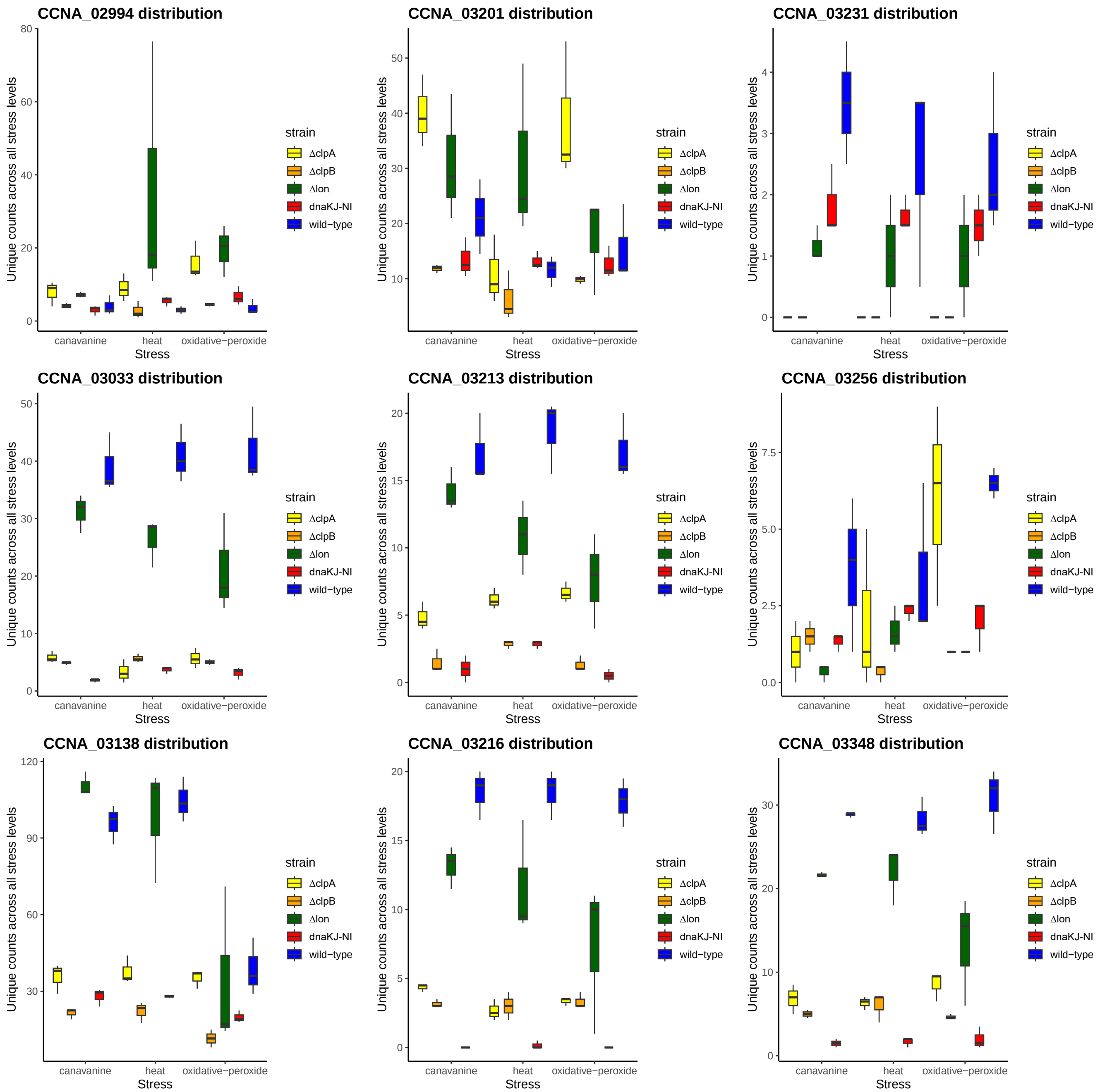


**Fig. S38. The unique count insertion distributions of fitness determinants** The Y-axis shows the unique count insertion distributions of genes that are predicted as fitness determinants by the model-Y knockoffs framework. This plot shows the distribution of CCNA_02994, CCNA_03201, CCNA_03231, CCNA_03033, CCNA_03213, CCNA_03256, CCNA_03138, CCNA_03216, CCNA_03348.

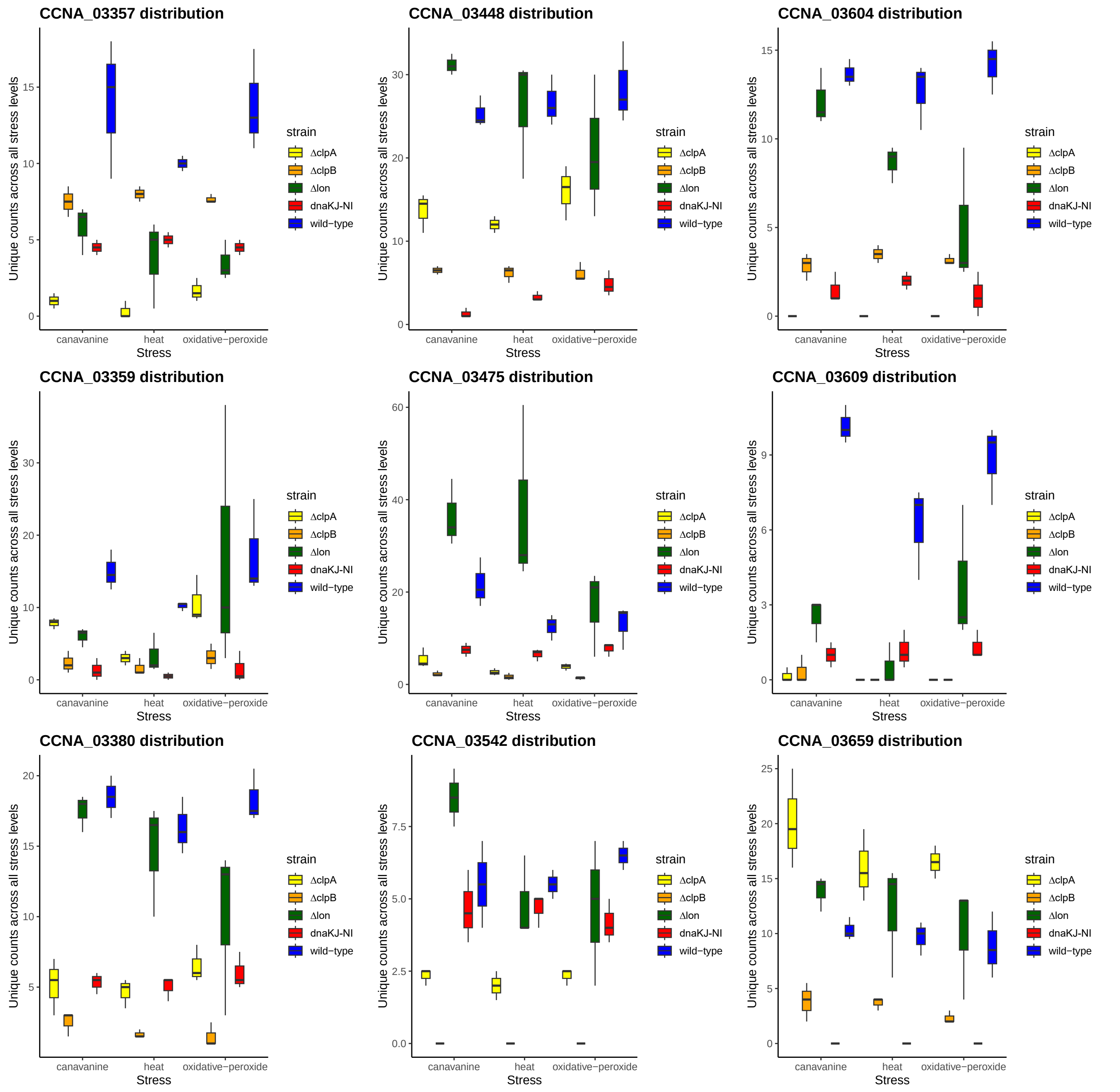


**Fig. S39. The unique count insertion distributions of fitness determinants** The Y-axis shows the unique count insertion distributions of genes that are predicted as fitness determinants by the model-Y knockoffs framework. This plot shows the distribution of CCNA_03357, CCNA_03448, CCNA_03604, CCNA_03359, CCNA_03475, CCNA_03609, CCNA_03380, CCNA_03542, CCNA_03659.

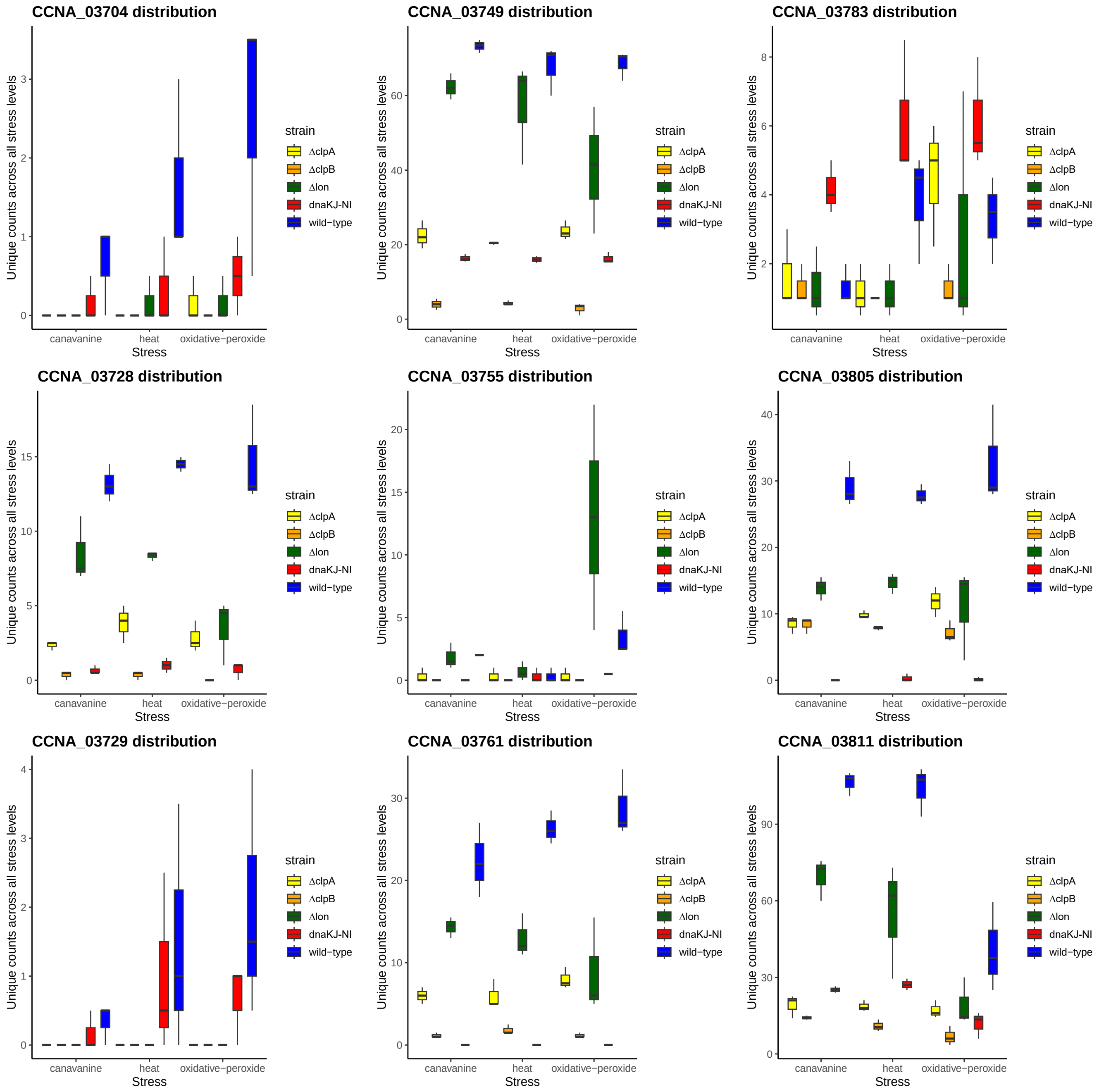


**Fig. S40. The unique count insertion distributions of fitness determinants** The Y-axis shows the unique count insertion distributions of genes that are predicted as fitness determinants by the model-Y knockoffs framework. This plot shows the distribution of CCNA_03704, CCNA_03749, CCNA_03783, CCNA_03728, CCNA_03755, CCNA_03805, CCNA_03729, CCNA_03761, CCNA_03811.

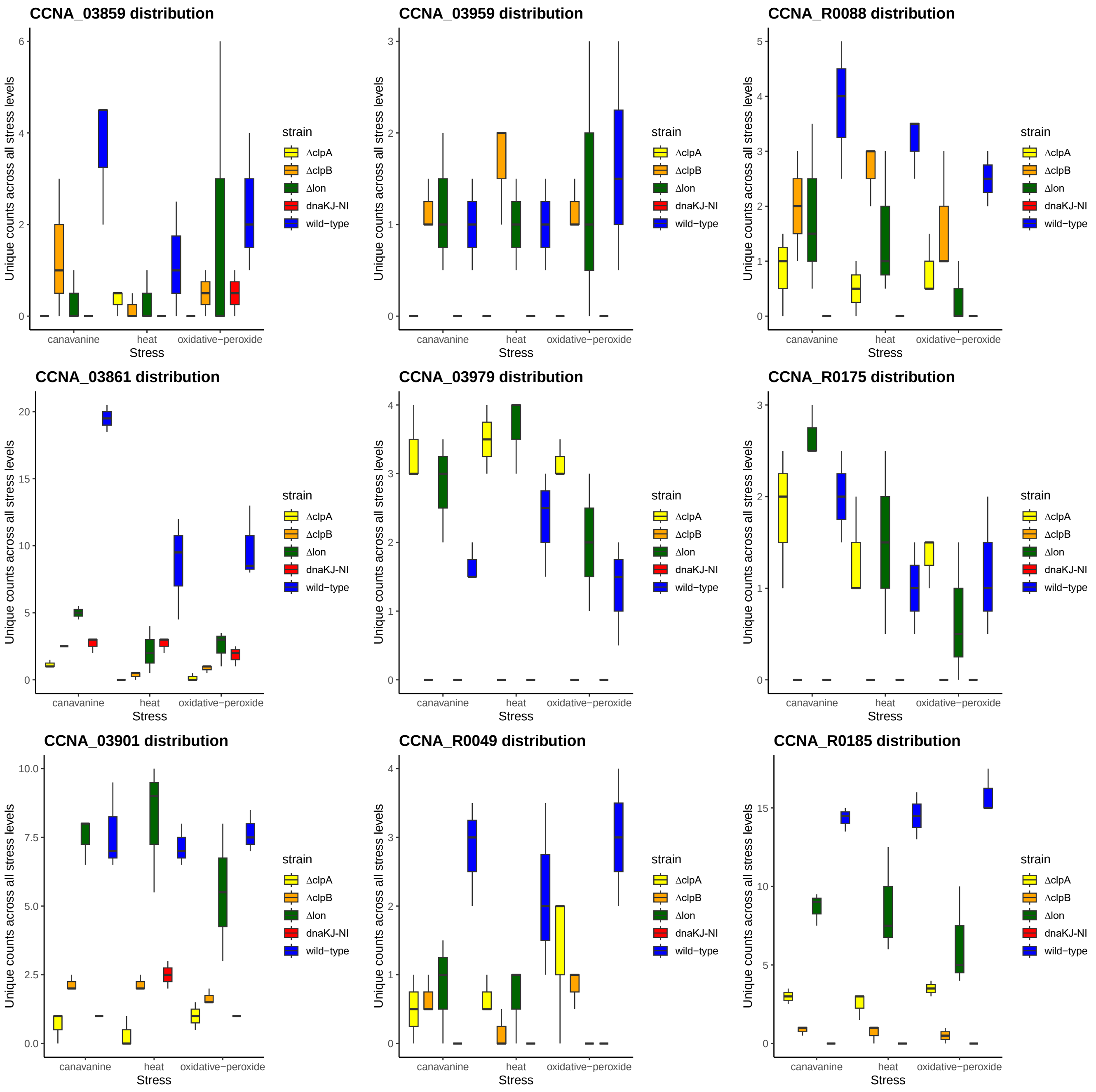


**Fig. S41. The unique count insertion distributions of fitness determinants** The Y-axis shows the unique count insertion distributions of genes that are predicted as fitness determinants by the model-Y knockoffs framework. This plot shows the distribution of CCNA_03859, CCNA_03959, CCNA_R0088, CCNA_03861, CCNA_03979, CCNA_R0175, CCNA_03901, CCNA_R0049, CCNA_R0185.

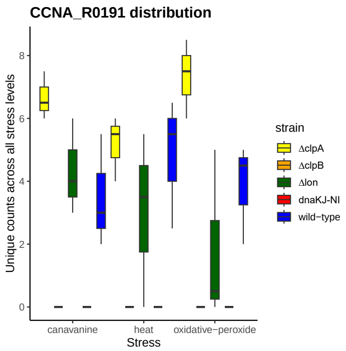


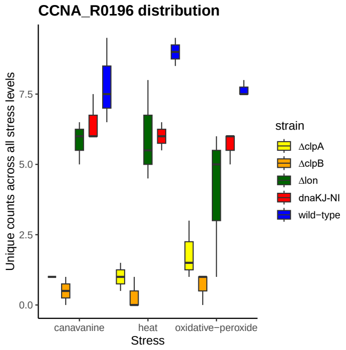


**Fig. S42. The unique count insertion distributions of fitness determinants** The Y-axis shows the unique count insertion distributions of genes that are predicted as fitness determinants by the model-Y knockoffs framework. This plot shows the distribution of CCNA_R0191, CCNA_R0196.

 

**Table S3. Protein names of shared fitness determinants among the strains. gene and protein names of the fitness predictors that are common, at least in two strains, are shown.**

| locus_tag | Protein names | gene names |
|---|---|---|
| CCNA_02154 | GCN5-related N-acetyltransferase | |
| CCNA_00922 | Chaperone protein ClpB | *clpB* |
| CCNA_03138 | Catalase-peroxidase (CP) (Peroxidase/catalase) | *katG* |
| CCNA_03811 | LysR-family transcriptional regulator | |
| CCNA_00001 | Putative pyruvate, phosphate dikinase regulatory protein | |
| CCNA_00706 | UrcA family protein | |
| CCNA_00240 | PTS system, IIA component | |
| CCNA_03729 | Probable transaldolase | *tal* |
| CCNA_00540 | Oxygen-dependent coproporphyrinogen-III oxidase | *hemF* |
| CCNA_02966 | Non-specific DNA-binding protein Dps/iron-binding ferritin-like antioxidant protein/ferroxidase | |
| CCNA_01213 | YjgP/YjgQ family membrane permease | |
| CCNA_03359 | Phosphoglycerate kinase (EC 2.7.2.3) | *pgk* |
| CCNA_01497 | ADP-L-glycero-D-manno-heptose-6-epimerase | *hldD* |
| CCNA_00823 | LuxR-like DNA-binding protein | |
| CCNA_02925 | Carbamoyl-phosphate synthase small chain | *carA* |
| CCNA_02572 | Adenylosuccinate lyase (ASL) | |
| CCNA_03861 | Pyridoxal phosphate homeostasis protein | |
| CCNA_00939 | MarR-family transcriptional regulator | |
| CCNA_00293 | Phosphate transport system permease protein PstA | *pstA* |

**Table S4. Gene ontology of shared fitness determinants among the strains. Gene ontology(GO) of the fitness predictors that are common, at least in two strains, are shown.**

| locus_tag | Gene ontology |
|---|---|
| CCNA_02154 | |
| CCNA_00922 | protein refolding [GO:0042026]; response to heat [GO:0009408] |
| CCNA_03138 | hydrogen peroxide catabolic process [GO:0042744]; response to oxidative stress [GO:0006979] |
| CCNA_03811 | |
| CCNA_00001 | protein dephosphorylation [GO:0006470]; protein phosphorylation [GO:0006468] |
| CCNA_00706 | |
| CCNA_00240 | phosphoenolpyruvate-dependent sugar phosphotransferase system [GO:0009401]; phosphorylation [GO:0016310] |
| CCNA_03729 | carbohydrate metabolic process [GO:0005975]; pentose-phosphate shunt [GO:0006098] |
| CCNA_00540 | protoporphyrinogen IX biosynthetic process [GO:0006782] |
| CCNA_02966 | |
| CCNA_01213 | |
| CCNA_03359 | glycolytic process [GO:0006096] |
| CCNA_01497 | ADP-L-glycero-beta-D-manno-heptose biosynthetic process [GO:0097171]; carbohydrate metabolic process [GO:0005975] |
| CCNA_00823 | regulation of DNA-templated transcription [GO:0006355] |
| CCNA_02925 | arginine biosynthetic process [GO:0006526]; glutamine metabolic process [GO:0006541] |
| CCNA_02572 | 'de novo' AMP biosynthetic process [GO:0044208]; 'de novo' IMP biosynthetic process [GO:0006189] |
| CCNA_03861 | |
| CCNA_00939 | |
| CCNA_00293 | phosphate ion transmembrane transport [GO:0035435] |

**Table S5. EMD (Earth Mover's Distance) based on both total and unique insertion counts for main Fig 4**

| Strain | Condition | EMD | Metric |
|---|---|---|---|
| wild-type | HT:L\|\|OS:L | 71.73507375 | Total_Counts |
| wild-type | HT:L\|\|OS:M | 88.9676351 | Total_Counts |
| wild-type | HT:L\|\|OS:H | 123.6415782 | Total_Counts |
| DCLPA | HT:L\|\|OS:L | 134.8418468 | Total_Counts |
| DCLPA | HT:L\|\|OS:M | 123.1473329 | Total_Counts |
| DCLPA | HT:L\|\|OS:H | 302.8568679 | Total_Counts |
| DCLPB | HT:L\|\|OS:L | 47.41432709 | Total_Counts |
| DCLPB | HT:L\|\|OS:M | 23.42034632 | Total_Counts |
| DCLPB | HT:L\|\|OS:H | 26.31701632 | Total_Counts |
| wild-type | HT:L\|\|OS:L | 3.466683818 | Unique_Counts |
| wild-type | HT:L\|\|OS:M | 3.128412525 | Unique_Counts |
| wild-type | HT:L\|\|OS:H | 6.94106338 | Unique_Counts |
| DCLPA | HT:L\|\|OS:L | 1.005183628 | Unique_Counts |
| DCLPA | HT:L\|\|OS:M | 1.160330579 | Unique_Counts |
| DCLPA | HT:L\|\|OS:H | 4.692307692 | Unique_Counts |
| DCLPB | HT:L\|\|OS:L | 0.313834649 | Unique_Counts |
| DCLPB | HT:L\|\|OS:M | 0.355353902 | Unique_Counts |
| DCLPB | HT:L\|\|OS:H | 0.429973475 | Unique_Counts |
